# Loss or Gain of Function? Effects of Ion Channel Mutations on Neuronal Firing Depend on the Cell Type

**DOI:** 10.1101/2023.01.16.524256

**Authors:** Nils A. Koch, Lukas Sonnenberg, Ulrike B.S. Hedrich, Stephan Lauxmann, Jan Benda

**Affiliations:** Institute for Neurobiology, Eberhard Karls Universitaet Tuebingen; Neurology and Epileptology, Hertie-Institute for Clinical Brain Research

## Abstract

Clinically relevant mutations to voltage-gated ion channels, called channelopathies, alter ion channel function, properties of ionic current and neuronal firing. The effects of ion channel mutations are routinely assessed and characterized as loss of function (LOF) or gain of function (GOF) at the level of ionic currents. Emerging personalized medicine approaches based on LOF/GOF characterization have limited therapeutic success. Potential reasons are that the translation from this binary characterization to neuronal firing especially when considering different neuronal cell types is currently not well understood. Here we investigate the impact of neuronal cell type on the firing outcome of ion channel mutations with simulations of a diverse collection of neuron models. We systematically analyzed the effects of changes in ion current properties on firing in different neuronal types. Additionally, we simulated the effects of mutations in the *KCNA1* gene encoding the K_V_1.1 potassium channel subtype associated with episodic ataxia type 1 (EA1). These simulations revealed that the outcome of a given change in ion channel properties on neuronal excitability is cell-type dependent. As a result, cell-type specific effects are vital to a full understanding of the effects of channelopathies on neuronal excitability and present an opportunity to further the efficacy and precision of personalized medicine approaches.

**Significance Statement:** Although the genetic nature of ion channel mutations as well as their effects on the biophysical properties of an ion channel are routinely assessed experimentally, determination of their role in altering neuronal firing is more difficult. In particular, cell-type dependency of ion channel mutations on firing has been observed experimentally, and should be accounted for. In this context, computational modelling bridges this gap and demonstrates that the cell type in which a mutation occurs is an important determinant in the effects of neuronal firing. As a result, classification of ion channel mutations as loss or gain of function is useful to describe the ionic current but should not be blindly extend to classification at the level of neuronal firing.

## Introduction

The properties and combinations of voltage-gated ion channels are vital in determining neuronal excitability (Bernard and Shevell, 2008; Carbone and Mori, 2020; Pospischil et al., 2008; Rutecki, 1992). However, ion channel function can be disturbed, for instance through genetic alterations, resulting in altered neuronal firing behavior (Carbone and Mori, 2020). In recent years, next generation sequencing has led to an increase in the discovery of clinically relevant ion channel mutations and has provided the basis for pathophysiological studies of genetic epilepsies, pain disorders, dyskinesias, intellectual disabilities, myotonias, and periodic paralyses (Bernard and Shevell, 2008; Carbone and Mori, 2020). Ongoing efforts of many research groups have contributed to the current understanding of underlying disease mechanism in channelopathies, however a complex pathophysiological landscape has emerged for many channelopathies and is likely a reason for limited therapeutic success with standard care.

Ion channel variants are frequently classified in heterologous expression systems as either a loss of function (LOF) or a gain of function (GOF) in the respective ionic current (Kim and Kang, 2021; Kullmann, 2002; Musto et al., 2020; Waxman, 2011). This LOF/GOF classification is often directly used to predict the effects on neuronal firing (Masnada et al., 2017; Niday and Tzingou-nis, 2018; Wei et al., 2017; Wolff et al., 2017), which in turn is important for understanding the pathophysiology of these disorders and for identification of potential therapeutic targets (Colasante et al., 2020; Orsini et al., 2018; Yang et al., 2018; Yu et al., 2006). Experimentally, the effects of channelopathies on neuronal firing are assessed using primary neuronal cultures (Liu et al., 2019; Scalmani et al., 2006; Smith et al., 2018) or *in vitro* recordings from slices of transgenic mouse lines (Habib et al., 2015; Hedrich et al., 2014; Lory et al., 2020; Mantegazza and Broccoli, 2019; Xie et al., 2010) but are restricted to limited number of neuronal types. Different neuron types differ in their composition of ionic currents (BRAIN Initiative Cell Census Network, 2021; Cadwell et al., 2016; Scala et al., 2021; Yao et al., 2021) and therefore likely respond differently to changes in the properties of a single ionic current. Expression level of an affected gene (Layer et al., 2021) and relative amplitudes of ionic currents (Barreiro et al., 2012; Golowasch et al., 2002; Kispersky et al., 2012; Pospischil et al., 2008; Rutecki, 1992) indeed dramatically influence the firing behavior and dynamics of neurons. Mutations in different sodium channel genes have been experimentally shown to affect firing in a cell-type specific manner based on differences in expression levels of the affected gene (Layer et al., 2021), but also on other cell-type specific mechanisms (Hedrich et al., 2014; Makinson et al., 2016).

Cell-type specificity is likely vital for successful precision medicine treatment approaches. For example, Dravet syndrome was identified as the consquence of LOF mutations in *SCN1A* (Claes et al., 2001; Fujiwara et al., 2003; Ohmori et al., 2002), however limited success in the treatment of Dravet syndrome persisted (Claes et al., 2001; Oguni et al., 2001) in part due to lack of understanding that inhibitory interneurons and not pyramidal neurons had altered excitability as a result of LOF *SCN1A* mutations (Colasante et al., 2020; Yu et al., 2006).

Taken together, these examples demonstrate the need to study the effects of ion channel mutations in many different cell types — a daunting if not impossible experimental challenge. In the context of this diversity, simulations of conductance-based neuronal models are a powerful tool bridging the gap between altered ionic currents and firing in a systematic and efficient way. Furthermore, simlutions allow to predict the potential effects of drugs needed to alleviate the pathophysiology of the respective mutation (Bayraktar et al., In Press; Johannesen et al., 2021; Lauxmann et al., 2021).

In this study, we therefore investigated how the outcome of ionic current kinetic changes on firing depend on neuronal cell type by (1) characterizing firing responses with two measures, (2) simulating the response of a repertoire of different neuronal models to changes in single current parameters as well as (3) to more complex changes in this case as they were observed for specific *KCNA1* mutations that are associated with episodic ataxia type 1 (Browne et al., 1995, 1994; Lauxmann et al., 2021).

## Materials and Methods

All modelling and simulation was done in parallel with custom written Python 3.8 software, run on a Cent-OS 7 server with an Intel(R) Xeon (R) E5-2630 v2 CPU.

### Different Cell Models

A group of neuronal models representing the major classes of cortical and thalamic neurons including regular spiking pyramidal (RS pyramidal; model D), regular spiking inhibitory (RS inhibitory; model B), and fast spiking (FS; model C) cells were used (Pospischil et al., 2008). Additionally, a K_V_1.1 current (I_K_V_1.1_; Ranjan et al. 2019) was added to each of these models (RS pyramidal +K_V_1.1; model H, RS inhibitory +K_V_1.1; model E, and FS +K_V_1.1; model G respectively). A cerebellar stellate cell model from Alexander et al. (2019) is used (Cb stellate; model A) in this study. This cell model was also extended by a K_V_1.1 current (Ranjan et al., 2019), either in addition to the A-type potassium current (Cb stellate +K_V_1.1; model F) or by replacing the A-type potassium current (Cb stellate ΔK_V_1.1; model J). A subthalamic nucleus (STN; model L) neuron model as described by Otsuka et al. (2004) was also used. The STN cell model (model L) was additionally extended by a K_V_1.1 current (Ranjan et al., 2019), either in addition to the A-type potassium current (STN +K_V_1.1; model I) or by replacing the A-type potassium current (STN ΔK_V_1.1; model K). Model letter naming corresponds to panel lettering in Figure 1. The properties and maximal conductances of each model are detailed in Table 1 and the gating properties are unaltered from the original Cb stellate (model A) and STN (model L) models (Alexander et al., 2019; Otsuka et al., 2004). For enabling the comparison of models with the typically reported electrophysiological data fitting reported and for ease of further gating curve manipulations, a modified Boltzmann function

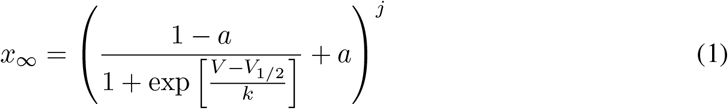

with slope *k*, voltage for half-maximal activation or inactivation (*V*_1/2_), exponent *j*, and persistent current 0 ≤ *a* ≤ 1 were fitted to the original formulism for RS pyramidal (model D), RS inhibitory (model B) and FS (model C) models from Pospischil et al. (2008). The properties of I_K_V_1.1_ were fitted to the mean wild type biophysical parameters of K_V_1.1 described in Lauxmann et al. (2021). Each of the original single-compartment models used here can reproduce physiological firing behavior of the neurons they represent (Figure 1; Alexander et al. 2019; Otsuka et al. 2004; Pospischil et al. 2008) and capture key aspects of the dynamics of these cell types.

**Figure 1:**
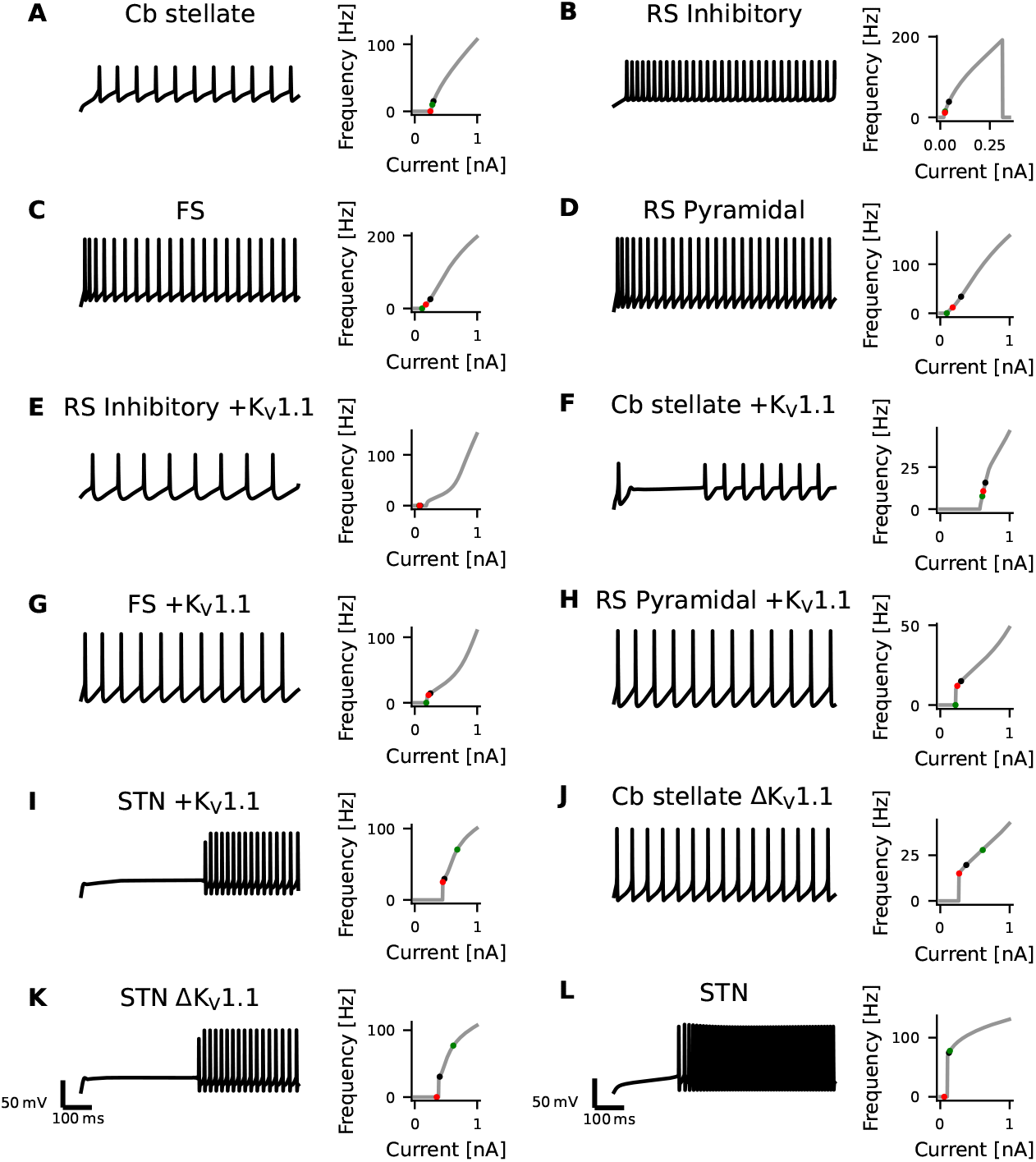
Diversity in Neuronal Model Firing. Spike trains (left), frequency-current (fI) curves (right) for Cb stellate (A), RS inhibitory (B), FS (C), RS pyramidal (D), RS inhibitory +K_V_1.1 (E), Cb stellate +K_V_1.1 (F), FS +K_V_1.1 (G), RS pyramidal +K_V_1.1 (H), STN +K_V_1.1 (I), Cb stellate ΔK_V_1.1 (J), STN ΔK_V_1.1 (K), and STN (L) neuron models. Models are sorted qualitatively based on their fI curves. Black markers on the fI curves indicate the current step at which the spike train occurs. The green marker indicates the current at which firing begins in response to an ascending current ramp, whereas the red marker indicates the current at which firing ceases in response to a descending current ramp (see Figure 1-1).

**Table 1:** Cell properties and conductances of regular spiking pyramidal neuron (RS Pyramidal; model D), regular spiking inhibitory neuron (RS Inhibitory; model B), fast spiking neuron (FS; model C) each with additional I_K_V_1.1_ (RS Pyramidal +K_V_1.1; model H, RS Inhibitory +K_V_1.1; model E, FS +K_V_1.1; model G respectively), cerebellar stellate cell (Cb Stellate; model A), with additional I_K_V_1.1_ (Cb Stellate +K_V_1.1; model F) and with I_K_V_1.1_ replacement of I_A_ (Cb Stellate ΔK_V_1.1; model J), and subthalamic nucleus neuron (STN; model L), with additional I_K_V_1.1_ (STN +K_V_1.1; model I) and with I_K_V_1.1_ replacement of I_A_ (STN K_V_1.1; model K) models. All conductances are given in mS/cm^2^. Capacitances (*C_m_*) and *τ_max_,_M_* are given in pF and ms respectively.

| | RS<br>Pyra-<br>midal<br>(+K<sub>V</sub>1.1) | RS<br>Inhib-<br>itory<br>(+K<sub>V</sub>1.1) | FS<br>(+K<sub>V</sub>1.1) | Cb<br>Stellate | Cb<br>Stellate<br>+K<sub>V</sub>1.1 | Cb<br>Stellate<br>ΔK<sub>V</sub>1.1 | STN | STN<br>+K<sub>V</sub>1.1 | STN<br>ΔK<sub>V</sub>1.1 |
| --- | --- | --- | --- | --- | --- | --- | --- | --- | --- |
| Model | D (H) | B (E) | C (G) | A | F | J | L | I | K |
| $g_{Na}$ | 56 | 10 | 58 | 3.4 | 3.4 | 3.4 | 49 | 49 | 49 |
| $g_K$ | 6 (5.4) | 2.1 (1.89) | 3.9 (3.51) | 9.0556 | 8.15 | 9.0556 | 57 | 56.43 | 57 |
| $g_{K_V1.1}$ | — (0.6) | — (0.21) | — (0.39) | — | 0.90556 | 1.50159 | — | 0.57 | 0.5 |
| $g_A$ | — | — | — | 15.0159 | 15.0159 | — | 5 | 5 | — |
| $g_M$ | 0.075 | 0.0098 | 0.075 | — | — | — | — | — | — |
| $g_L$ | — | — | — | — | — | — | 5 | 5 | 5 |
| $g_T$ | — | — | — | 0.45045 | 0.45045 | 0.45045 | 5 | 5 | 5 |
| $g_{Ca,K}$ | — | — | — | — | — | — | 1 | 1 | 1 |
| $g_{Leak}$ | 0.0205 | 0.0205 | 0.038 | 0.07407 | 0.07407 | 0.07407 | 0.035 | 0.035 | 0.035 |
| $\tau_{max,M}$ | 608 | 934 | 502 | — | — | — | — | — | — |
| $C_m$ | 118.44 | 119.99 | 101.71 | 177.83 | 177.83 | 177.83 | 118.44 | 118.44 | 118.44 |

**Table 2:** For comparability to typical electrophysiological data fitting reported and for ease of further gating curve manipulations, a sigmoid function (Eqn.1) with slope *k*, voltage for half-maximal activation or inactivation (*V*_1/2_), exponent *j*, and persistent current 0 ≤ *a* ≤ 1 were fitted for the models originating from Pospischil et al. (2008) (models B, C, D, E, G, H) where *α_x_* and *β_x_* are used. Gating parameters for I_K_V_1.1_ are taken from Ranjan et al. (2019) and fit to mean wild type parameters in Lauxmann et al. (2021). Model gating parameters not listed are taken directly from source publication.

| | Gating | $V_{1/2}$ [mV] | $k$ | $j$ | $a$ |
| --- | --- | --- | --- | --- | --- |
| Models<br>B, C, D, E, G, H | $I_{Na}$ activation | -34.33054521 | -8.21450277 | 1.42295686 | — |
| | $I_{Na}$ inactivation | -34.51951036 | 4.04059373 | 1 | 0.05 |
| | $I_{Kd}$ activation | -63.76096946 | -13.83488194 | 7.35347425 | — |
| | $I_L$ activation | -39.03684525 | -5.57756176 | 2.25190197 | — |
| | $I_L$ inactivation | -57.37 | 20.98 | 1 | — |
| | $I_M$ activation | -45 | -9.9998807337 | 1 | — |
| $I_{Kv1.1}$ | $I_{Kv1.1}$ activation | -30.01851852 | -7.73333333 | 1 | — |
| | $I_{Kv1.1}$ Inactivation | -46.85851852 | 7.67266667 | 1 | 0.245 |

**Table 3:** Statistical Table. Descriptive statistics including non-parametric Kendall *τ* rank correlations are used. Statistical hypothesis tests are not used.

|  | Data Structure | Type of test | Power |
| --- | --- | --- | --- |
| a | Non-normal distribution | Kendal $\tau$ rank correlation | — |

### Firing Frequency Analysis

The membrane responses to 200 equidistant two second long current steps were simulated using the forward-Euler method with a Δt = 0.01 ms from steady state. Current steps ranged from 0 to 1 nA (step size 5 pA) for all models except for the RS inhibitory neuron models where a range of 0 to 0.35 nA (step size 1.75 pA) was used to ensure repetitive firing across the range of input currents. For each current step, action potentials were detected as peaks with at least 50mV prominence, or the relative height above the lowest contour line encircling it, and a minimum interspike interval of 1 ms. The interspike interval was computed and used to determine the instantaneous firing frequencies elicited by the current step.

To ensure accurate firing frequencies at low firing rates and reduced spike sampling bias, steady-state firing was defined as the mean firing frequency in a 500 ms window in the last second of the current steps starting at the inital action potential in this last second. Firing characterization was performed in the last second of current steps to ensure steady-state firing is captured and adaptation processes are neglected in our analysis. Alteration in current magnitudes can have different effects on rheobase and the initial slope of the fI curve (Kispersky et al., 2012). For this reason, we quantified neuronal firing using the rheobase as well as the area under the curve (AUC) of the initial portion of the fI curve as a measure of the initial slope of the fI curve Figure 2 A.

**Figure 2:**
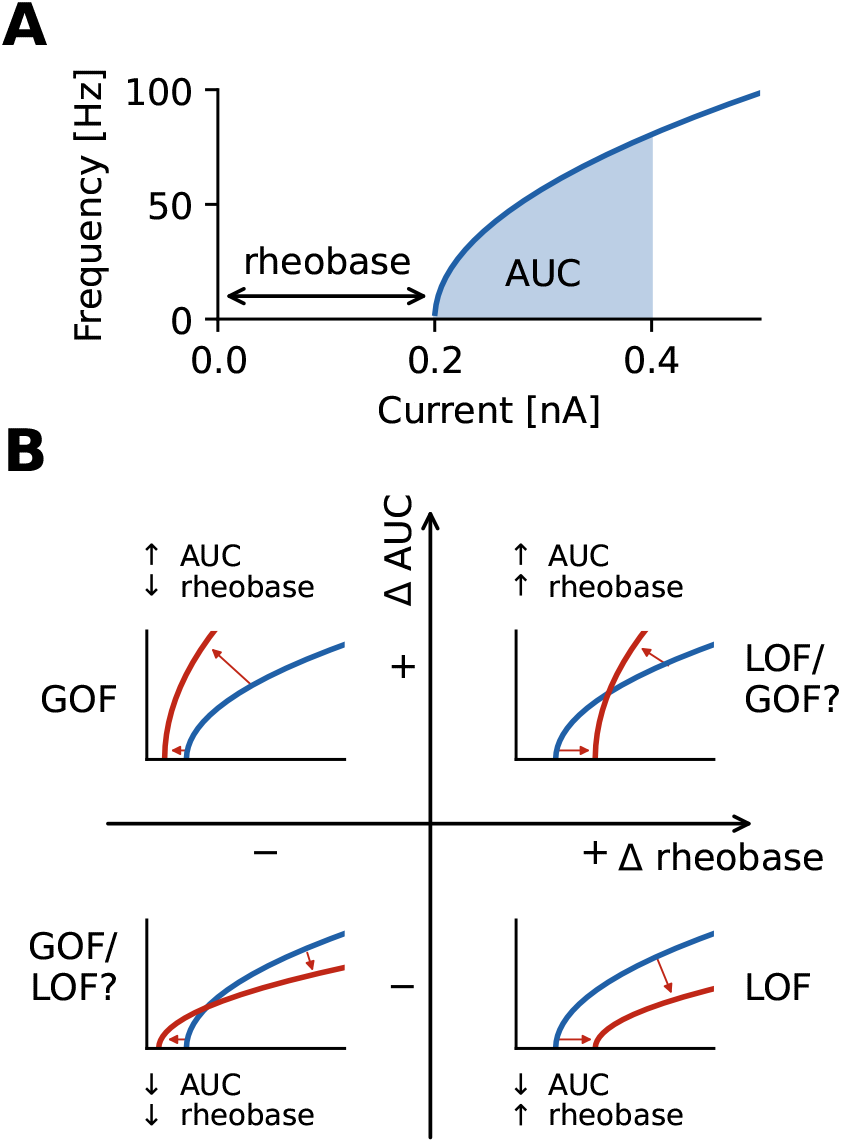
Characterization of firing with AUC and rheobase. (A) The area under the curve (AUC) of the repetitive firing frequency-current (fI) curve. (B) Changes in firing as characterized by ΔAUC and Δrheobase occupy four quadrants separated by no changes in AUC and rheobase. Representative schematic fI curves in red with respect to a reference (or wild type) fI curve (blue) depict the general changes associated with each quadrant.

The smallest current at which steady state firing occured was identified and the current step interval preceding the occurrence of steady state firing was simulated at higher resolution (100 current steps) to determine the current at which steady state firing began. Firing was simulated with 100 current steps from this current upwards for 1/5 of the overall current range. Over this range a fI curve was constructed and the integral, or area under the curve (AUC), of the fI curve over this interval was computed with the composite trapezoidal rule and used as a measure of firing rate independent from rheobase.

To obtain the rheobase at a higher current resolution than the fI curve, the current step interval preceding the occurrence of action potentials was explored at higher resolution with 100 current steps spanning the interval (step sizes of 0.05 pA and 0.0175 pA, respectively). Membrane responses to these current steps were then analyzed for action potentials and the rheobase was considered the lowest current step for which an action potential was elicited.

All models exhibited tonic steady-state firing with default parameters. In limited instances, variations of parameters elicited periodic bursting, however these instances were excluded from further analysis.

### Sensitivity Analysis and Comparison of Models

Properties of ionic currents common to all models (I_Na_, I_K_, I_A_/I_K_V_1.1_, and I_Leak_) were systematically altered in a one-factor-at-a-time sensitivity analysis for all models. The gating curves for each current were shifted (Δ*V*_1/2_) from −10 to 10mV in increments of 1mV. The voltage dependence of the time constant associated with the shifted gating curve was correspondingly shifted. The slope (*k*) of the gating curves were altered from half to twice the initial slope. Similarly, the maximal current conductance (*g*) was also scaled from half to twice the initial value. For both slope and conductance alterations, alterations consisted of 21 steps spaced equally on a log_2_ scale. We neglected the variation of time constants for the practical reason that estimation and assessment of time constants and changes to them is not straightforward (Clerx et al., 2019; Whittaker et al., 2020).

### Model Comparison

Changes in rheobase (Δrheobase) were calculated in relation to the original model rheobase. The contrast of each AUC value (*AUC_i_*) was computed in comparison to the AUC of the unaltered wild type model (*AUC_wt_*)

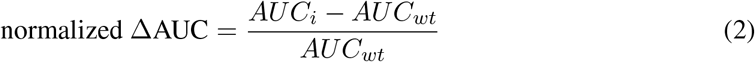

To assess whether the effects of a given alteration on normalized ΔAUC or Δrheobase were robust across models, the correlation between normalized ΔAUC or Δrheobase and the magnitude of the alteration of a current property was computed for each alteration in each model and compared across alteration types. The Kendall’s *τ* coefficient, a non-parametric rank correlation, is used to describe the relationship between the magnitude of the alteration and AUC or rheobase values. A Kendall *τ* value of −1 or 1 is indicative of monotonically decreasing and increasing relationships respectively.

### *KCNA1* Mutations

Known episodic ataxia type 1 associated *KCNA1* mutations and their electrophysiological charac-terization have been reviewed in Lauxmann et al. (2021). The mutation-induced changes in I_K_V_1.1_ amplitude and activation slope (*k*) were normalized to wild type measurements and changes in activation *V*_1/2_ were used relative to wild type measurements. Although initially described to lack fast activation, K_V_1.1 displays prominent inactivation at physiologically relevant temperatures (Ranjan et al., 2019). The effects of a mutation were also applied to I_A_ when present as both potassium currents display inactivation. In all cases, the mutation effects were applied to half of the K_V_1.1 or I_A_ under the assumption that the heterozygous mutation results in 50% of channels carrying the mutation. Frequency-current curves for each mutation in each model were obtained through simulation and used to characterize firing behavior as described above. For each model the differences in mutation AUC to wild type AUC were normalized by wild type AUC (normalized ΔAUC) and mutation rheobases were compared to wild type rheobase values (Δrheobase). Pairwise Kendall rank correlations (Kendall *τ*) were used to compare the correlation in the effects of K_V_1.1 mutations on AUC and rheobase between models.

### Code Accessibility

The code/software described in the paper is freely available online at https://github.com/nkoch1/LOFGOF2023. The code is available as Extended Data 1.

## Results

To examine the role of cell-type specific ionic current environments on the impact of altered ion currents properties on firing behavior: (1) firing responses were characterized with rheobase and ΔAUC, (2) a set of neuronal models was used and properties of channels common across models were altered systematically one at a time, and (3) the effects of a set of episodic ataxia type 1 associated *KCNA1* mutations on firing was then examined across different neuronal models with different ionic current environments.

### Variety of model neurons

Neuronal firing is heterogenous across the CNS and a set of neuronal models with heterogenous firing due to different ionic currents is desirable to reflect this heterogeneity. The set of singlecompartment, conductance-based neuronal models used here has considerable diversity as evident in the variability seen across neuronal models both in spike trains and their fI curves (Figure 1). The models chosen for this study all fire tonically and do not exhibit bursting (see methods for details and naming of the models). Models are qualitatively sorted based on their firing curves and labeled model A through L accordingly. Some models, such as models A and B, display type I firing, whereas others such as models J and L exhibit type II firing. Type I firing is characterized by continuous fI curves (i.e. firing rate increases from 0 in a continuous fashion) whereas type II firing is characterized by a discontinuity in the fI curve (i.e. a jump occurs from no firing to firing at a certain frequency; Ermentrout 1996; Rinzel and Ermentrout 1989). The other models used here lie on a continuum between these prototypical firing classifications. Most neuronal models exhibit hysteresis with ascending and descending ramps eliciting spikes at different current thres-holds. However, the models I, J, and K have large hysteresis (Figures 1 and 1-1). Different types of underlying current dynamics are known to generate these different firing types and hysteresis (Ermentrout, 1996; Ermentrout and Chow, 2002; Izhikevich, 2006).

### Characterization of Neuronal Firing Properties

Neuronal firing is a complex phenomenon, and a quantification of firing properties is required for comparisons across cell types and between different conditions. Here we focus on two aspects of firing: rheobase, the smallest injected current at which the cell fires an action potential, and the shape of the frequency-current (fI) curve as quantified by the area under the curve (AUC) for a fixed range of input currents above rheobase (Figure 2 A). The characterization of the firing properties of a neuron by using rheobase and AUC allows to characterize both a neuron’s excitability in the sub-threshold regime (rheobase) and periodic firing in the super-threshold regime (AUC) by two independent measures. Note that AUC is essentially quantifying the slope of a neuron’s fI curve.

Using these two measures we quantified the effects a changed property of an ionic current has on neural firing by the differences in both rheobase, Δrheobase, and in AUC, ΔAUC, relative to the wild type neuron. ΔAUC is in addition normalized to the AUC of the wild type neuron, see Eq. (2). Each fI curve resulting from an altered ionic current is a point in a two-dimensional coordinate system spanned by Δrheobase and normalized ΔAUC (Figure 2 B). An fI curve similar to the one of the wild type neuron is marked by a point close to the origin. In the upper left quadrant, fI curves become steeper (positive difference of AUC values: +ΔAUC) and are shifted to lower rheobases (negative difference of rheobases: −Δrheobase), unambigously indicating an increased firing that clearly might be classified as a gain of function (GOF) of neuronal firing. The opposite happens in the bottom right quadrant where the slope of fI curves decreases (−ΔAUC) and the rheobase is shifted to higher currents (+Δrheobase), indicating a decreased, loss of function (LOF) firing. In the lower left (−ΔAUC and −Δrheobase) and upper right (+ΔAUC and +Δrheobase) quadrants, the effects on firing are less clear-cut, because the changes in rheobase and AUC have opposite effects on neuronal firing. Changes in a neuron’s fI curves in these two quadrants cannot uniquely be described as a gain or loss of excitability.

### Sensitivity Analysis

Sensitivity analyses are used to understand how input model parameters contribute to determining the output of a model (Saltelli, 2002). In other words, sensitivity analyses are used to understand how sensitive the output of a model is to a change in input or model parameters. One-factor-a-time sensitivity analyses involve altering one parameter at a time and assessing the impact of this parameter on the output. This approach enables the comparison of given alterations in parameters of ionic currents across models.

For example, when shifting the half activation voltage *V*_1/2_ of the delayed rectifier potassium current in the model G to more depolarized values, then the rheobase of the resulting fI curves shifted to lower currents −Δrheobase, making the neuron more sensitive to weak inputs, but at the same time the slope of the fI curves was reduced (−normalized ΔAUC), which resulted in a reduced firing rate (Figure 3 A). As a result the effect of a depolarizing shift in the delayed rectifier potassium current half activation *V*_1/2_ in model C is in the bottom left quadrant of Figure 2 B and characterization as LOF or GOF in excitability is not possible. Plotting the corresponding changes in AUC against the change in half activation potential *V*_1/2_ results in a monotonically falling curve (thick orange line in Figure 3 B). For each of the many models we got a different relation between the changes in AUC and the shifts in half maximal potential *V*_1/2_ (thin lines in Figure 3 B). To further summarize these different dependencies of the various models we characterized each of these curves by a single number, the Kendall *τ* correlation coefficient. A monotonically increasing curve resulted in a Kendall *τ* close to +1 a monotonously decreasing curve in Kendall *τ* ≈ − 1, and a non-monotonous, non-linear relation in Kendall *τ* close to zero (compare lines in Figure 3 B with dots in black box in panel C).

**Figure 3:**
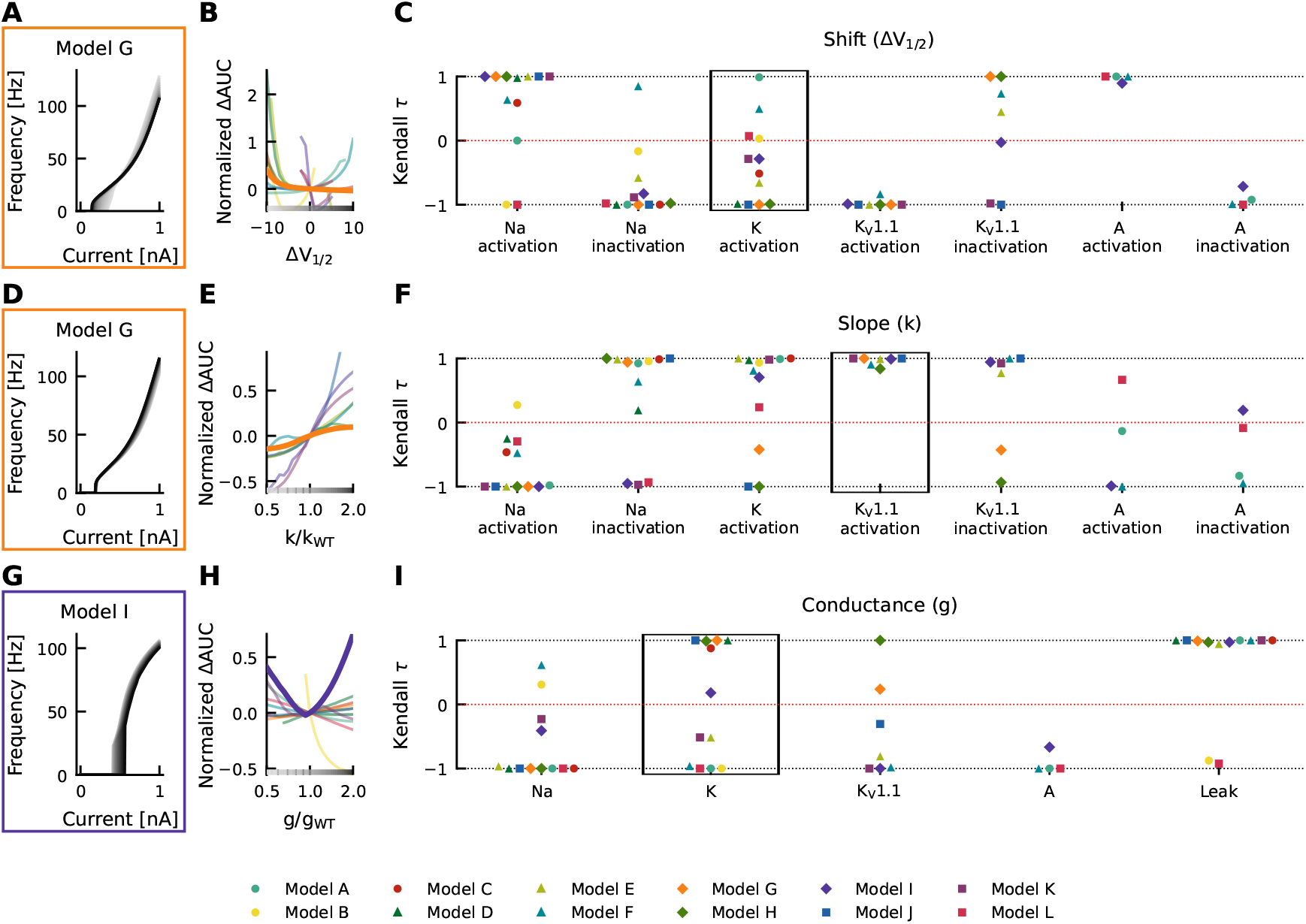
Effects of altered channel kinetics on AUC in various neuron models. The fI curves corresponding to shifts in model G delayed rectifier K half activation *V*_1/2_ (A), changes K_V_1.1 activation slope factor k in model G (D), and changes in maximal conductance of delayed rectifier K current in the model I (G) are shown. The fI curves from the smallest (grey) to the largest (black) alterations are seen for (A,D, and G) in accordance to the greyscale of the x-axis in B, E, and H. The normalized ΔAUC of fI curves is plotted against delayed rectifier K half activation potential (Δ*V*_1/2_; D), K_V_1.1 activation slope factor *k* (k/k_*WT*_; E) and maximal conductance *g* of the delayed rectifier K current (g/g_*WT*_; H) for all models (thin lines) with relationships from the fI curve examples (A, D, G respectively) highlighted by thick lines with colors corresponding to the box highlighting each set of fI curves. The Kendall rank correlation (Kendall *τ*) coefficients between shifts in half maximal potential *V*_1/2_ and normalized ΔAUC (C), slope factor k and normalized ΔAUC (F) as well as maximal current conductances and normalized ΔAUC (I) for each model and current property is computed. The relationships between Δ*V*_1/2_, k/k_*WT*_, and g/g_*WT*_ and normalized ΔAUC for the Kendall rank correlations highlighted in the black boxes are depicted in (B), (E) and (H) respectively.

Changes in gating half activation potential *V*_1/2_ and slope factor *k* as well as the maximum conductance *g* affected the AUC (Figure 3), but how exactly the AUC was affected usually depended on the specific neuronal model. Increasing the slope factor of the K_V_1.1 activation curve for example increased the AUC in all models (Kendall *τ* ≈ +1), but with different slopes (Figure 3 D,E,F). Similar consistent positive correlations could be found for shifts in A-current activation *V*_1/2_. Changes in K_V_1.1 half activation *V*_1/2_ and in maximal A-current conductance resulted in negative correlations with the AUC in all models (Kendall *τ* ≈ − 1).

Qualitative differences could be found, for example, when increasing the maximal conductance of the delayed rectifier (Figure 3 G,H,I). In some model neurons this increased AUC (Kendall *τ* ≈ +1), whereas in others AUC was decreased (Kendall *τ* ≈ −1). In model I, AUC depended in a non-linear way on the maximal conductance of the delayed rectifier, resulting in a Kendall *τ* close to zero. Even more dramatic qualitative differences between models resulted from shifts of the activation curve of the delayed rectifier, as discussed already above (Figure 3 A,B,C). Some model neurons did almost not depend on changes in K-current half activation *V*_1/2_ or showed strong non-linear dependencies, both resulting in Kendall *τ* close to zero. Many model neurons showed strongly negative correlations, and a few displayed positive correlations with shifting the activation curve of the delayed rectifier.

Changes in gating half activation potential *V*_1/2_ and slope factor *k* as well as the maximum conductance *g* affected rheobase (Figure 4). However, in contrast to AUC, qualitatively consistent effects on rheobase across models could be observed. An increasing of the maximal conductance of the leak current in the model A increased the rheobase (Figure 4 G). When these changes were plotted against the change in maximal conductance a monotonically increasing relationship was evident (thick teal line in Figure 4 H). This monotonically increasing relationship was evident in all models (Kendall *τ* ≈ +1), but with different slopes (thin lines in Figure 4 H). Similarly, positive correlations were consistently found across models for maximal conductances of delayed rectifier K, K_V_1.1, and A type currents, whereas the maximal conductance of the sodium current was consistently associated with negative correlations (Kendall *τ* ≈ − 1; Figure 4 I), i.e. rheobase decreased with increasing maximum conductance in all models.

**Figure 4:**
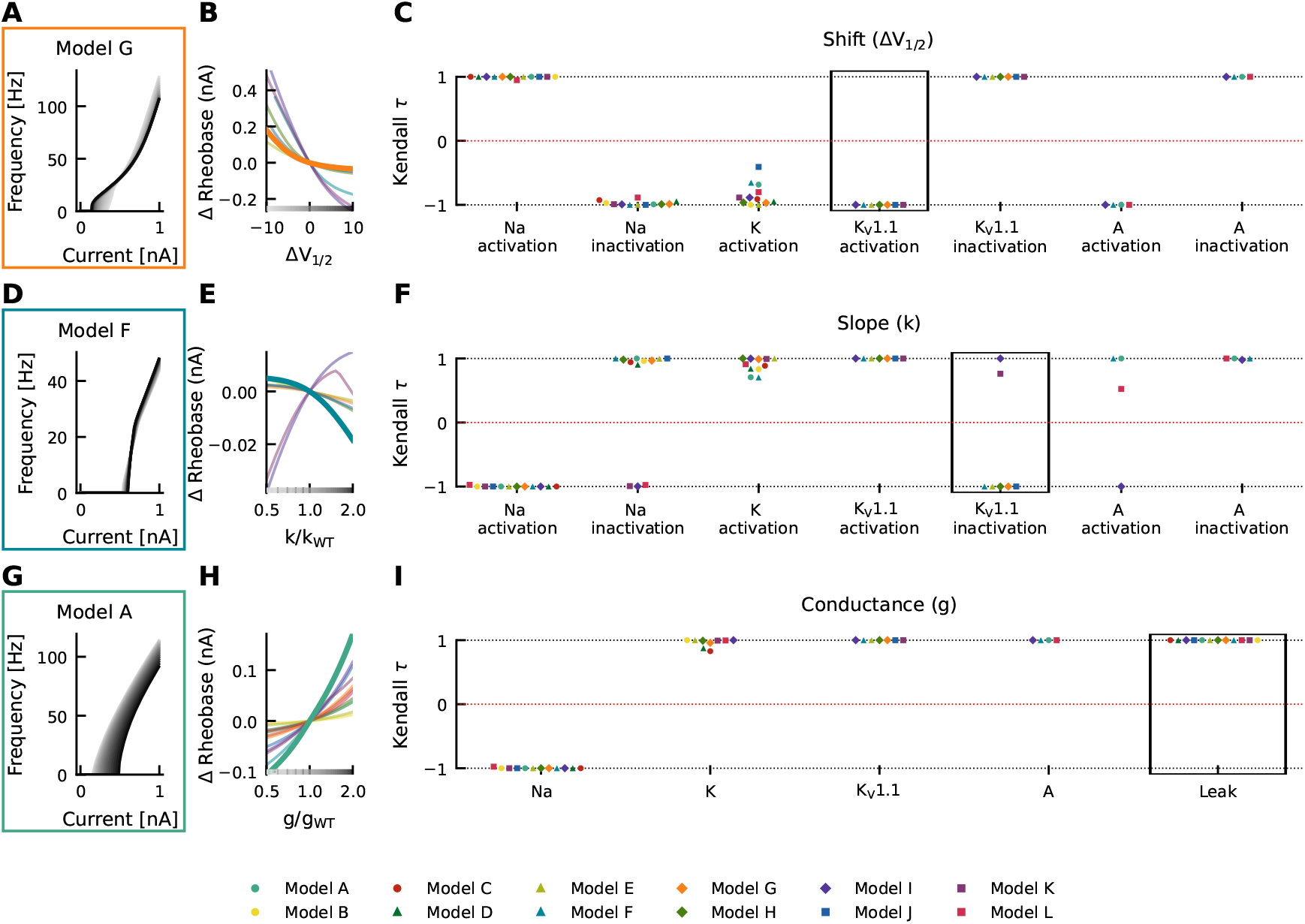
Effects of altered channel kinetics on rheobase. The fI curves corresponding to shifts in model G K_V_1.1 activation *V*_1/2_ (A), changes K_V_1.1 inactivation slope factor *k* in model F (D), and changes in maximal conductance of the leak current in model A (G) are shown. The fI curves from the smallest (grey) to the largest (black) alterations are seen for (A,D, and G) in accordance to the greyscale of the x-axis in B, E, and H. The Δrheobase of fI curves is plotted against K_V_1.1 half activation potential (Δ*V*_1/2_; B), K_V_1.1 inactivation slope factor *k* (k/k_*WT*_; E) and maximal conductance *g* of the leak current (g/g_*WT*_; H) for all models (thin lines) with relationships from the fI curve examples (A, D, G respectively) highlighted by thick lines with colors corresponding to the box highlighting each set of fI curves. The Kendall rank correlation (Kendall *τ*) coefficients between shifts in half maximal potential *V*_1/2_ and Δrheobase (C), slope factor k and Δrheobase (F) as well as maximal current conductances and Δrheobase (I) for each model and current property is computed. The relationships between Δ*V*_1/2_, k/k_*WT*_, and g/g_*WT*_ and Δrheobase for the Kendall rank correlations highlighted in the black boxes are depicted in (B), (E) and (H) respectively.

Although changes in half maximal potential *V*_1/2_ and slope factor *k* generally correlated with rheobase similarly across models there were some exceptions. Rheobase was affected with both with positive and negative correlations in different models as a result of changing slope factor of Na^+^-current inactivation (positive: models A–H and J; negative: models I, K and L), K_V_1.1-current inactivation (positive: models I and K; negative: models E–G, J, H), and A-current activation (positive: models A, F and L; negative: model I; Figure 4 F). Departures from monotonic relationships also occurred in some models as a result of K^+^-current activation *V*_1/2_ (e.g. model J) and slope factor *k* (models F and G), K_V_1.1-current inactivation slope factor *k* (model K), and A-current activation slope factor *k* (model L). Thus, identical changes in current gating properties such as the half maximal potential V_1_/_2_ or slope factor k can have differing effects on firing depending on the model in which they occur.

### *KCNA1* Mutations

Mutations in *KCNA1* are associated with episodic ataxia type 1 (EA1) and have been characterized biophysically (as reviewed by Lauxmann et al. (2021)). Here they were used as a test case in the effects of various ionic current environments on neuronal firing and on the outcomes of channelopathies. The changes in AUC and rheobase from wild type values for reported EA1 associated *KCNA1* mutations were heterogeneous across models containing K_V_1.1, but generally showed decreases in rheobase (Figure 5A–I). Pairwise non-parametric Kendall *τ* rank correlations between the simulated effects of these K_V_1.1 mutations on rheobase were highly correlated across models (Figure 5J) indicating that EA1 associated *KCNA1* mutations generally decrease rheobase across diverse cell-types. However, the effects of the K_V_1.1 mutations on AUC were more heterogenous as refIected by both weak and strong positive and negative pairwise correlations between models (Figure 5K), suggesting that the effects of ion-channel variant on super-threshold neuronal firing depend both quantitatively and qualitatively on the specific composition of ionic currents in a given neuron.

**Figure 5:**
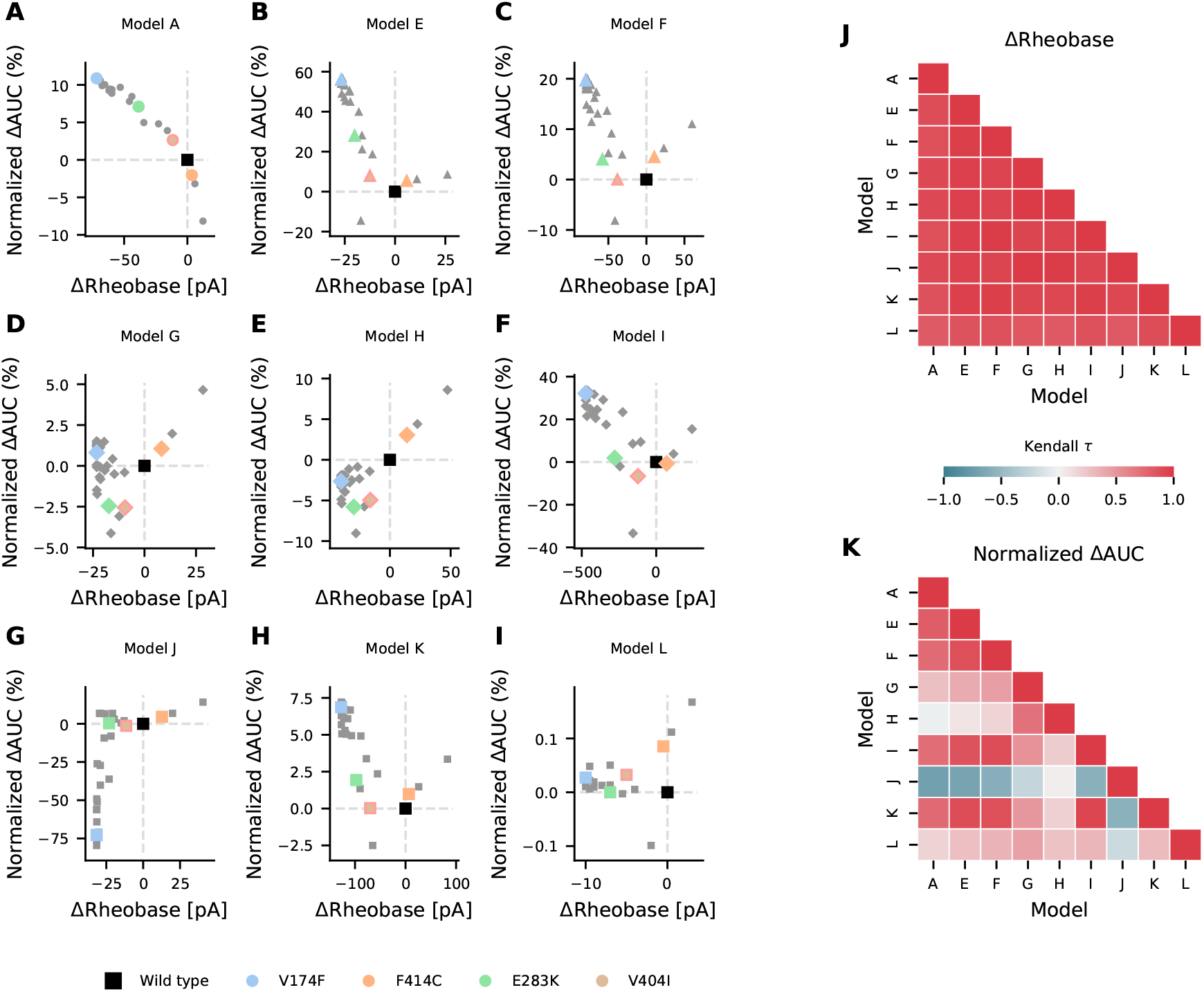
Effects of episodic ataxia type 1 associated *KCNA1* mutations on firing. Effects of *KCNA1* mutations on AUC (percent change in normalized ΔAUC) and rheobase (ΔRheobase) compared to wild type for model H (A), model E (B), model G (C), model A (D), model F (E), model J (F), model L (G), model I (H) and model K (I). All *KCNA1* Mutations are marked in grey with the V174F, F414C, E283K, and V404I *KCNA1* mutations highlighted in color for each model. Pairwise Kendall rank correlation coefficients (Kendall *τ*) between the effects of *KCNA1* mutations on rheobase and on AUC are shown in J and K respectively. Marker shape is indicative of model/firing type, and grey dashed lines denote the quadrants of firing characterization (see Figure 2).

## Discussion

To compare the effects of ion channel mutations on neuronal firing of different neuron types, a diverse set of conductance-based models was used and the effect of changes in individual channel properties across conductance-based neuronal models. Additionally, the effects of episodic ataxia type 1 associated (EA1) *KCNA1* mutations were simulated. Changes to single ionic current properties, as well as known EA1 associated *KCNA1* mutations showed consistent effects on the rheobase across cell types, whereas the effects on AUC of the steady-state fI-curve depended on the cell type. Our results demonstrate that loss of function (LOF) and gain of function (GOF) on the biophysical level cannot be uniquely transferred to the level of neuronal firing. Thus, the effects caused by different mutations depend on the properties of the other ion channels expressed in a cell and are therefore depend on the channel ensemble of a specific cell type.

### Firing Frequency Analysis

Although, firing differences can be characterized by an area under the curve of the fI curve for fixed current steps this approach characterizes firing as a mixture of key features: rheobase and the initial slope of the fI curve. By probing rheobase directly and using an AUC relative to rheobase, we disambiguate these features and enable insights into the effects on rheobase and initial fI curve steepness. This increases the specificity of our understanding of how ion channel mutations alter firing across cells types and enable classification as described in Figure 2. Importantly, in cases when ion channel mutations alter rheobase and initial fI curve steepness in ways that opposing effects on firing (upper left and bottom right quadrants of Figure 2 B) this disamgibuation is important for understanding the outcome of the mutation. In these cases, the regime the neuron is operating in is vital in determining the cells firing outcome. If it is in its excitable regime and only occasionally generates an action potential, then the effect on the rheobase is more important. If it is firing periodically with high rates, then the change in AUC might be more relevant.

### Modelling Limitations

The models used here are simple and while they all capture key aspects of the firing dynamics for their respective cell, they fall short of capturing the complex physiology and biophysics of real cells. However, for the purpose of understanding how different cell-types, or current environments, contribute to diversity in firing outcomes of ion channel mutations, the fidelity of the models to the physiological cells they represent is of a minor concern. For exploring possible cell-type specific effects, variety in currents and dynamics across models is of utmost importance. With this context in mind, the collection of models used here are labelled as models A-L to highlight that the physiological cells they represent is not of chief concern, but rather that the collection of models with different attributes respond heterogeneously to the same perturbation. Additionally, the development of more realistic models is a high priority and will enable cell-type specific predictions that may aid in precision medicine approaches. Thus, weight should not be put on any single predicted firing outcome here in a specific model, but rather on the differences in outcomes that occur across the cell-type spectrum the models used here represent.

### Neuronal Diversity

The nervous system consists of a vastly diverse and heterogenous collection of neurons with variable properties and characteristics including diverse combinations and expression levels of ion channels which are vital for neuronal firing dynamics.

Advances in high-throughput techniques have enabled large-scale investigation into single-cell properties across the CNS (Poulin et al., 2016) that have revealed large diversity in neuronal gene expression, morphology and neuronal types in the motor cortex (Scala et al., 2021), neocortex (Cadwell et al., 2016, 2020), GABAergic neurons in the cortex and retina (Huang and Paul, 2019; Laturnus et al., 2020), cerebellum (Kozareva et al., 2021), spinal cord (Alkaslasi et al., 2021), visual cortex (Gouwens et al., 2019) as well as the retina (Baden et al., 2016; Berens and Euler, 2017; Voigt et al., 2019; Yan et al., 2020a,b).

Diversity across neurons is not limited to gene expression and can also be seen electrophysiologically (Baden et al., 2016; Berens and Euler, 2017; Cadwell et al., 2020; Gouwens et al., 2018,2019; Scala et al., 2021; Tripathy et al., 2015, 2017) with correlations existing between gene expression and electrophysiological properties (Tripathy et al., 2017). At the ion channel level, diversity exists not only between the specific ion channels the different cell types express but heterogeneity also exists in ion channel expression levels within cell types (Barreiro et al., 2012; Goaillard and Marder, 2021; Marder and Taylor, 2011). As ion channel properties and expression levels are key deter-minents of neuronal dynamics and firing (Århem and Blomberg, 2007; Balachandar and Prescott, 2018; Gu and Chen, 2014; Gu et al., 2014; Kispersky et al., 2012; Qi et al., 2013; Zeberg et al., 2010, 2015; Zhou et al., 2020) neurons with different ion channel properties and expression levels display different firing properties.

To capture the diversity in neuronal ion channel expression and its relevance in the outcome of ion channel mutations, we used multiple neuronal models with different ionic currents and underlying firing dynamics here.

### Ionic Current Environments Determine the Effect of Ion Channel Mutations

To our knowledge, no comprehensive evaluation of how ionic current environment and cell type affect the outcome of ion channel mutations have been reported. However, comparisons between the effects of such mutations between certain cell types were described. For instance, the R1648H mutation in *SCN1A* does not alter the excitability of cortical pyramidal neurons, but causes hypoexcitability of adjacent inhibitory GABAergic neurons (Hedrich et al., 2014). In the CA3 region of the hippocampus, the equivalent mutation in *SCN8A*, R1627H, increases the excitability of pyramidal neurons and decreases the excitability of parvalbumin positive interneurons (Makinson et al., 2016). Additionally, the L858H mutation in Na_V_1.7, associated with erythermyalgia, has been shown to cause hypoexcitability in sympathetic ganglion neurons and hyperexcitability in dorsal root ganglion neurons (Rush et al., 2006; Waxman, 2007). The differential effects of L858H Na_V_1.7 on firing is dependent on the presence or absence of another sodium channel, namely the Na_V_1.8 subunit (Rush et al., 2006; Waxman, 2007). These findings, in concert with our findings emphasize that the ionic current environment in which a channelopathy occurs is vital in determining the outcomes of the channelopathy on firing.

Cell type specific differences in ionic current properties are important in the effects of ion channel mutations. However, within a cell type heterogeneity in channel expression levels exists and it is often desirable to generate a population of neuronal models and to screen them for plausibility to biological data in order to capture neuronal population diversity (Marder and Taylor, 2011; O’Leary and Marder, 2016). The models we used here are originally generated by characterization of current gating properties and by fitting of maximal conductances to experimental data (Alexander et al., 2019; Otsuka et al., 2004; Pospischil et al., 2008; Ranjan et al., 2019). This practice of fixing maximal conductances based on experimental data is limiting as it does not reproduce the variability in channel expression and neuronal firing behavior of a heterogeneous neuron population (Verma et al., 2020). For example, a model derived from the mean conductances in a neuronal sub-population within the stomatogastric ganglion, the so-called “one-spike bursting” neurons fire three spikes instead of one per burst due to an L-shaped distribution of sodium and potassium conductances (Golowasch et al., 2002). Multiple sets of conductances can give rise to the same patterns of activity also termed degeneracy and differences in neuronal dynamics may only be evident with perturbations (Goaillard and Marder, 2021; Marder and Taylor, 2011). The variability in ion channel expression often correlates with the expression of other ion channels (Goaillard and Marder, 2021) and neurons whose behavior is similar may possess correlated variability across different ion channels resulting in stability in the neuronal phenotype (Lamb and Calabrese, 2013; Soofi et al., 2012; Taylor et al., 2009). The variability of ionic currents and degeneracy of neurons may account, at least in part, for the observation that the effect of toxins within a neuronal type is frequently not constant (Khaliq and Raman, 2006; Puopolo et al., 2007; Ransdell et al., 2013).

### Effects of *KCNA1* Mutations

Changes in delayed rectifier potassium currents, analogous to those seen in LOF *KCNA1* mutations, change the underlying firing dynamics of the Hodgkin Huxley model result in reduced thresholds for repetitive firing and thus contribute to increased excitability (Hafez and Gottschalk, 2020). Although the Hodgkin Huxley delayed rectifier lacks inactivation, the increases in excitability observed by Hafez and Gottschalk (2020) are in line with our simulation-based predictions of the outcomes of *KCNA1* mutations. LOF *KCNA1* mutations generally increase neuronal excitability, however the varying susceptibility on rheobase and different effects on AUC of the fI-curve of *KCNA1* mutations across models are indicative that a certain cell type specific complexity exists. Increased excitability is seen experimentally with K_V_1.1 null mice (Smart et al., 1998; Zhou et al., 1998), with pharmacological K_V_1.1 block (Chi and Nicol, 2007; Morales-Villagrán et al., 1996) and by Hafez and Gottschalk (2020) with simulation-based predictions of *KCNA1* mutations. Contrary to these results, Zhao et al. (2020) predicted *in silico* that the depolarizing shifts seen as a result of *KCNA1* mutations broaden action potentials and interfere negatively with high frequency action potential firing. However, they varied stimulus duration between different models and therefore comparability of firing rates is lacking in this study.

In our simulations, different current properties alter the impact of *KCNA1* mutations on firing as evident in the differences seen in the impact of I_A_ and I_K_V_1.1_ in the Cb stellate and STN model families on *KCNA1* mutation firing. This highlights that not only knowledge of the biophysical properties of a channel but also its neuronal expression and other neuronal channels present is vital for the holistic understanding of the effects of a given ion channel mutation both at the single cell and network level.

### Loss or Gain of Function Characterizations Do Not Fully Capture Ion Channel Mutation Effects on Firing

The effects of changes in channel properties depend in part on the neuronal model in which they occur and can be seen in the variance of correlations (especially in AUC of the fI-curve) across models for a given current property change. Therefore, relative conductances and gating properties of currents in the ionic current environment in which an alteration in current properties occurs plays an important role in determining the outcome on firing. The use of LOF and GOF is useful at the level of ion channels to indicate whether a mutation results in more or less ionic current. However, the extension of this thinking onto whether mutations induce LOF or GOF at the level of neuronal firing based on the ionic current LOF/GOF is problematic due to the dependency of neuronal firing changes on the ionic channel environment. Thus, the direct leap from current level LOF/GOF characterizations to effects on firing without experimental or modelling-based evidence, although tempting, should be refrained from and viewed with caution when reported. This is especially relevant in the recent development of personalized medicine for channelopathies, where a patient’s specific channelopathy is identified and used to tailor treatments (Ackerman et al., 2013; Brunklaus et al., 2022; Gnecchi et al., 2021; Hedrich et al., 2021; Helbig and Ellis, 2020; Musto et al., 2020; Weber et al., 2017). However, in these cases the effects of specific ion channel mutations are often characterized based on ionic currents in expression systems and classified as LOF or GOF to aid in treatment decisions (Brunklaus et al., 2022; Johannesen et al., 2021; Musto et al., 2020). Although positive treatment outcomes occur with sodium channel blockers in patients with GOF Na_V_1.6 mutations, patients with both LOF and GOF Na_V_1.6 mutations can benefit from treatment with sodium channel blockers (Johannesen et al., 2021). This example suggests that the relationship between effects at the level of ion channels and effects at the level of firing and therapeutics is not linear or evident without further contextual information.

Therefore, the transferring of LOF or GOF from the current to the firing level should be used with caution; the cell type in which the mutant ion channel is expressed may provide valuable insight into the functional consequences of an ion channel mutation. Experimental assessment of the effects of a patient’s specific ion channel mutation *in vivo* is not generally feasible at a large scale. Therefore, modelling approaches investigating the effects of patient specific channelopathies provide an alternative bridge between characterization of changes in biophysical properties of ionic currents and the firing consequences of these effects. In both experimental and modelling investigation into the effects of ion channel mutations on neuronal firing the specific cell-type dependency should be considered.

The effects of altered ion channel properties on firing is generally influenced by the other ionic currents present in the cell. In channelopathies the effect of a given ion channel mutation on neuronal firing therefore depends on the cell type in which those changes occur (Hedrich et al., 2014; Makinson et al., 2016; Rush et al., 2006; Waxman, 2007). Although certain complexities of neurons such as differences in cell-type sensitivities to current property changes, interactions between ionic currents, cell morphology and subcellular ion channel distribution are neglected here, it is likely that this increased complexity *in vivo* would contribute to the cell-type dependent effects on neuronal firing. The complexity and nuances of the nervous system, including cell-type dependent firing effects of channelopathies explored here, likely underlie shortcomings in treatment approaches in patients with channelopathies. Accounting for cell-type dependent firing effects provides an opportunity to further the efficacy and precision in personalized medicine approaches.

With this study we suggest that cell-type specific effects are vital to a full understanding of the effects of channelopathies at the level of neuronal firing. Furthermore, we highlight the use of modelling approaches to enable relatively fast and efficient insight into channelopathies.

## Supporting information

ExtendedData1

## Extended Data

Extended Data 1: Python code for simulations and analysis in zip file. Simulation code for each model, the sensitivity analysis of each model, the simulation of *KCNA1* mutations in each model, and all analysis are provided herein.

**Figure 1-1:**
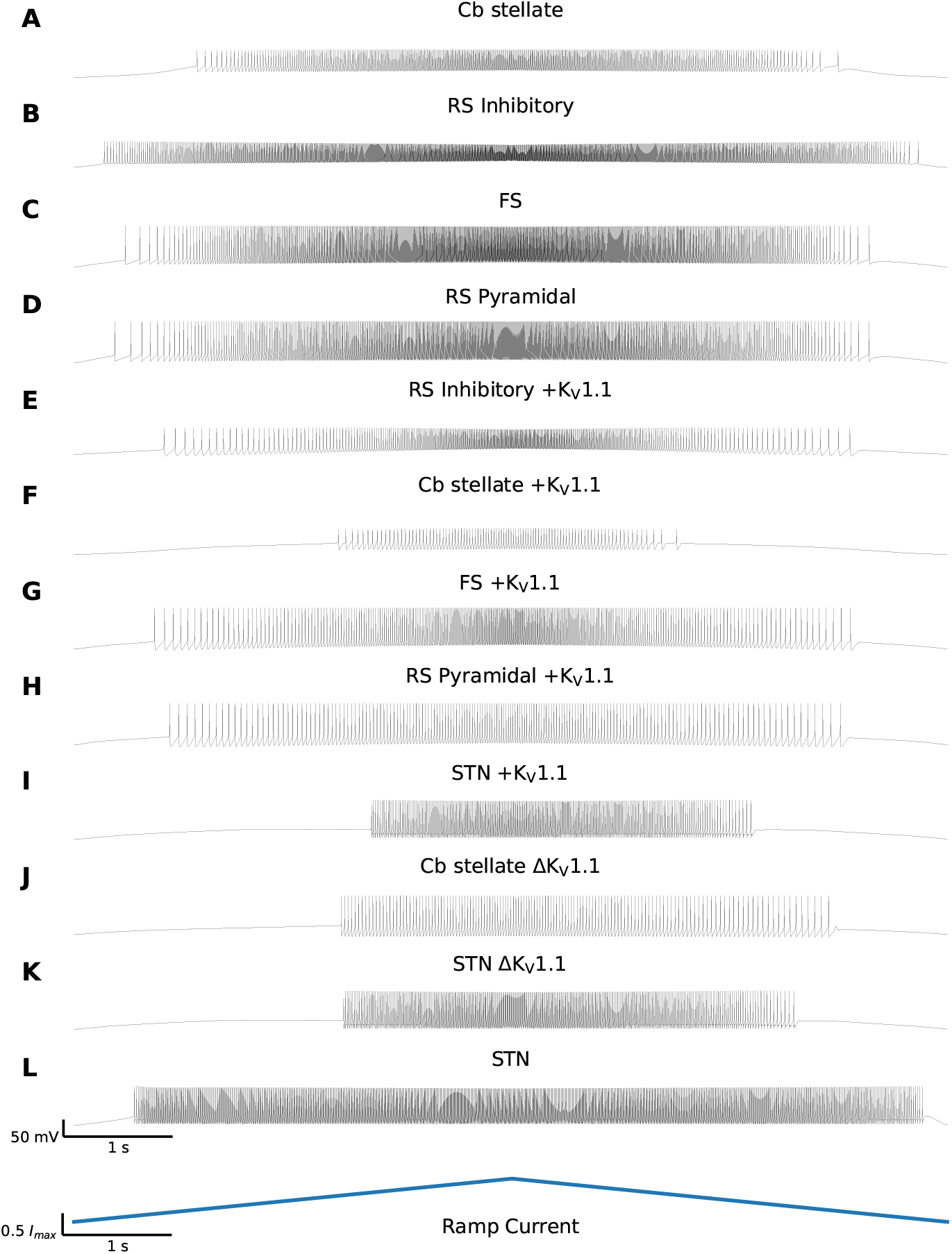
Diversity in Neuronal Model Firing Responses to a Current Ramp. Spike trains for Cb stellate (A), RS inhibitory (B), FS (C), RS pyramidal (D), RS inhibitory +K_V_1.1 (E), Cb stellate +K_V_1.1 (F), FS +K_V_1.1 (G), RS pyramidal +K_V_1.1 (H), STN +K_V_1.1 (I), Cb stellate ΔK_V_1.1 (J), STN ΔK_V_1.1 (K), and STN (L) neuron models in response to a slow ascending current ramp followed by the descending version of the current ramp (bottom). Models are ordered based on the qualitative fI curve sorting in Figure 1. The current at which firing begins in response to an ascending current ramp and the current at which firing ceases in response to a descending current ramp are depicted on the frequency current (fI) curves in Figure 1 for each model.

