## ExtendedData1 for "Loss or Gain of Function? Effects of Ion Channel Mutations on Neuronal Firing Depend on the Cell Type": Plotting_data_collection.py

```
"""
Script to analyse and collect data from simulations


"""
__author__ = "Nils A. Koch"
__copyright__ = 'Copyright 2022, Nils A. Koch'
__license__ = "MIT"
import pandas as pd
import h5py
import json
import os
import numpy as np
from ast import literal_eval
import string
import matplotlib.cm as cm

# %% rheo_{}_ex.csv, AUC_{}_ex.csv #####################################################################################
# Collect examples of effects of a shift, a slope change and a change in g on AUC and rheobase
## AUC ###################
AUC_shift = pd.DataFrame(columns=['alteration', 'RS Pyramidal', 'RS Inhibitory', 'FS', 'IB',
                                   'RS Pyramidal +$K_V1.1$','RS Inhibitory +$K_V1.1$', 'FS +$K_V1.1$','Cb stellate',
                                  'Cb stellate +$K_V1.1$', 'Cb stellate $\Delta$$K_V1.1$', 'STN', 'STN +$K_V1.1$',
                                  'STN $\Delta$$K_V1.1$'])

AUC_slope = pd.DataFrame(columns=['alteration', 'RS Pyramidal', 'RS Inhibitory', 'FS', 'IB',
                                   'RS Pyramidal +$K_V1.1$','RS Inhibitory +$K_V1.1$', 'FS +$K_V1.1$','Cb stellate',
                                  'Cb stellate +$K_V1.1$', 'Cb stellate $\Delta$$K_V1.1$', 'STN', 'STN +$K_V1.1$',
                                  'STN $\Delta$$K_V1.1$'])

AUC_g = pd.DataFrame(columns=['alteration', 'RS Pyramidal', 'RS Inhibitory', 'FS', 'IB',
                                   'RS Pyramidal +$K_V1.1$','RS Inhibitory +$K_V1.1$', 'FS +$K_V1.1$','Cb stellate',
                                  'Cb stellate +$K_V1.1$', 'Cb stellate $\Delta$$K_V1.1$', 'STN', 'STN +$K_V1.1$',
                                  'STN $\Delta$$K_V1.1$'])

script_dir = os.path.dirname(os.path.realpath("__file__"))
fname = os.path.join(script_dir, )
models = ['RS_pyramidal', 'RS_inhib', 'FS','RS_pyramidal_Kv', 'RS_inhib_Kv', 'FS_Kv',  'Cb_stellate','Cb_stellate_Kv',
          'Cb_stellate_Kv_only','STN','STN_Kv', 'STN_Kv_only']
model_labels = ['RS Pyramidal','RS Inhibitory', 'FS','RS Pyramidal +$K_V1.1$','RS Inhibitory +$K_V1.1$',
                'FS +$K_V1.1$','Cb stellate','Cb stellate +$K_V1.1$',
                'Cb stellate $\Delta$$K_V1.1$','STN','STN +$K_V1.1$', 'STN $\Delta$$K_V1.1$']

shift_interest = 'n'
for i in range(len(models)):
    with open('../Sensitivity_Analysis/Data/SA_summary_df/{}_shift_AUC.json'.format(models[i])) as json_file:
        data = pd.read_json(json_file, convert_dates=False, convert_axes=False)
        data.replace(0., np.NaN, inplace=True)
        data = (data - data.loc['0', :])/ data.loc['0', :] # normalize AUC
        data.sort_index(inplace=True)
        AUC_shift[model_labels[i]] =data[shift_interest]
AUC_shift['alteration'] = AUC_shift.index


slope_interest = 's'
for i in range(len(models)):
    with open('../Sensitivity_Analysis/Data/SA_summary_df/{}_slope_AUC.json'.format(models[i])) as json_file:
        data = pd.read_json(json_file, convert_dates=False, convert_axes=False)
        data.replace(0., np.NaN, inplace=True)
        data = (data - data.loc['1.0', :])/ data.loc['1.0', :] # normalize AUC
        data.sort_index(inplace=True)
        try:
            AUC_slope[model_labels[i]] = data[slope_interest]
        except:
            pass
AUC_slope['alteration'] = AUC_slope.index


g_interest = 'Kd'
for i in range(len(models)):
    with open('../Sensitivity_Analysis/Data/SA_summary_df/{}_g_AUC.json'.format(models[i])) as json_file:
        data = pd.read_json(json_file, convert_dates=False, convert_axes=False)
        data.replace(0., np.NaN, inplace=True)
        data = (data - data.loc['1.0', :])/ data.loc['1.0', :] # normalize AUC
        data.sort_index(inplace=True)
        AUC_g[model_labels[i]] =data[g_interest]
AUC_g['alteration'] = AUC_g.index

AUC_shift.to_csv('AUC_shift_ex.csv')
AUC_slope.to_csv('AUC_slope_ex.csv')
AUC_g.to_csv('AUC_g_ex.csv')


## rheobase ###################
rheo_shift = pd.DataFrame(columns=['alteration', 'RS Pyramidal', 'RS Inhibitory', 'FS', 'IB',
                                   'RS Pyramidal +$K_V1.1$','RS Inhibitory +$K_V1.1$', 'FS +$K_V1.1$','Cb stellate',
                                  'Cb stellate +$K_V1.1$', 'Cb stellate $\Delta$$K_V1.1$', 'STN', 'STN +$K_V1.1$',
                                  'STN $\Delta$$K_V1.1$'])

rheo_slope = pd.DataFrame(columns=['alteration', 'RS Pyramidal', 'RS Inhibitory', 'FS', 'RS Pyramidal +$K_V1.1$',
                                   'RS Inhibitory +$K_V1.1$', 'FS +$K_V1.1$', 'Cb stellate', 'Cb stellate +$K_V1.1$',
                                  'Cb stellate $\Delta$$K_V1.1$', 'STN', 'STN +$K_V1.1$','STN $\Delta$$K_V1.1$'])

rheo_g = pd.DataFrame(columns=['alteration', 'RS Pyramidal', 'RS Inhibitory', 'FS', 'IB',
                                   'RS Pyramidal +$K_V1.1$','RS Inhibitory +$K_V1.1$', 'FS +$K_V1.1$','Cb stellate',
                                  'Cb stellate +$K_V1.1$', 'Cb stellate $\Delta$$K_V1.1$', 'STN', 'STN +$K_V1.1$',
                                  'STN $\Delta$$K_V1.1$'])

script_dir = os.path.dirname(os.path.realpath("__file__"))
fname = os.path.join(script_dir, )

models = ['RS_pyramidal', 'RS_inhib', 'FS', 'RS_pyramidal_Kv', 'RS_inhib_Kv', 'FS_Kv', 'Cb_stellate', 'Cb_stellate_Kv', 'Cb_stellate_Kv_only', 'STN',
          'STN_Kv',
          'STN_Kv_only']
model_labels = ['RS Pyramidal','RS Inhibitory', 'FS','RS Pyramidal +$K_V1.1$', 'RS Inhibitory +$K_V1.1$', 'FS +$K_V1.1$',
                'Cb stellate', 'Cb stellate +$K_V1.1$', 'Cb stellate $\Delta$$K_V1.1$', 'STN', 'STN +$K_V1.1$',
                'STN $\Delta$$K_V1.1$']

shift_interest = 's'
for i in range(len(models)):
    with open('../Sensitivity_Analysis/Data/SA_summary_df/{}_shift_rheo.json'.format(models[i])) as json_file:
        data = pd.read_json(json_file, convert_dates=False, convert_axes=False)
        data.replace(0., np.NaN, inplace=True)
        data = (data - data.loc['0', :]) #/ data.loc['0', :]  # normalize AUC
        data.sort_index(inplace=True)
        try:
            rheo_shift[model_labels[i]] = data[shift_interest]
        except:
            pass
rheo_shift['alteration'] = rheo_shift.index

slope_interest = 'u'
for i in range(len(models)):
    with open('../Sensitivity_Analysis/Data/SA_summary_df/{}_slope_rheo.json'.format(models[i])) as json_file:

        data = pd.read_json(json_file, convert_dates=False, convert_axes=False)
        data.replace(0., np.NaN, inplace=True)
        data = (data - data.loc['1.0', :]) #/ data.loc['1.0', :]  # normalize AUC
        data.sort_index(inplace=True)
        try:
            rheo_slope[model_labels[i]] = data[slope_interest]
        except:
            pass
rheo_slope['alteration'] = rheo_slope.index

g_interest = 'Leak'
for i in range(len(models)):
    with open('../Sensitivity_Analysis/Data/SA_summary_df/{}_g_rheo.json'.format(models[i])) as json_file:
        data = pd.read_json(json_file, convert_dates=False, convert_axes=False)
        data.replace(0., np.NaN, inplace=True)
        data = (data - data.loc['1.0', :]) #/ data.loc['1.0', :]  # normalize AUC
        data.sort_index(inplace=True)
        rheo_g[model_labels[i]] = data[g_interest]
rheo_g['alteration'] = rheo_g.index

rheo_shift.to_csv('rheo_shift_ex.csv')
rheo_slope.to_csv('rheo_slope_ex.csv')
rheo_g.to_csv('rheo_g_ex.csv')


# %% Model fI ##########################################################################################################
# for each model generate summary csv of sensitivity analysis results on AUC and rheobase
# | (index) | mag | alt   | type  | F     | I     |
# | 0       | -10 | m     | shift | array | array |

models = ['RS Pyramidal', 'RS Inhibitory', 'FS', 'RS Pyramidal +$\mathrm{K}_{\mathrm{V}}\mathrm{1.1}$',
          'RS Inhibitory +$\mathrm{K}_{\mathrm{V}}\mathrm{1.1}$', 'FS +$\mathrm{K}_{\mathrm{V}}\mathrm{1.1}$',
          'Cb stellate', 'Cb stellate +$\mathrm{K}_{\mathrm{V}}\mathrm{1.1}$',
          'Cb stellate $\Delta$$\mathrm{K}_{\mathrm{V}}\mathrm{1.1}$', 'STN',
          'STN +$\mathrm{K}_{\mathrm{V}}\mathrm{1.1}$', 'STN $\Delta$$\mathrm{K}_{\mathrm{V}}\mathrm{1.1}$']

model_save_name = {'RS Pyramidal': 'RS_pyr_posp', 'RS Inhibitory': 'RS_inhib_posp', 'FS': 'FS_posp',
                   'RS Pyramidal +$\mathrm{K}_{\mathrm{V}}\mathrm{1.1}$':'RS_pyr_Kv',
                   'RS Inhibitory +$\mathrm{K}_{\mathrm{V}}\mathrm{1.1}$':'RS_inhib_Kv',
                   'FS +$\mathrm{K}_{\mathrm{V}}\mathrm{1.1}$':'FS_Kv',
                   'Cb stellate': 'Cb_stellate',
                   'Cb stellate +$\mathrm{K}_{\mathrm{V}}\mathrm{1.1}$':'Cb_stellate_Kv',
                   'Cb stellate $\Delta$$\mathrm{K}_{\mathrm{V}}\mathrm{1.1}$':'Cb_stellate_Kv_only',
                   'STN': 'STN',
                   'STN +$\mathrm{K}_{\mathrm{V}}\mathrm{1.1}$': 'STN_Kv',
                   'STN $\Delta$$\mathrm{K}_{\mathrm{V}}\mathrm{1.1}$': 'STN_Kv_only'}
top_file_dir = '../Sensitivity_Analysis/Data'
data_dir = {'RS Pyramidal': os.path.join(top_file_dir, 'RS_pyramidal_{}_fI.json'),
            'RS Inhibitory': os.path.join(top_file_dir, 'RS_inhib_{}_fI.json'),
            'FS': os.path.join(top_file_dir, 'FS_{}_fI.json'),
            'RS Pyramidal +$\mathrm{K}_{\mathrm{V}}\mathrm{1.1}$': os.path.join(top_file_dir, 'RS_pyramidal_{}_fI.json'),
            'RS Inhibitory +$\mathrm{K}_{\mathrm{V}}\mathrm{1.1}$': os.path.join(top_file_dir, 'RS_inhib_{}_fI.json'),
            'FS +$\mathrm{K}_{\mathrm{V}}\mathrm{1.1}$': os.path.join(top_file_dir, 'FS_{}_fI.json'),
            'Cb stellate': os.path.join(top_file_dir, 'Cb_stellate_{}_fI.json'),
            'Cb stellate +$\mathrm{K}_{\mathrm{V}}\mathrm{1.1}$': os.path.join(top_file_dir, 'Cb_stellate_Kv_{}_fI.json'),
            'Cb stellate $\Delta$$\mathrm{K}_{\mathrm{V}}\mathrm{1.1}$': os.path.join(top_file_dir, 'Cb_stellate_Kv_only_{}_fI.json'),
            'STN': os.path.join(top_file_dir, 'STN_{}_fI.json'),
            'STN +$\mathrm{K}_{\mathrm{V}}\mathrm{1.1}$': os.path.join(top_file_dir, 'STN_Kv_{}_fI.json'),
            'STN $\Delta$$\mathrm{K}_{\mathrm{V}}\mathrm{1.1}$': os.path.join(top_file_dir, 'STN_Kv_only_{}_fI.json')}

I_dir = {'RS Pyramidal': os.path.join(top_file_dir, 'RS_pyramidal_{}_I_mag.json'),
            'RS Inhibitory': os.path.join(top_file_dir, 'RS_pyramidal_{}_I_mag.json'),
            'FS': os.path.join(top_file_dir, 'RS_pyramidal_{}_I_mag.json'),
            'RS Pyramidal +$\mathrm{K}_{\mathrm{V}}\mathrm{1.1}$': os.path.join(top_file_dir, 'RS_pyramidal_{}_I_mag.json'),
            'RS Inhibitory +$\mathrm{K}_{\mathrm{V}}\mathrm{1.1}$': os.path.join(top_file_dir, 'RS_inhib_{}_I_mag.json'),
            'FS +$\mathrm{K}_{\mathrm{V}}\mathrm{1.1}$': os.path.join(top_file_dir, 'FS_{}_I_mag.json'),
            'Cb stellate': os.path.join(top_file_dir, 'Cb_stellate_{}_I_mag.json'),
            'Cb stellate +$\mathrm{K}_{\mathrm{V}}\mathrm{1.1}$': os.path.join(top_file_dir, 'Cb_stellate_Kv_{}_I_mag.json'),
            'Cb stellate $\Delta$$\mathrm{K}_{\mathrm{V}}\mathrm{1.1}$': os.path.join(top_file_dir, 'Cb_stellate_Kv_only_{}_I_mag.json'),
            'STN': os.path.join(top_file_dir, 'STN_{}_I_mag.json'),
            'STN +$\mathrm{K}_{\mathrm{V}}\mathrm{1.1}$': os.path.join(top_file_dir, 'STN_Kv_{}_I_mag.json'),
            'STN $\Delta$$\mathrm{K}_{\mathrm{V}}\mathrm{1.1}$': os.path.join(top_file_dir, 'STN_Kv_only_{}_I_mag.json')}


def concat_dfs(df, data_dir, I_dir, type='shift'):
    with open(data_dir.format(type)) as json_file:
        data = pd.read_json(json_file, convert_axes=False)
    with open(I_dir.format(type)) as json_file:
        I = pd.read_json(json_file, convert_axes=False)
    for c in data.columns:
        # print(c)
        df = df.append(pd.DataFrame(data=(np.array([data.index,['{}'.format(c) for i in range(0, len(data[c]))],
                                                      [type for i in range(0, len(data[c]))], data[c], I[c]]).T),
                                      columns=['mag','alt', 'type', 'F', 'I']))
    return df


summary_df = pd.DataFrame(columns=['mag','alt', 'type', 'F', 'I'])
for m in models:
    print(m)
    df = pd.DataFrame(columns=['mag','alt', 'type', 'F', 'I'])
    df = concat_dfs(df, data_dir[m], I_dir[m], type='shift')
    df = concat_dfs(df, data_dir[m], I_dir[m], type='slope')
    df = concat_dfs(df, data_dir[m], I_dir[m], type='g')
    folder = os.path.join(os.path.dirname(os.path.realpath("__file__")), 'Model_fI')
    if not os.path.isdir(folder):
        os.makedirs(folder)
    df.to_csv(os.path.join(folder, '{}_fI.csv'.format(model_save_name[m])))


# %% firing_values.csv, model_spiking.csv, model_F_inf.csv #############################################################
# generate firing_values.csv with values for hysteresis from ramp current input
# generate model_spiking.csv with example model response to step current
# generate model_F_inf.csv with model fI curve data
import numpy as np
import pandas as pd

models = ['RS_pyramidal', 'RS_inhib', 'FS', 'RS_pyramidal_Kv', 'RS_inhib_Kv', 'FS_Kv', 'Cb_stellate', 'Cb_stellate_Kv',
          'Cb_stellate_Kv_only', 'STN', 'STN_Kv', 'STN_Kv_only']
model_names = ['RS pyramidal', 'RS inhibitory', 'FS', 'RS pyramidal +Kv1.1', 'RS inhibitory +Kv1.1', 'FS +Kv1.1',
               'Cb stellate', 'Cb stellate +Kv1.1', 'Cb stellate $\Delta$Kv1.1', 'STN', 'STN +Kv1.1',
               'STN $\Delta$Kv1.1']
firing_values = pd.DataFrame(columns=models, index=['spike_ind', 'ramp_up', 'ramp_down'])
models = ['RS_pyr', 'RS_pyr_Kv', 'RS_inhib', 'RS_inhib_Kv', 'FS', 'FS_Kv',
          'Cb_stellate', 'Cb_stellate_Kv', 'Cb_stellate_Delta_Kv',
          'STN', 'STN_Kv', 'STN_Delta_Kv']
col_names = ['I', 'I_inhib']
for mod in models: col_names.append(mod)
model_F_inf = pd.DataFrame(columns=col_names)
col_names = ['t']
for mod in models: col_names.append(mod)
spiking = pd.DataFrame(columns=col_names)

# index for example trace
spike_ind = {'RS_pyramidal': 60, 'RS_inhib':25, 'FS':50, 'RS_pyramidal_Kv':60,
            'RS_inhib_Kv':50, 'FS_Kv':50, 'Cb_stellate':60, 'Cb_stellate_Kv':130,
            'Cb_stellate_Kv_only':75, 'STN': 25, 'STN_Kv':95, 'STN_Kv_only':80}

for model_name in models:
    folder = '../Neuron_models/{}'.format(model_name)
    fname = os.path.join(folder, "{}.hdf5".format(model_name))  # RS_inhib
    with h5py.File(fname, "r+") as f:
        I_mag = np.arange(f['data'].attrs['I_low'], f['data'].attrs['I_high'],
                          (f['data'].attrs['I_high'] - f['data'].attrs['I_low']) / f['data'].attrs['stim_num']) * 1000
        start = np.int(f['data'].attrs['initial_period'] * 1 / f['data'].attrs['dt'])
        stim_len = np.int((f['data'].attrs['stim_time'] - start) * f['data'].attrs['dt'])
        time = np.arange(0, stim_len, f['data'].attrs['dt'])
        spiking[model_name] = f['data']['V_m'][spike_ind[model_name]][start:]
        model_F_inf[model_name] = f['analysis']['F_inf'][:]
        firing_values.loc['spike_ind', model_name] = I_mag[spike_ind[model_name]]
        firing_values.loc['ramp_down', model_name] = f['analysis']['ramp_I_down'][()]
        firing_values.loc['ramp_up', model_name] = f['analysis']['ramp_I_up'][()]
firing_values.to_csv('firing_values.csv')
spiking.to_csv('model_spiking.csv')
model_F_inf.to_csv('model_F_inf.csv')
# %% model_ramp.csv ####################################################################################################
# generate model_ramp.csv with model responses to ramp input
# | (index) | t | models ....
import numpy as np
import pandas as pd

models = ['RS_pyramidal_Kv', 'RS_inhib_Kv', 'FS_Kv', 'Cb_stellate', 'Cb_stellate_Kv', 'Cb_stellate_Kv_only', 'STN',
          'STN_Kv', 'STN_Kv_only']
model_names = ['RS pyramidal', 'RS inhibitory', 'FS', 'Cb stellate', 'Cb stellate +Kv1.1', 'Cb stellate $\Delta$Kv1.1',
               'STN', 'STN +Kv1.1', 'STN $\Delta$Kv1.1']
col_names = ['t']
for mod in models: col_names.append(mod)
model_ramp = pd.DataFrame(columns=col_names)
sec = 4
dt = 0.01
ramp_len = int(sec * 1000 * 1 / dt)
t_ramp = np.arange(0, ramp_len) * dt
model_ramp.loc[:, 't'] = t_ramp

for model_name in models:
    folder = '../Neuron_models/{}'.format(model_name)
    fname = os.path.join(folder, "{}.hdf5".format(model_name))  # RS_inhib
    with h5py.File(fname, "r+") as f:
        model_ramp.loc[:, model_name] = f['analysis']['V_m_ramp'][()]

model_ramp.to_csv('model_ramp.csv')

#%% sim_mut_AUC.csv, sim_mut_rheo.csv ##################################################################################
# generate mutation plot data in sim_mut_AUC.csv and sim_mut_rheo.csv for the AUC and rheobase effects respectively for
# each mutation in each model
mutations = json.load(open("../KCNA1_mutations/mutations_effects_dict.json"))
keys_to_remove = ['V408L', 'T226R', 'R239S', 'R324T']
for key in keys_to_remove:
    del mutations[key]
mutations_f = []
mutations_n = []
for mut in mutations:
    mutations_n.append(mut)
    mutations_f.append(mut.replace(" ", "_"))

models = ['RS_pyramidal_Kv', 'RS_inhib_Kv', 'FS_Kv', 'Cb_stellate', 'Cb_stellate_Kv', 'Cb_stellate_Kv_only', 'STN',
          'STN_Kv', 'STN_Kv_only']
model_names = ['RS pyramidal', 'RS inhibitory', 'FS', 'Cb stellate', 'Cb stellate +Kv1.1', 'Cb stellate $\Delta$Kv1.1',
               'STN', 'STN +Kv1.1', 'STN $\Delta$Kv1.1']
AUC = pd.DataFrame(columns=mutations_n)
rheobase = pd.DataFrame(columns=mutations_n)
save_folder = '../KCNA1_mutations'
if not os.path.isdir(save_folder):
    os.makedirs(save_folder)
for model_name in models:
    folder = '../KCNA1_mutations/{}'.format(model_name)
    for mut in list(mutations_n):
        fname = os.path.join(folder, "{}.hdf5".format(mut.replace(" ", "_")))
        with h5py.File(fname, "r+") as f:
            rheobase.loc[mut.replace(" ", "_"), model_name] = f['analysis']['rheobase'][()]
            AUC.loc[mut.replace(" ", "_"), model_name] = f['analysis']['AUC'][()]
AUC.replace(0., np.NaN, inplace=True)
rheobase.replace(0., np.NaN, inplace=True)
rheobase = (rheobase - rheobase.loc['WT', :]) /rheobase.loc['WT', :]
AUC = (AUC - AUC.loc['WT', :]) /AUC.loc['WT', :]
AUC.to_csv(os.path.join(save_folder, 'sim_mut_AUC.csv'))
rheobase.to_csv(os.path.join(save_folder, 'sim_mut_rheobase.csv'))
```
