## ExtendedData1 for "Loss or Gain of Function? Effects of Ion Channel Mutations on Neuronal Firing Depend on the Cell Type": SA_collection.py

```
"""
Script to analyse and collect data from sensitivity analysis simulations


"""
__author__ = "Nils A. Koch"
__copyright__ = 'Copyright 2022, Nils A. Koch'
__license__ = "MIT"

import numpy as np
import h5py
import pandas as pd
import os
from ast import literal_eval
import json
import scipy.stats as stats

# folder
top_dir = '../Sensitivity_Analysis/Data'

#%% create JSON files for each alt type for each model in '../Sensitivity_Analysis/Data/SA_summary_df/file.json'
for dir_name in next(os.walk(top_dir))[1]: # for each folder in top_file_dir (each model)
    folder = os.path.join(top_dir, dir_name)
    print(folder)
    # for each alt_type create pandas dataframe
    shift_AUC = pd.DataFrame()
    shift_rheo = pd.DataFrame()
    shift_fI = pd.DataFrame(dtype=object)
    shift_I_mag = pd.DataFrame(dtype=object)

    slope_AUC = pd.DataFrame()
    slope_rheo = pd.DataFrame()
    slope_fI = pd.DataFrame(dtype=object)
    slope_I_mag = pd.DataFrame(dtype=object)

    g_AUC = pd.DataFrame()
    g_rheo = pd.DataFrame()
    g_fI = pd.DataFrame(dtype=object)
    g_I_mag = pd.DataFrame(dtype=object)


    for root, dirs, files in os.walk(folder):
        for file in files:
            if file.endswith('.hdf5'):
                with h5py.File(os.path.join(folder, file), "r+") as f:
                    alt = f['data'].attrs['alteration']

                    test = f['data'].attrs['alteration_info'].replace(' ', ',')
                    alt_info = literal_eval(test)
                    var = alt_info[0]
                    alt_type = alt_info[1]
                    if alt_type == 'shift':
                        shift_AUC.loc[alt, var] = f['analysis']['AUC'][()]
                        try:
                            shift_rheo.loc[alt, var] = f['analysis']['rheobase'][()]
                        except:
                            print('shift', var, alt)
                        shift_fI.loc[alt, var] = 0
                        shift_fI = shift_fI.astype(object)
                        shift_fI.at[alt, var] = f['analysis']['F_inf'][:].tolist()
                        shift_I_mag.loc[alt, var] = 0
                        shift_I_mag = shift_I_mag.astype(object)
                        shift_I_mag.at[alt, var] = ((np.arange(f['data'].attrs['I_low'], f['data'].attrs['I_high'],
                                                               (f['data'].attrs['I_high'] - f['data'].attrs['I_low']) /
                                                               f['data'].attrs['stim_num'])) * 1000).tolist() #nA
                    elif alt_type == 'slope':
                        slope_AUC.loc[alt, var] = f['analysis']['AUC'][()]
                        try:
                            slope_rheo.loc[alt, var] = f['analysis']['rheobase'][()]
                        except:
                            print('slope', var, alt)
                        slope_fI.loc[alt, var] = 0
                        slope_fI = slope_fI.astype(object)
                        slope_fI.at[alt, var] = f['analysis']['F_inf'][:].tolist()
                        slope_I_mag.loc[alt, var] = 0
                        slope_I_mag = slope_I_mag.astype(object)
                        slope_I_mag.at[alt, var] = ((np.arange(f['data'].attrs['I_low'], f['data'].attrs['I_high'],
                                                               (f['data'].attrs['I_high'] - f['data'].attrs['I_low']) /
                                                               f['data'].attrs['stim_num'])) * 1000).tolist()
                    elif alt_type == 'g':
                        g_AUC.loc[alt, var] = f['analysis']['AUC'][()]
                        try:
                            g_rheo.loc[alt, var] = f['analysis']['rheobase'][()]
                        except:
                            print('g', var, alt)
                        g_fI.loc[alt, var] = 0
                        g_fI = g_fI.astype(object)
                        g_fI.at[alt, var] = f['analysis']['F_inf'][:].tolist()
                        g_I_mag.loc[alt, var] = 0
                        g_I_mag = g_I_mag.astype(object)
                        g_I_mag.at[alt, var] = ((np.arange(f['data'].attrs['I_low'], f['data'].attrs['I_high'],
                                                               (f['data'].attrs['I_high'] - f['data'].attrs['I_low']) /
                                                               f['data'].attrs['stim_num'])) * 1000).tolist()
                    else:
                        print(file, 'Unknown alteration type')

    #save df with folder+alt_type
    save_folder = os.path.join(top_dir, 'SA_summary_df')
    if not os.path.isdir(save_folder):
        os.makedirs(save_folder)
    shift_AUC.to_json(os.path.join(save_folder, '{}_shift_AUC.json'.format(dir_name)))
    shift_rheo.to_json(os.path.join(save_folder, '{}_shift_rheo.json'.format(dir_name)))
    shift_fI.to_json(os.path.join(save_folder, '{}_shift_fI.json'.format(dir_name)))
    shift_I_mag.to_json(os.path.join(save_folder, '{}_shift_I_mag.json'.format(dir_name)))

    slope_AUC.to_json(os.path.join(save_folder, '{}_slope_AUC.json'.format(dir_name)))
    slope_rheo.to_json(os.path.join(save_folder, '{}_slope_rheo.json'.format(dir_name)))
    slope_fI.to_json(os.path.join(save_folder, '{}_slope_fI.json'.format(dir_name)))
    slope_I_mag.to_json(os.path.join(save_folder, '{}_slope_I_mag.json'.format(dir_name)))

    g_AUC.to_json(os.path.join(save_folder, '{}_g_AUC.json'.format(dir_name)))
    g_rheo.to_json(os.path.join(save_folder, '{}_g_rheo.json'.format(dir_name)))
    g_fI.to_json(os.path.join(save_folder, '{}_g_fI.json'.format(dir_name)))
    g_I_mag.to_json(os.path.join(save_folder, '{}_g_I_mag.json'.format(dir_name)))


#%% AUC Correlation analysis
alt_dict = {}
alt_dict['m'] = 'Na activation'
alt_dict['h'] = 'Na inactivation'
alt_dict['n'] = 'K activation'
alt_dict['s'] = '$K_V1.1$ activation'
alt_dict['u'] = '$K_V1.1$inactivation'
alt_dict['a'] = 'A activation'
alt_dict['b'] = 'A inactivation'
alt_dict['n_A'] = 'A activation'
alt_dict['h_A'] = 'A inactivation'
alt_dict['Na'] = 'Na'
alt_dict['Kd'] = 'K'
alt_dict['Kv'] = '$K_V1.1$'
alt_dict['A'] = 'A'
alt_dict['Leak'] = 'Leak'

# models = directory names in top_file_dir and what json files are named
models = ['RS_pyramidal', 'RS_inhib', 'FS', 'RS_pyramidal_Kv', 'RS_inhib_Kv', 'FS_Kv', 'Cb_stellate', 'Cb_stellate_Kv',
          'Cb_stellate_Kv_only', 'STN', 'STN_Kv', 'STN_Kv_only']
# model_names = names of models for json files and ultimately csv files for plotting
model_names = ['RS Pyramidal','RS Inhibitory','FS', 'RS Pyramidal +$K_V1.1$','RS Inhibitory +$K_V1.1$', 'FS +$K_V1.1$',
               'Cb stellate','Cb stellate +$K_V1.1$', 'Cb stellate $\Delta$$K_V1.1$','STN','STN +$K_V1.1$', 'STN $\Delta$$K_V1.1$']

shift_df = pd.DataFrame(columns=['model', 'corr', 'p_value', 'local corr', 'local p_value', 'ratio', '$\Delta V_{1/2}$', 'color'])  # for boxplots
for mod in range(len(models)):
    with open('../Sensitivity_Analysis/Data/SA_summary_df/{}_shift_AUC.json'.format(models[mod])) as json_file:
        df = pd.read_json(json_file)
        df.sort_index(inplace=True)
        df.replace(0., np.NaN, inplace=True)
        df = (df - df.loc[0, :]) / df.loc[0, :]
        zero_ind = np.argwhere(df.index == 0)[0][0]
        ind = [df.index[zero_ind - 1], df.index[zero_ind], df.index[zero_ind + 1]]
        df2 = df.loc[ind, :]
    for c in df.keys():
        tau, p = stats.kendalltau(df.index, df[c], nan_policy='omit')
        tau_l, p_l = stats.kendalltau(df2.index, df2[c], nan_policy='omit')
        ratio_tau = tau_l / tau
        shift_df = shift_df.append(pd.Series([model_names[mod], tau, p,tau_l, p_l, ratio_tau, alt_dict[c],clr_dict[models[mod]]], index=shift_df.columns), ignore_index=True)


# models = directory names in top_file_dir and what json files are named
models = ['RS_pyramidal', 'RS_inhib', 'FS', 'RS_pyramidal_Kv', 'RS_inhib_Kv', 'FS_Kv', 'Cb_stellate', 'Cb_stellate_Kv',
          'Cb_stellate_Kv_only', 'STN', 'STN_Kv', 'STN_Kv_only']
# model_names = names of models for json files and ultimately csv files for plotting
model_names = ['RS Pyramidal','RS Inhibitory','FS', 'RS Pyramidal +$K_V1.1$','RS Inhibitory +$K_V1.1$', 'FS +$K_V1.1$',
               'Cb stellate','Cb stellate +$K_V1.1$', 'Cb stellate $\Delta$$K_V1.1$','STN','STN +$K_V1.1$', 'STN $\Delta$$K_V1.1$']

slope_df = pd.DataFrame(columns=['model', 'corr', 'p_value','local corr', 'local p_value', 'ratio', 'Slope (k)', 'color'])  # for boxplots
for mod in range(len(models)):
    with open('../Sensitivity_Analysis/Data/SA_summary_df//{}_slope_AUC.json'.format(models[mod])) as json_file:
        df = pd.read_json(json_file, convert_dates=False, convert_axes=False)
        df.index = df.index.map(float)
        df.sort_index(inplace=True)
        df.replace(0., np.NaN, inplace=True)
        df = (df - df.loc[1.0, :]) / df.loc[1.0, :]
        zero_ind = np.argwhere(df.index == 1)[0][0]
        ind = [df.index[zero_ind - 1], df.index[zero_ind], df.index[zero_ind + 1]]
        df2 = df.loc[ind, :]
    for c in df.keys():
        tau, p = stats.kendalltau(df.index, df[c], nan_policy='omit')
        tau_l, p_l = stats.kendalltau(df2.index, df2[c], nan_policy='omit')
        ratio_tau = tau_l/tau
        slope_df = slope_df.append(pd.Series([model_names[mod], tau, p, tau_l, p_l, ratio_tau, alt_dict[c],clr_dict[models[mod]]], index=slope_df.columns), ignore_index=True)

# models = directory names in top_file_dir and what json files are named
models = ['RS_pyramidal', 'RS_inhib', 'FS', 'RS_pyramidal_Kv', 'RS_inhib_Kv', 'FS_Kv', 'Cb_stellate', 'Cb_stellate_Kv',
          'Cb_stellate_Kv_only', 'STN', 'STN_Kv', 'STN_Kv_only']
# model_names = names of models for json files and ultimately csv files for plotting
model_names = ['RS Pyramidal','RS Inhibitory','FS', 'RS Pyramidal +$K_V1.1$','RS Inhibitory +$K_V1.1$', 'FS +$K_V1.1$',
               'Cb stellate','Cb stellate +$K_V1.1$', 'Cb stellate $\Delta$$K_V1.1$','STN','STN +$K_V1.1$', 'STN $\Delta$$K_V1.1$']

g_df = pd.DataFrame(columns=['model', 'corr', 'p_value', 'local corr', 'local p_value', 'ratio', 'g', 'color'])  # for boxplots
for mod in range(len(models)):
    with open('../Sensitivity_Analysis/Data/SA_summary_df//{}_g_AUC.json'.format(models[mod])) as json_file:
        df = pd.read_json(json_file, convert_dates=False, convert_axes=False)
        df.index = df.index.map(float)
        df.sort_index(inplace=True)
        df.replace(0., np.NaN, inplace=True)
        df = (df - df.loc[1.0, :]) / df.loc[1.0, :]
        zero_ind = np.argwhere(df.index == 1)[0][0]
        ind = [df.index[zero_ind - 1], df.index[zero_ind], df.index[zero_ind + 1]]
        df2 = df.loc[ind, :]
    for c in df.keys():
        tau, p = stats.kendalltau(df.index, df[c], nan_policy='omit')
        tau_l, p_l = stats.kendalltau(df2.index, df2[c], nan_policy='omit')
        ratio_tau = tau_l / tau
        g_df = g_df.append(pd.Series([model_names[mod], tau, p,tau_l, p_l, ratio_tau, alt_dict[c],clr_dict[models[mod]]], index=g_df.columns), ignore_index=True)

shift_df.to_json('./Sensitivity_analysis/shift_box_kendall_corr.json')
slope_df.to_json('./Sensitivity_analysis/slope_box_kendall_corr.json')
g_df.to_json('./Sensitivity_analysis/g_kendall_corr.json')


#%% rheobase correlation analysis

# models = directory names in top_file_dir and what json files are named
models = ['RS_pyramidal', 'RS_inhib', 'FS', 'RS_pyramidal_Kv', 'RS_inhib_Kv', 'FS_Kv', 'Cb_stellate', 'Cb_stellate_Kv',
          'Cb_stellate_Kv_only', 'STN', 'STN_Kv', 'STN_Kv_only']
# model_names = names of models for json files and ultimately csv files for plotting
model_names = ['RS Pyramidal','RS Inhibitory','FS', 'RS Pyramidal +$K_V1.1$','RS Inhibitory +$K_V1.1$', 'FS +$K_V1.1$',
               'Cb stellate','Cb stellate +$K_V1.1$', 'Cb stellate $\Delta$$K_V1.1$','STN','STN +$K_V1.1$', 'STN $\Delta$$K_V1.1$']

shift_df = pd.DataFrame(columns=['model', 'corr', 'p_value', 'local corr', 'local p_value', 'ratio', '$\Delta V_{1/2}$', 'color'])  # for boxplots
for mod in range(len(models)):
    with open('../Sensitivity_Analysis/Data/SA_summary_df//{}_shift_rheo.json'.format(models[mod])) as json_file:
        df = pd.read_json(json_file)
        df.sort_index(inplace=True)
        df.replace(0., np.NaN, inplace=True)
        df = (df - df.loc[0, :]) / df.loc[0, :]
        zero_ind = np.argwhere(df.index == 0)[0][0]
        ind = [df.index[zero_ind - 1], df.index[zero_ind], df.index[zero_ind + 1]]
        df2 = df.loc[ind, :]
    for c in df.keys():
        tau, p = stats.kendalltau(df.index, df[c], nan_policy='omit')
        tau_l, p_l = stats.kendalltau(df2.index, df2[c], nan_policy='omit')
        ratio_tau = tau_l / tau
        shift_df = shift_df.append(pd.Series([model_names[mod], tau, p,tau_l, p_l, ratio_tau, alt_dict[c],clr_dict[models[mod]]], index=shift_df.columns), ignore_index=True)


# models = directory names in top_file_dir and what json files are named
models = ['RS_pyramidal', 'RS_inhib', 'FS', 'RS_pyramidal_Kv', 'RS_inhib_Kv', 'FS_Kv', 'Cb_stellate', 'Cb_stellate_Kv',
          'Cb_stellate_Kv_only', 'STN', 'STN_Kv', 'STN_Kv_only']
# model_names = names of models for json files and ultimately csv files for plotting
model_names = ['RS Pyramidal','RS Inhibitory','FS', 'RS Pyramidal +$K_V1.1$','RS Inhibitory +$K_V1.1$', 'FS +$K_V1.1$',
               'Cb stellate','Cb stellate +$K_V1.1$', 'Cb stellate $\Delta$$K_V1.1$','STN','STN +$K_V1.1$', 'STN $\Delta$$K_V1.1$']

slope_df = pd.DataFrame(columns=['model', 'corr', 'p_value','local corr', 'local p_value', 'ratio', 'Slope (k)', 'color'])  # for boxplots
for mod in range(len(models)):
    with open('../Sensitivity_Analysis/Data/SA_summary_df//{}_slope_rheo.json'.format(models[mod])) as json_file:
        df = pd.read_json(json_file, convert_dates=False, convert_axes=False)
        df.index = df.index.map(float)
        df.sort_index(inplace=True)
        df.replace(0., np.NaN, inplace=True)
        df = (df - df.loc[1.0, :]) / df.loc[1.0, :]
        zero_ind = np.argwhere(df.index == 1)[0][0]
        ind = [df.index[zero_ind - 1], df.index[zero_ind], df.index[zero_ind + 1]]
        df2 = df.loc[ind, :]
    for c in df.keys():
        tau, p = stats.kendalltau(df.index, df[c], nan_policy='omit')
        tau_l, p_l = stats.kendalltau(df2.index, df2[c], nan_policy='omit')
        ratio_tau = tau_l/tau
        slope_df = slope_df.append(pd.Series([model_names[mod], tau, p, tau_l, p_l, ratio_tau, alt_dict[c],clr_dict[models[mod]]], index=slope_df.columns), ignore_index=True)

# models = directory names in top_file_dir and what json files are named
models = ['RS_pyramidal', 'RS_inhib', 'FS', 'RS_pyramidal_Kv', 'RS_inhib_Kv', 'FS_Kv', 'Cb_stellate', 'Cb_stellate_Kv',
          'Cb_stellate_Kv_only', 'STN', 'STN_Kv', 'STN_Kv_only']
# model_names = names of models for json files and ultimately csv files for plotting
model_names = ['RS Pyramidal','RS Inhibitory','FS', 'RS Pyramidal +$K_V1.1$','RS Inhibitory +$K_V1.1$', 'FS +$K_V1.1$',
               'Cb stellate','Cb stellate +$K_V1.1$', 'Cb stellate $\Delta$$K_V1.1$','STN','STN +$K_V1.1$', 'STN $\Delta$$K_V1.1$']

g_df = pd.DataFrame(columns=['model', 'corr', 'p_value', 'local corr', 'local p_value', 'ratio', 'g', 'color'])  # for boxplots
for mod in range(len(models)):
    with open('../Sensitivity_Analysis/Data/SA_summary_df//{}_g_rheo.json'.format(models[mod])) as json_file:
        df = pd.read_json(json_file, convert_dates=False, convert_axes=False)
        df.index = df.index.map(float)
        df.sort_index(inplace=True)
        df.replace(0., np.NaN, inplace=True)
        df = (df - df.loc[1.0, :]) / df.loc[1.0, :]
        zero_ind = np.argwhere(df.index == 1)[0][0]
        ind = [df.index[zero_ind - 1], df.index[zero_ind], df.index[zero_ind + 1]]
        df2 = df.loc[ind, :]
    for c in df.keys():
        tau, p = stats.kendalltau(df.index, df[c], nan_policy='omit')
        tau_l, p_l = stats.kendalltau(df2.index, df2[c], nan_policy='omit')
        ratio_tau = tau_l / tau
        g_df = g_df.append(pd.Series([model_names[mod], tau, p,tau_l, p_l, ratio_tau, alt_dict[c],clr_dict[models[mod]]], index=g_df.columns), ignore_index=True)

shift_df.to_json('./Sensitivity_analysis/rheo_shift_box_kendall_corr.json')
slope_df.to_json('./Sensitivity_analysis/rheo_slope_box_kendall_corr.json')
g_df.to_json('./Sensitivity_analysis/rheo_g_kendall_corr.json')

#%% create csv files for plotting
# | (index) | model | corr   | p_value  | g | color
#% AUC
AUC_shift_json = pd.read_json('./Sensitivity_analysis/shift_kendall_corr_rel.json', orient='records')
AUC_slope_json = pd.read_json('./Sensitivity_analysis/slope_kendall_corr_rel.json', orient='records') #, lines=True)
AUC_g_json = pd.read_json('./Sensitivity_analysis/g_kendall_corr_rel.json', orient='records')
AUC_shift_df = AUC_shift_json[['model', 'corr', 'p_value', 'g', 'color']]
AUC_slope_df = AUC_slope_json[['model', 'corr', 'p_value', 'g', 'color']]
AUC_g_df = AUC_g_json[['model', 'corr', 'p_value', 'g', 'color']]
AUC_shift_df.to_csv('AUC_shift_corr.csv')
AUC_slope_df.to_csv('AUC_scale_corr.csv')
AUC_g_df.to_csv('AUC_g_corr.csv')

#% rheo
rheo_shift_json = pd.read_json('./Sensitivity_analysis/rheo_shift_kendall_corr.json', orient='records')
rheo_slope_json = pd.read_json('./Sensitivity_analysis/rheo_slope_kendall_corr.json', orient='records') #, lines=True)
rheo_g_json = pd.read_json('./Sensitivity_analysis/rheo_g_kendall_corr.json', orient='records')
rheo_shift_df = rheo_shift_json[['model', 'corr', 'p_value', 'g', 'color']]
rheo_slope_df = rheo_slope_json[['model', 'corr', 'p_value', 'g', 'color']]
rheo_g_df = rheo_g_json[['model', 'corr', 'p_value', 'g', 'color']]
rheo_shift_df.to_csv('rheo_shift_corr.csv')
rheo_slope_df.to_csv('rheo_scale_corr.csv')
rheo_g_df.to_csv('rheo_g_corr.csv')
```
