## ExtendedData1 for "Loss or Gain of Function? Effects of Ion Channel Mutations on Neuronal Firing Depend on the Cell Type": Cb_stellate_fxns_Kv.py

```
"""
Functions for Cb Stellate with Kv1.1 model


"""
__author__ = "Nils A. Koch"
__copyright__ = 'Copyright 2022, Nils A. Koch'
__license__ = "MIT"

import h5py
import scipy
import json
import copy
import os
import gc
import numpy as np
from numba import types, njit
from numba.typed import Dict
from scipy import signal
from Code.Functions.Utility_fxns import stimulus_init, NumpyEncoder


@njit
def Cb_stellate_Kv(V, g, E, I_in, dt, currents_included, stim_time, stim_num, C, shift, scale, b_param,
                             slope_shift, gating, current, ind_dict):
    """ Simulate current and Vm for  Cb stellate cell model from Alexander et al 2019 (https://dx.doi.org/10.1523%2FENEURO.0126-19.2019)
        with Kv1.1 from Ranjan et al. 2019 (https://dx.doi.org/10.3389/fncel.2019.00358) added for I_in

        Parameters
        ----------
        V : float
            Initial membrane potentia
        g : dict
            maximal conductance for currents
        E : dict
            Reversal potentials for currents
        I_in : array
            Current stimulus input to model
        dt : float
            fixed time step
        currents_included : dict
            Boolean to include current in simulation or not
        stim_time :
            length of stimulus ()
        stim_num : int
            number of different stimulus currents
        C : float
            membrane capacitance
        shift : dict
            for each gating parameter
        scale : dict
            for each gating parameter
        b_param : dict
            Boltzmann parameter values for each gating parameter
        slope_shift : dict
            for each gating parameter
        gating : array
            array of gating values over time
        current : array
            array of current values over time
        ind_dict: dict
            Dictionary of indices in arrays for gating variables


        Returns
        -------
        V_m : array
            Simulated membrane potential in response to I_in
        """

    V_m = np.zeros((stim_num, stim_time))
    # initialize gating

    # Na activation
    m_inf = (1 / (1 + np.exp((V - shift["m"] - slope_shift["m"] - b_param["m"][0]) / b_param["m"][1]))) ** \
            b_param["m"][2]
    gating[ind_dict['m'], :] = np.ones((stim_num)) * m_inf

    # Na inactivation
    h_inf = ((1 - b_param["h"][3]) / (1 + np.exp((V - shift["h"] - slope_shift["h"] - b_param["h"][0])
                                                 / b_param["h"][1])) + b_param["h"][3]) ** b_param["h"][2]
    gating[ind_dict['h'], :] = np.ones((stim_num)) * h_inf

    # K activation
    n_inf = (1 / (1 + np.exp((V - shift["n"] - slope_shift["n"] - b_param["n"][0]) / b_param["n"][1]))) ** \
            b_param["n"][2]
    gating[ind_dict['n'], :] = np.ones((stim_num)) * n_inf

    # A-type K activation
    n_A_inf = (1 / (1 + np.exp((V - shift["n_A"] - slope_shift["n_A"] - b_param["n_A"][0]) / b_param["n_A"][1]))) ** \
              b_param["n_A"][2]
    gating[ind_dict['n_A'], :] = np.ones((stim_num)) * n_A_inf

    # A-type K inactivation
    h_A_inf = (1 / (1 + np.exp((V - shift["h_A"] - slope_shift["h_A"] - b_param["h_A"][0]) / b_param["h_A"][1]))) ** \
              b_param["h_A"][2]
    gating[ind_dict['h_A'], :] = np.ones((stim_num)) * h_A_inf

    # T-type Ca activation
    m_T_inf = (1 / (1 + np.exp((V - shift["m_T"] - slope_shift["m_T"] - b_param["m_T"][0]) / b_param["m_T"][1]))) ** \
              b_param["m_T"][2]
    gating[ind_dict['m_T'], :] = np.ones((stim_num)) * m_T_inf

    # T-type Ca inactivation
    h_T_inf = (1 / (1 + np.exp((V - shift["h_T"] - slope_shift["h_T"] - b_param["h_T"][0]) / b_param["h_T"][1]))) ** \
              b_param["h_T"][2]
    gating[ind_dict['h_T'], :] = np.ones((stim_num)) * h_T_inf

    # Kv1.1 activation
    s_inf = (1 / (1 + np.exp((V - shift["s"] - slope_shift["s"] - b_param["s"][0]) / b_param["s"][1]))) ** \
            b_param["s"][2]
    gating[ind_dict['s'], :] = np.ones((stim_num)) * s_inf

    # Kv1.1 inactivation
    u_inf = ((1 - b_param["u"][3]) / (1 + np.exp((V - shift["u"] - slope_shift["u"] - b_param["u"][0])
                                                 / b_param["u"][1])) + b_param["u"][3]) ** b_param["u"][2]
    gating[ind_dict['u'], :] = np.ones((stim_num)) * u_inf

    # mutant Kv1.1 activation
    s_mut_inf = (1 / (1 + np.exp((V - shift["s_mut"] - slope_shift["s_mut"] - b_param["s_mut"][0]) / b_param["s_mut"][1]))) ** \
                b_param["s_mut"][2]
    gating[ind_dict['s_mut'], :] = np.ones((stim_num)) * s_mut_inf

    # mutant Kv1.1 inactivation
    u_mut_inf = ((1 - b_param["u_mut"][3]) / (1 + np.exp((V - shift["u"] - slope_shift["u_mut"] - b_param["u_mut"][0])
                       / b_param["u_mut"][1])) + b_param["u_mut"][3]) ** b_param["u_mut"][2]
    gating[ind_dict['u_mut'], :] = np.ones((stim_num)) * u_mut_inf

    # initialize currents and Vm
    current[ind_dict['Na'], :] = np.zeros((stim_num))
    current[ind_dict['Kd'], :] = np.zeros((stim_num))
    current[ind_dict['Kv'], :] = np.zeros((stim_num))
    current[ind_dict['Kv_mut'], :] = np.zeros((stim_num))
    current[ind_dict['Leak'], :] = np.zeros((stim_num))
    current[ind_dict['A'], :] = np.zeros((stim_num))
    current[ind_dict['T'], :] = np.zeros((stim_num))
    V_m[:, 0] = V
    t = 1
    while t < stim_time:
        # Na activation
        m_inf = (1 / (1 + np.exp((V_m[:, t - 1] - shift["m"] - slope_shift["m"] - b_param["m"][0]) / b_param["m"][1]))) ** \
                b_param["m"][2]
        gating[ind_dict['m'], :] = m_inf
        gating[ind_dict['m'], :][gating[ind_dict['m'], :] > 1.] = 1.
        gating[ind_dict['m'], :][gating[ind_dict['m'], :] < 0.] = 0.

        # Na inactivation
        h_inf = ((1 - b_param["h"][3]) / (1 + np.exp((V_m[:, t - 1] - shift["h"] - slope_shift["h"] - b_param["h"][0])
                                                     / b_param["h"][1])) + b_param["h"][3]) ** b_param["h"][2]
        tau_h = 0.1 + (2 * 322 * 46) / (4 * np.pi * (V_m[:, t - 1] - shift["h"] - -74) ** 2 + (46 ** 2))
        h_dot = (h_inf - gating[ind_dict['h'], :]) * (1 / (tau_h * scale["h"]))
        gating[ind_dict['h'], :] = gating[ind_dict['h'], :] + h_dot * dt
        gating[ind_dict['h'], :][gating[ind_dict['h'], :] > 1.] = 1.
        gating[ind_dict['h'], :][gating[ind_dict['h'], :] < 0.] = 0.

        # K activation
        n_inf = (1 / (1 + np.exp((V_m[:, t - 1] - shift["n"] - slope_shift["n"] - b_param["n"][0]) / b_param["n"][1]))) ** \
                b_param["n"][2]
        tau_n = 6 / (1 + np.exp((V_m[:, t - 1] - shift["n"] + 23) / 15))
        n_dot = (n_inf - gating[ind_dict['n'], :]) * (1 / (tau_n * scale["n"]))
        gating[ind_dict['n'], :] = gating[ind_dict['n'], :] + n_dot * dt
        gating[ind_dict['n'], :][gating[ind_dict['n'], :] > 1.] = 1.
        gating[ind_dict['n'], :][gating[ind_dict['n'], :] < 0.] = 0.

        # A-type K activation
        n_A_inf = (1 / (1 + np.exp((V_m[:, t - 1] - shift["n_A"] - slope_shift["n_A"] - b_param["n_A"][0]) / b_param["n_A"][1]))) ** \
                  b_param["n_A"][2]
        tau_n_A = 5
        n_A_dot = (n_A_inf - gating[ind_dict['n_A'], :]) * (1 / (tau_n_A * scale["n_A"]))
        gating[ind_dict['n_A'], :] = gating[ind_dict['n_A'], :] + n_A_dot * dt
        gating[ind_dict['n_A'], :][gating[ind_dict['n_A'], :] > 1.] = 1.
        gating[ind_dict['n_A'], :][gating[ind_dict['n_A'], :] < 0.] = 0.

        # A-type K inactivation
        h_A_inf = (1 / (1 + np.exp((V_m[:, t - 1] - shift["h_A"] - slope_shift["h_A"] - b_param["h_A"][0]) / b_param["h_A"][1]))) ** \
                  b_param["h_A"][2]
        tau_h_A = 10
        h_A_dot = (h_A_inf - gating[ind_dict['h_A'], :]) * (1 / (tau_h_A * scale["h_A"]))
        gating[ind_dict['h_A'], :] = gating[ind_dict['h_A'], :] + h_A_dot * dt
        gating[ind_dict['h_A'], :][gating[ind_dict['h_A'], :] > 1.] = 1.
        gating[ind_dict['h_A'], :][gating[ind_dict['h_A'], :] < 0.] = 0.

        # T-type Ca activation
        m_T_inf = (1 / (1 + np.exp((V_m[:, t - 1] - shift["m_T"] - slope_shift["m_T"] - b_param["m_T"][0]) / b_param["m_T"][1]))) ** \
                  b_param["m_T"][2]
        gating[ind_dict['m_T'], :] = m_T_inf
        gating[ind_dict['m_T'], :][gating[ind_dict['m_T'], :] > 1.] = 1.
        gating[ind_dict['m_T'], :][gating[ind_dict['m_T'], :] < 0.] = 0.

        # T-type Ca inactivation
        h_T_inf = (1 / (1 + np.exp((V_m[:, t - 1] - shift["h_T"] - slope_shift["h_T"] - b_param["h_T"][0]) / b_param["h_T"][1]))) ** \
                  b_param["h_T"][2]
        tau_h_T = 15
        h_T_dot = (h_T_inf - gating[ind_dict['h_T'], :]) * (1 / (tau_h_T * scale["h_T"]))
        gating[ind_dict['h_T'], :] = gating[ind_dict['h_T'], :] + h_T_dot * dt
        gating[ind_dict['h_T'], :][gating[ind_dict['h_T'], :] > 1.] = 1.
        gating[ind_dict['h_T'], :][gating[ind_dict['h_T'], :] < 0.] = 0.

        # Kv1.1 activation
        s_inf = (1 / (1 + np.exp((V_m[:, t - 1] - shift["s"] - slope_shift["s"] - b_param["s"][0]) / b_param["s"][1]))) ** \
                b_param["s"][2]
        temp = 36
        vBreak = -46.7
        amp1 = 52.7
        amp2 = 15.98
        vh1 = -49.87
        vh2 = -41.64
        offset = 0.9
        slope1 = 5
        slope2 = 24.99
        sigswitch = 1 / (1 + np.exp((V_m[:, t - 1] - shift["s"] - vBreak) / 3))
        sig1 = sigswitch * amp1 / (1 + np.exp((V_m[:, t - 1] - shift["s"] - vh1) / slope1))
        sig2 = (1 - sigswitch) * offset + (amp2 - offset) / (1 + np.exp((V_m[:, t - 1] - shift["s"] - vh2) / slope2))
        sTauFunc = sig1 + sig2
        s_tau_Q10 = (7.54 * np.exp(-(V_m[:, t - 1] - shift["s"]) / 379.7) * np.exp(-temp / 35.66)) ** ((temp - 25) / 10)
        tau_s = sTauFunc / s_tau_Q10
        s_dot = (s_inf - gating[ind_dict['s'], :]) * (1 / (tau_s * scale["s"]))
        gating[ind_dict['s'], :] = gating[ind_dict['s'], :] + s_dot * dt
        gating[ind_dict['s'], :][gating[ind_dict['s'], :] > 1.] = 1.
        gating[ind_dict['s'], :][gating[ind_dict['s'], :] < 0.] = 0.

        # Kv1.1 inactivation
        u_inf = ((1 - b_param["u"][3]) / (1 + np.exp((V_m[:, t - 1] - shift["u"] - slope_shift["u"] - b_param["u"][0])
                                                     / b_param["u"][1])) + b_param["u"][3]) ** b_param["u"][2]
        u_tau_Q10 = 2.7 ** ((temp - 25) / 10)
        tau_u = (86.86 + (408.78 / (1 + np.exp((V_m[:, t - 1] - shift["u"] + 13.6) / 7.46)))) / u_tau_Q10
        u_dot = (u_inf - gating[ind_dict['u'], :]) * (1 / (tau_u * scale["u"]))
        gating[ind_dict['u'], :] = gating[ind_dict['u'], :] + u_dot * dt
        gating[ind_dict['u'], :][gating[ind_dict['u'], :] > 1.] = 1.
        gating[ind_dict['u'], :][gating[ind_dict['u'], :] < 0.] = 0.

        # mutant Kv1.1 activation
        s_mut_inf = (1 / (1 + np.exp((V_m[:, t - 1] - shift["s_mut"] - slope_shift["s_mut"] - b_param["s_mut"][0]) / b_param["s_mut"][1]))) ** \
                    b_param["s_mut"][2]

        s_mut_dot = (s_mut_inf - gating[ind_dict['s_mut'], :]) * (1 / (tau_s * scale["s_mut"]))
        gating[ind_dict['s_mut'], :] = gating[ind_dict['s_mut'], :] + s_mut_dot * dt
        gating[ind_dict['s_mut'], :][gating[ind_dict['s_mut'], :] > 1.] = 1.
        gating[ind_dict['s_mut'], :][gating[ind_dict['s_mut'], :] < 0.] = 0.

        # mutant Kv1.1 inactivation
        u_mut_inf = ((1 - b_param["u_mut"][3]) / (1 + np.exp((V_m[:, t - 1] - shift["u_mut"] - slope_shift["u_mut"] - b_param["u_mut"][0])
                           / b_param["u_mut"][1])) + b_param["u_mut"][3]) ** b_param["u_mut"][2]
        u_mut_dot = (u_mut_inf - gating[ind_dict['u_mut'], :]) * (1 / (tau_u * scale["u_mut"]))
        gating[ind_dict['u_mut'], :] = gating[ind_dict['u_mut'], :] + u_mut_dot * dt
        gating[ind_dict['u_mut'], :][gating[ind_dict['u_mut'], :] > 1.] = 1.
        gating[ind_dict['u_mut'], :][gating[ind_dict['u_mut'], :] < 0.] = 0.

        # update currents
        current[ind_dict['Leak'], :] = g["Leak"] * (V_m[:, t - 1] - E["Leak"]) * currents_included["Leak"]
        current[ind_dict['Na'], :] = g["Na"] * (gating[ind_dict['m'], :] ** 3) * gating[ind_dict['h'], :] * (V_m[:, t - 1] - E["Na"]) * \
                                     currents_included["Na"]
        current[ind_dict['Kd'], :] = g["Kd"] * (gating[ind_dict['n'], :] ** 4) * (V_m[:, t - 1] - E["K"]) * \
                                     currents_included["Kd"]
        current[ind_dict['Kv'], :] = g["Kv"] * gating[ind_dict['s'], :] * gating[ind_dict['u'], :] * (
                V_m[:, t - 1] - E["K"]) * currents_included["Kv"]
        current[ind_dict['Kv_mut'], :] = g["Kv_mut"] * gating[ind_dict['s_mut'], :] * gating[ind_dict['u_mut'], :] * \
                                         (V_m[:, t - 1] - E["K"]) * currents_included["Kv_mut"]
        current[ind_dict['A'], :] = g["A"] * gating[ind_dict['n_A'], :] * gating[ind_dict['h_A'], :] * (
                V_m[:, t - 1] - E["K"]) * currents_included["A"]
        current[ind_dict['T'], :] = g["T"] * gating[ind_dict['m_T'], :] * gating[ind_dict['h_T'], :] * (
                V_m[:, t - 1] - E["Ca"]) * currents_included["T"]

        # update dV/dt
        V_dot = (1 / C) * (I_in[t, :] - (current[ind_dict['Leak'], :] + current[ind_dict['Na'], :] +
                                         current[ind_dict['Kd'], :] + current[ind_dict['A'], :] +
                                         current[ind_dict['Kv'], :] + current[ind_dict['Kv_mut'], :] +
                                         current[ind_dict['T'], :]))
        # update V_m
        V_m[:, t] = V_m[:, t - 1] + V_dot * dt
        t += 1

    return V_m


def Cb_stellate_Kv_mut(V_init, g, E, I_in, dt, currents_included, stim_time, stim_num, C, shift, scale, b_param,
                       slope_shift, gating, current, prominence, desired_AUC_width, mutations, mut, folder, high,
                       low, number_steps, initial_period, sec):
    """ Simulate mutations for Cb Stellate Kv model

           Parameters
           ----------
           V_init : float
               Initial membrane potential
           g : dict
               maximal conductance for currents
           E : dict
               Reversal potentials for currents
           I_in : array
               Current stimulus input to model
           dt : float
               fixed time step
           currents_included : dict
               Boolean to include current in simulation or not
           stim_time :
               length of stimulus ()
           stim_num : int
               number of different stimulus currents
           C : float
               membrane capacitance
           shift : dict
               for each gating parameter
           scale : dict
               for each gating parameter
           b_param : dict
               Boltzmann parameter values for each gating parameter
           slope_shift : dict
               for each gating parameter
           gating : array
               array of gating values over time
           current : array
               array of current values over time
           prominence : float
               required spike prominence
           desired_AUC_width : float
               width of AUC from rheobase to rheobase+desired_AUC_width
           mutations : dict
               dictionary of mutation effects
           mut : str
               mutation
           folder : str
               folder to save hdf5 file in
           high : float
               upper current step magnitude
           low : float
               lowest current step magnitude
           number_steps : int
               number of current steps between low and high
           initial_period : float
               length  of  I = 0 before current step
           sec:
               length of current step in seconds

           Returns
           -------
           saves mut.hdf5 file to folder
           """
    # setup for simulation
    g_multi = copy.deepcopy(g)
    E_multi = copy.deepcopy(E)
    b_param_multi = copy.deepcopy(b_param)
    shift_multi = copy.deepcopy(shift)
    scale_multi = copy.deepcopy(scale)
    slope_shift_multi = copy.deepcopy(slope_shift)
    currents_included_multi = copy.deepcopy(currents_included)
    gating2 = copy.deepcopy(gating)
    current2 = copy.deepcopy(current)

    g_effect = Dict.empty(key_type=types.unicode_type, value_type=types.float64, )
    for k, v in g_multi.items():
        g_effect[k] = v
    b_param_effect = Dict.empty(key_type=types.unicode_type, value_type=types.float64[:], )
    for k, v in b_param_multi.items():
        b_param_effect[k] = np.array(v)
    shift2 = Dict.empty(key_type=types.unicode_type, value_type=types.float64, )
    for k, v in shift_multi.items():
        shift2[k] = v
    scale2 = Dict.empty(key_type=types.unicode_type, value_type=types.float64, )
    for k, v in scale_multi.items():
        scale2[k] = v
    slope_shift2 = Dict.empty(key_type=types.unicode_type, value_type=types.float64, )
    for k, v in slope_shift_multi.items():
        slope_shift2[k] = v
    E2 = Dict.empty(key_type=types.unicode_type, value_type=types.float64, )
    for k, v in E_multi.items():
        E2[k] = v
    currents_included2 = Dict.empty(key_type=types.unicode_type, value_type=types.boolean, )
    for k, v in currents_included_multi.items():
        currents_included2[k] = v
    ind_dict = Dict.empty(key_type=types.unicode_type, value_type=types.int64, )
    i = 0
    for var in np.array(['m', 'h', 'n', 'n_A', 'h_A', 'm_T', 'h_T', 's', 'u', 's_mut', 'u_mut']):
        ind_dict[var] = i
        i += 1
    i = 0
    for var in np.array(['Na', 'Kd', 'A', 'Kv', 'Kv_mut', 'T', 'Leak']):
        ind_dict[var] = i
        i += 1

    # effect of mutation
    g_effect['Kv_mut'] = g['Kv_mut'] * mutations[mut]['g_ratio']  # mutation_properties[mut, 2]  # g ratio A
    b_param_effect['s_mut'][0] = b_param_effect['s_mut'][0] + mutations[mut]['activation_Vhalf_diff']
    b_param_effect['s_mut'][1] = b_param_effect['s_mut'][1] * mutations[mut]['activation_k_ratio']

    # initial simulation for current range from low to high
    V_m = Cb_stellate_Kv(V_init * np.ones((stim_num)), g_effect, E2, I_in, dt, currents_included2, stim_time, stim_num,
                         C, shift2, scale2, b_param_effect, slope_shift2, gating2, current2, ind_dict)
    stim_start = np.int(initial_period * 1 / dt)
    min_spike_height = V_m[0, stim_start] + prominence

    # create hdf5 file for the mutation and save data and metadata
    fname = os.path.join(folder, "{}.hdf5".format(mut.replace(" ", "_")))
    with h5py.File(fname, "a") as f:
        data = f.create_group("data")
        data.create_dataset('V_m', data=V_m, dtype='float64', compression="gzip", compression_opts=9)
        metadata = {'I_high': high, 'I_low': low, 'stim_num': number_steps, 'stim_time': stim_time, 'sec': sec, 'dt': dt,
                    'initial_period': initial_period, 'g': json.dumps(dict(g_effect)), 'E': json.dumps(dict(E2)), 'C': C,
                    'V_init': V_init, 'ind_dict': json.dumps(dict(ind_dict)),
                    'b_param': json.dumps(dict(b_param_effect), cls=NumpyEncoder),
                    'currents': json.dumps(dict(currents_included2)), 'shift': json.dumps(dict(shift2)),
                    'scale': json.dumps(dict(scale2)), 'slope_shift': json.dumps(dict(slope_shift2)),
                    'prominence': prominence, 'desired_AUC_width': desired_AUC_width, 'mut': str(mut),
                    'min_spike_height': min_spike_height,}
        data.attrs.update(metadata)

    # firing frequency analysis and rheobase
    with h5py.File(fname, "r+") as f:
        I_mag = np.arange(low, high, (high - low) / stim_num)
        analysis = f.create_group("analysis")
        dtyp = h5py.special_dtype(vlen=np.dtype('float64'))
        for i in np.array(['spike_times', 'spike_times_rheo', 'ISI', 'Freq', 'amplitude',]):
            analysis.create_dataset(i, (I_mag.shape[0],), dtype=dtyp, compression="gzip", compression_opts=9)
        analysis.create_dataset('F_inf', (I_mag.shape[0],), dtype='float64')
        analysis.create_dataset('F_null', (I_mag.shape[0],), dtype='float64')
        analysis.create_dataset('AP', (I_mag.shape[0],), dtype='bool', compression="gzip", compression_opts=9)
        analysis.create_dataset('AP_rheo', (I_mag.shape[0],), dtype='bool', compression="gzip", compression_opts=9)
        analysis.create_dataset('rheobase', (1,), dtype='float64', compression="gzip", compression_opts=9)

        # Firing analysis
        for stim in range(I_mag.shape[0]):
            # spike detection
            ap_all, ap_prop = scipy.signal.find_peaks(V_m[stim, stim_start:], min_spike_height,
                                                      prominence=prominence, distance=1 * 1 / dt)
            analysis['spike_times'][stim] = ap_all * dt
            if analysis['spike_times'][stim].size == 0:
                analysis['AP'][stim] = False
            else:
                analysis['AP'][stim] = True
            analysis['amplitude'][stim] = np.array(ap_prop["peak_heights"])
            analysis['ISI'][stim] = np.array([x - analysis['spike_times'][stim][i - 1] if i else None for i, x in
                                              enumerate(analysis['spike_times'][stim])][1:])  # msec
            analysis['Freq'][stim] = 1 / (analysis['ISI'][stim] / 1000)

            if analysis['Freq'][stim].size != 0:
                analysis['F_null'][stim] = analysis['Freq'][stim][0]
                last_second_spikes = analysis['spike_times'][stim] > np.int(
                    f['data'].attrs['stim_time'] * dt - (500 + initial_period))
                if np.any(last_second_spikes == True):
                    last_second_spikes_1 = np.delete(last_second_spikes,
                                                     0)  # remove first spike because it has no ISI/freq
                    analysis['F_inf'][stim] = np.mean(analysis['Freq'][stim][last_second_spikes_1])
            else: # if no ISI detected
                analysis['F_null'][stim] = 0
                analysis['F_inf'][stim] = 0

        # rheobase
        # find where the first AP occurs
        if np.any(analysis['AP'][:] == True):
            # setup current range to be between the current step before and with first AP
            I_first_spike = I_mag[np.argwhere(analysis['AP'][:] == True)[0][0]]
            I_before_spike = I_mag[np.argwhere(analysis['AP'][:] == True)[0][0] - 1]
            analysis.create_dataset('I_first_spike', data=I_first_spike, dtype='float64')
            analysis.create_dataset('I_before_spike', data=I_before_spike, dtype='float64')
            number_steps_rheo = 100
            stim_time, I_in_rheo, stim_num_rheo, V_m_rheo = stimulus_init(I_before_spike, I_first_spike,
                                                                               number_steps_rheo, initial_period,
                                                                               dt, sec)

            # simulate response to this current range and save
            V_m_rheo = Cb_stellate_Kv(V_init * np.ones((stim_num_rheo)), g_effect, E2, I_in_rheo, f['data'].attrs['dt'],
                                      currents_included2, stim_time, stim_num_rheo, f['data'].attrs['C'], shift2,
                                      scale2, b_param_effect, slope_shift2, np.zeros((len(b_param), stim_num_rheo)),
                                      np.zeros((len(currents_included), stim_num_rheo)), ind_dict)
            f['data'].create_dataset('V_m_rheo', data=V_m_rheo, dtype='float64', compression="gzip", compression_opts=9)

            # run AP detection
            stim_start = np.int(initial_period * 1 / dt)
            for stim in range(I_in_rheo.shape[1]):
                ap_all, ap_prop = scipy.signal.find_peaks(V_m_rheo[stim, stim_start:], min_spike_height,
                                                          prominence=prominence, distance=1 * 1 / dt)
                analysis['spike_times_rheo'][stim] = ap_all * dt
                if analysis['spike_times_rheo'][stim].size == 0:
                    analysis['AP_rheo'][stim] = False
                else:
                    analysis['AP_rheo'][stim] = True
            # find rheobase as smallest current step to elicit AP and save
            analysis['rheobase'][:] = I_in_rheo[-1, np.argwhere(analysis['AP_rheo'][:] == True)[0][0]] * 1000

    # AUC
    with h5py.File(fname, "a") as f:
        # find rheobase of steady state firing (F_inf)
        F_inf = f['analysis']['F_inf'][:]
        try:
            # setup current range to be in current step between no and start of steady state firing
            F_inf_start_ind = np.argwhere(F_inf != 0)[0][0]
            dI = (high - low) / stim_num
            I_start = I_mag[F_inf_start_ind - 1]
            I_end = I_mag[F_inf_start_ind]
            I_rel = np.arange(I_start, I_end + (I_end - I_start) / 100, (I_end - I_start) / 100)
            I_rel = np.reshape(I_rel, (1, I_rel.shape[0]))
            I_in = np.zeros((stim_time + np.int(initial_period * 1 / dt), 1)) @ I_rel
            I_in[np.int(initial_period * 1 / dt):, :] = np.ones((stim_time, 1)) @ I_rel
            stim_num = I_in.shape[1]

            # simulate response to this current step range
            V_m_inf_rheo = Cb_stellate_Kv(V_init * np.ones(stim_num), g_effect, E2, I_in, dt, currents_included2,
                                          stim_time, stim_num, C, shift2, scale2, b_param_effect, slope_shift2,
                                          np.zeros((len(b_param), stim_num)),
                                          np.zeros((len(currents_included), stim_num)), ind_dict)

            # save response, analyse and save
            for i in np.array(['spike_times_Finf', 'ISI_Finf', 'Freq_Finf']):  # analysis_names:
                f['analysis'].require_dataset(i, shape=(I_rel.shape[1],), dtype=dtyp, compression="gzip", compression_opts=9)
            f['analysis'].require_dataset('F_inf_Finf', shape=(I_rel.shape[1],), dtype='float64')
            f['analysis'].require_dataset('F_null_Finf', shape=(I_rel.shape[1],), dtype='float64')
            for stim in range(I_rel.shape[1]):
                # spike peak detection
                ap_all, ap_prop = scipy.signal.find_peaks(V_m_inf_rheo[stim, stim_start:], min_spike_height,
                                                          prominence=prominence, distance=1 * 1 / dt)
                f['analysis']['spike_times_Finf'][stim] = ap_all * dt
                f['analysis']['ISI_Finf'][stim] = np.array(
                    [x - f['analysis']['spike_times_Finf'][stim][i - 1] if i else None for i, x in
                     enumerate(f['analysis']['spike_times_Finf'][stim])][1:])  # msec
                f['analysis']['Freq_Finf'][stim] = 1 / (f['analysis']['ISI_Finf'][stim] / 1000)
                if f['analysis']['Freq_Finf'][stim].size != 0:
                    last_second_spikes = f['analysis']['spike_times_Finf'][stim] > np.int(
                        f['data'].attrs['stim_time'] * dt - (500 + f['data'].attrs['initial_period']))
                    if np.any(last_second_spikes == True):
                        last_second_spikes_1 = np.delete(last_second_spikes,
                                                         0)  # remove first spike because it has no ISI/freq
                        f['analysis']['F_inf_Finf'][stim] = np.mean(
                            f['analysis']['Freq_Finf'][stim][last_second_spikes_1])
                    else:
                        f['analysis']['F_null_Finf'][stim] = 0
                        f['analysis']['F_inf_Finf'][stim] = 0

            # find AUC relative to F_inf rheobase
            # setup current inputs with current range around F_inf rheobase
            I_first_inf = I_rel[0, np.argwhere(f['analysis']['F_inf_Finf'][:] != 0)[0][0]]
            I_start = I_first_inf - 0.00005
            I_end = I_first_inf + desired_AUC_width * high
            I_rel2 = np.arange(I_start, I_end + (I_end - I_start) / 200, (I_end - I_start) / 200)
            I_rel2 = np.reshape(I_rel2, (1, I_rel2.shape[0]))
            I_in2 = np.zeros((stim_time + np.int(initial_period * 1 / dt), 1)) @ I_rel2
            I_in2[np.int(initial_period * 1 / dt):, :] = np.ones((stim_time, 1)) @ I_rel2
            stim_num = I_in2.shape[1]

            # simulate response to this current step range
            V_m_AUC = Cb_stellate_Kv(V_init * np.ones(stim_num), g_effect, E2, I_in2, dt, currents_included2, stim_time,
                                     stim_num, C, shift2, scale2, b_param_effect, slope_shift2,
                                     np.zeros((len(b_param), stim_num)), np.zeros((len(currents_included), stim_num)),
                                     ind_dict)

            # save data and analysis for this current step range including AUC
            f['data'].require_dataset('V_m_AUC', shape=V_m_AUC.shape, dtype='float64')
            f['data']['V_m_AUC'][:] = V_m_AUC
            dtyp = h5py.special_dtype(vlen=np.dtype('float64'))
            for i in np.array(['spike_times_Finf_AUC', 'ISI_Finf_AUC', 'Freq_Finf_AUC']):
                f['analysis'].require_dataset(i, shape=(I_rel2.shape[1],), dtype=dtyp, compression="gzip",
                                              compression_opts=9)
            f['analysis'].require_dataset('F_inf_Finf_AUC', shape=(I_rel2.shape[1],), dtype='float64')
            f['analysis'].require_dataset('F_null_Finf_AUC', shape=(I_rel2.shape[1],), dtype='float64')
            for stim in range(I_rel2.shape[1]):
                # spike peak detection
                ap_all, ap_prop = scipy.signal.find_peaks(V_m_AUC[stim, stim_start:], min_spike_height,
                                                          prominence=prominence, distance=1 * 1 / dt)
                f['analysis']['spike_times_Finf_AUC'][stim] = ap_all * dt
                f['analysis']['ISI_Finf_AUC'][stim] = np.array(
                    [x - f['analysis']['spike_times_Finf_AUC'][stim][i - 1] if i else None for i, x in
                     enumerate(f['analysis']['spike_times_Finf_AUC'][stim])][1:])  # msec
                f['analysis']['Freq_Finf_AUC'][stim] = 1 / (f['analysis']['ISI_Finf_AUC'][stim] / 1000)
                if f['analysis']['Freq_Finf_AUC'][stim].size != 0:
                    last_second_spikes = f['analysis']['spike_times_Finf_AUC'][stim] > np.int(
                        f['data'].attrs['stim_time'] * dt - (500 + f['data'].attrs['initial_period']))
                    if np.any(last_second_spikes == True):
                        last_second_spikes_1 = np.delete(last_second_spikes, 0)  # remove first spike because it has no ISI/freq
                        f['analysis']['F_inf_Finf_AUC'][stim] = np.mean(
                            f['analysis']['Freq_Finf_AUC'][stim][last_second_spikes_1])
                    else:
                        f['analysis']['F_null_Finf_AUC'][stim] = 0
                        f['analysis']['F_inf_Finf_AUC'][stim] = 0
            f['analysis'].require_dataset('AUC_rel_acc', shape=(1,), dtype='float64', compression="gzip",
                                          compression_opts=9)
            f['analysis']['AUC_rel_acc'][:] = np.trapz(f['analysis']['F_inf_Finf_AUC'][:],
                                                       I_rel2[:] * 1000, dI * 1000)  #
        except:
            f['analysis'].require_dataset('AUC', shape=(1,), dtype='float64', compression="gzip",
                                          compression_opts=9)
            f['analysis']['AUC'][:] = 0

    # ramp protocol #####################
    # setup ramp current
    ramp_len = np.int(sec *1000 *1/dt)
    stim_time = ramp_len *2
    I_amp = np.array([0, -0.000001])
    I_amp = np.reshape(I_amp, (1,I_amp.shape[0]))
    I_step = np.zeros((stim_time + np.int(initial_period * 1 / dt), 1))@ I_amp
    I_step[np.int(initial_period * 1 / dt):, :] = np.ones((stim_time, 1)) @ I_amp
    stim_num_step = I_step.shape[1]
    start = np.int(initial_period * 1 / dt)
    I_step[start:np.int(start+ramp_len),0] = np.linspace(0, high, ramp_len)
    I_step[np.int(start + ramp_len):np.int(start + ramp_len*2),0] = np.linspace(high, 0, ramp_len)
    stim_time_step = I_step.shape[0]

    # simulate response to ramp
    V_m_step = Cb_stellate_Kv(-70 * np.ones(stim_num_step), g_effect, E2, I_step, dt, currents_included2,
                              stim_time_step, stim_num_step, C, shift2, scale2, b_param_effect, slope_shift2,
                              np.zeros((len(b_param), stim_num_step)),
                              np.zeros((len(currents_included), stim_num_step)), ind_dict)


    # spike peak detection
    ap_all, ap_prop = scipy.signal.find_peaks(V_m_step[0, stim_start:], min_spike_height,
                                                          prominence=prominence, distance=1 * 1 / dt)
    # save ramp firing properties
    with h5py.File(fname, "a") as f:
        f['data'].create_dataset('V_m_ramp', data=V_m_step, dtype='float64', compression="gzip", compression_opts=9)
        if ap_all.shape[0] != 0:
            if ap_all[0] < stim_start + ramp_len:
                f['analysis'].create_dataset('ramp_I_up', data =I_step[stim_start+ap_all[0], 0] *1000,  dtype='float64')
            else:
                f['analysis'].create_dataset('ramp_I_up', data =np.NaN,  dtype='float64')
            if ap_all[-1] > stim_start + ramp_len:
                f['analysis'].create_dataset('ramp_I_down', data =I_step[stim_start+ap_all[-1], 0]*1000,  dtype='float64')
            else:
                f['analysis'].create_dataset('ramp_I_down', data =np.NaN,  dtype='float64')
            f['analysis'].create_dataset('hysteresis', data =f['analysis']['ramp_I_down'][()] - f['analysis']['ramp_I_up'][()],  dtype='float64')
        else:# if no spikes in response to ramp
            f['analysis'].create_dataset('ramp_I_up', data =np.NaN,  dtype='float64')
            f['analysis'].create_dataset('ramp_I_down', data =np.NaN,  dtype='float64')
            f['analysis'].create_dataset('hysteresis', data =np.NaN,  dtype='float64')
    if f.__bool__():
        f.close()
    gc.collect()


def SA_Cb_stellate_Kv(V_init, g, E, I_in, dt, currents_included, stim_time, stim_num, C, shift, scale, b_param,
                      slope_shift, gating, current, prominence, desired_AUC_width, folder, high, low, number_steps,
                      initial_period, sec, lin_array, log_array, alt_types, alt_ind, alt):
    """ Sensitivity analysis for Cb Stellate Kv model

            Parameters
            ----------
            V_init : float
                Initial membrane potential
            g : dict
                maximal conductance for currents
            E : dict
                Reversal potentials for currents
            I_in : array
                Current stimulus input to model
            dt : float
                fixed time step
            currents_included : dict
                Boolean to include current in simulation or not
            stim_time :
                length of stimulus ()
            stim_num : int
                number of different stimulus currents
            C : float
                membrane capacitance
            shift : dict
                for each gating parameter
            scale : dict
                for each gating parameter
            b_param : dict
                Boltzmann parameter values for each gating parameter
            slope_shift : dict
                for each gating parameter
            gating : array
                array of gating values over time
            current : array
                array of current values over time
            prominence: float
                required spike prominence
            desired_AUC_width: float
                width of AUC from rheobase to rheobase + desired_AUC_width
            folder : str
                folder to save hdf5 in
            high : float
                upper current step magnitude
            low : float
                lowest current step magnitude
            number_steps : int
                number of current steps between low and high
            initial_period: float
                length  of  I = 0 before current step
            sec : float
                length of current step in seconds
            lin_array : array
                array of linear shifts from -10 tp 10
            log_array : array
                array of log2 values from -1 to 1
            alt_types : array
                array of arrays of strings of [variable, alteration type]
            alt_ind : int
                index for alt_type
            alt :
                index for lin_array of log_array


            Returns
            -------
            saves hdf5 file to folder
            """
    # setup for simulation
    g_multi = copy.deepcopy(g)
    E_multi = copy.deepcopy(E)
    b_param_multi = copy.deepcopy(b_param)
    shift_multi = copy.deepcopy(shift)
    scale_multi = copy.deepcopy(scale)
    slope_shift_multi = copy.deepcopy(slope_shift)
    currents_included_multi = copy.deepcopy(currents_included)
    gating2 = copy.deepcopy(gating)
    current2 = copy.deepcopy(current)

    g2 = Dict.empty(key_type=types.unicode_type, value_type=types.float64, )
    for k, v in g_multi.items():
        g2[k] = v
    b_param2 = Dict.empty(key_type=types.unicode_type, value_type=types.float64[:], )
    for k, v in b_param_multi.items():
        b_param2[k] = np.array(v)
    shift2 = Dict.empty(key_type=types.unicode_type, value_type=types.float64, )
    for k, v in shift_multi.items():
        shift2[k] = v
    scale2 = Dict.empty(key_type=types.unicode_type, value_type=types.float64, )
    for k, v in scale_multi.items():
        scale2[k] = v
    slope_shift2 = Dict.empty(key_type=types.unicode_type, value_type=types.float64, )
    for k, v in slope_shift_multi.items():
        slope_shift2[k] = v
    E2 = Dict.empty(key_type=types.unicode_type, value_type=types.float64, )
    for k, v in E_multi.items():
        E2[k] = v
    currents_included2 = Dict.empty(key_type=types.unicode_type, value_type=types.boolean, )
    for k, v in currents_included_multi.items():
        currents_included2[k] = v
    ind_dict = Dict.empty(key_type=types.unicode_type, value_type=types.int64, )
    i = 0
    for var in np.array(['m', 'h', 'n', 'n_A', 'h_A', 'm_T', 'h_T', 's', 'u', 's_mut', 'u_mut']):
        ind_dict[var] = i
        i += 1
    i = 0
    for var in np.array(['Na', 'Kd', 'A', 'Kv', 'Kv_mut', 'T', 'Leak']):
        ind_dict[var] = i
        i += 1

    # gating alteration ##########################################################################
    if alt_types[alt_ind, 1] == 'shift':
        shift2[alt_types[alt_ind, 0]] = lin_array[alt]
        fname = os.path.join(folder,
                             "{}_{}_{}.hdf5".format(alt_types[alt_ind, 1], alt_types[alt_ind, 0], lin_array[alt]))

    elif alt_types[alt_ind, 1] == 'slope':
        b_param2[alt_types[alt_ind, 0]][1] = b_param2[alt_types[alt_ind, 0]][1] * log_array[alt]
        fname = os.path.join(folder, "{}_{}_{}.hdf5".format(alt_types[alt_ind, 1], alt_types[alt_ind, 0],
                                                            np.int(np.around(log_array[alt], 2) * 100)))
        if b_param2[alt_types[alt_ind, 0]][2] != 1:
            # adjust slope_shift to account for slope not changing (rotating) around 0.5
            V_test = np.arange(-150, 100, 0.001)
            if b_param2[alt_types[alt_ind, 0]].shape[0] == 4:
                steady_state = ((1 - b_param2[alt_types[alt_ind, 0]][3]) / (
                        1 + np.exp(
                    (V_test - b_param2[alt_types[alt_ind, 0]][0]) / b_param2[alt_types[alt_ind, 0]][1])) +
                                b_param2[alt_types[alt_ind, 0]][3]) ** b_param2[alt_types[alt_ind, 0]][2]
                orig = ((1 - b_param[alt_types[alt_ind, 0]][3]) / (
                        1 + np.exp((V_test - b_param[alt_types[alt_ind, 0]][0]) / b_param[alt_types[alt_ind, 0]][1])) +
                        b_param[alt_types[alt_ind, 0]][3]) ** b_param[alt_types[alt_ind, 0]][2]
            else:
                steady_state = (1 / (1 + np.exp(
                    (V_test - b_param2[alt_types[alt_ind, 0]][0]) / b_param2[alt_types[alt_ind, 0]][1]))) ** \
                               b_param2[alt_types[alt_ind, 0]][2]
                orig = (1 / (1 + np.exp(
                    (V_test - b_param[alt_types[alt_ind, 0]][0]) / b_param[alt_types[alt_ind, 0]][1]))) ** \
                       b_param[alt_types[alt_ind, 0]][2]
            orig_V_half = V_test[(np.abs(orig - 0.5)).argmin()]
            V_half_new = V_test[(np.abs(steady_state - 0.5)).argmin()]
            slope_shift2[alt_types[alt_ind, 0]] = orig_V_half - V_half_new

    elif alt_types[alt_ind, 1] == 'g':
        g2[alt_types[alt_ind, 0]] = g2[alt_types[alt_ind, 0]] * log_array[alt]
        fname = os.path.join(folder, "{}_{}_{}.hdf5".format(alt_types[alt_ind, 1], alt_types[alt_ind, 0],
                                                            np.int(np.around(log_array[alt], 2) * 100)))
    ###########################################################################

    # initial simulation for current range from low to high
    V_m = Cb_stellate_Kv(V_init * np.ones((stim_num)), g2, E2, I_in, dt, currents_included2, stim_time, stim_num, C,
                         shift2, scale2, b_param2, slope_shift2, gating2, current2, ind_dict)
    stim_start = np.int(initial_period * 1 /dt)
    min_spike_height = V_m[0, stim_start] + prominence

    # create hdf5 file and save data and metadata
    with h5py.File(fname, "a") as f:
        data = f.create_group("data")
        data.create_dataset('V_m', data=V_m, dtype='float64', compression="gzip", compression_opts=9)
        metadata = {'I_high': high, 'I_low': low, 'stim_num': number_steps, 'stim_time': stim_time, 'sec': sec,
                    'dt': dt, 'initial_period': initial_period, 'g': json.dumps(dict(g2)), 'E': json.dumps(dict(E2)),
                    'C': C, 'V_init': V_init, 'ind_dict': json.dumps(dict(ind_dict)),
                    'b_param': json.dumps(dict(b_param2), cls=NumpyEncoder),
                    'currents': json.dumps(dict(currents_included2)), 'shift': json.dumps(dict(shift2)),
                    'var_change': str(alt_types[alt_ind, 1]), 'var': str(alt_types[alt_ind, 0]),
                    'scale': json.dumps(dict(scale2)), 'slope_shift': json.dumps(dict(slope_shift2)),
                    'prominence': prominence, 'desired_AUC_width': desired_AUC_width,
                    'alteration_info': str(alt_types[alt_ind, :]), 'alt_index': str(alt),
                    'min_spike_height': min_spike_height,}
        data.attrs.update(metadata)
        if alt_types[alt_ind, 1] == 'shift':
            meta2 = {'alteration': lin_array[alt]}
        else:
            meta2 = {'alteration': log_array[alt]}
        data.attrs.update(meta2)

    # firing frequency analysis and rheobase
    with h5py.File(fname, "r+") as f:
        I_mag = np.arange(low, high, (high - low) / stim_num)
        analysis = f.create_group("analysis")
        dtyp = h5py.special_dtype(vlen=np.dtype('float64'))
        for i in np.array(['spike_times', 'spike_times_rheo', 'ISI', 'Freq']):  # analysis_names:
            analysis.create_dataset(i, (I_mag.shape[0],), dtype=dtyp, compression="gzip", compression_opts=9)
        analysis.create_dataset('F_inf', (I_mag.shape[0],), dtype='float64')
        analysis.create_dataset('F_null', (I_mag.shape[0],), dtype='float64')
        analysis.create_dataset('AP', (I_mag.shape[0],), dtype='bool', compression="gzip", compression_opts=9)
        analysis.create_dataset('AP_rheo', (I_mag.shape[0],), dtype='bool', compression="gzip", compression_opts=9)
        analysis.create_dataset('rheobase', (1,), dtype='float64', compression="gzip", compression_opts=9)
        try:
            for stim in range(I_mag.shape[0]):
                # spike detection
                ap_all, ap_prop = scipy.signal.find_peaks(V_m[stim, stim_start:], min_spike_height,
                                                          prominence=prominence, distance=1 * 1 / dt)
                analysis['spike_times'][stim] = ap_all * dt
                if analysis['spike_times'][stim].size == 0:
                    analysis['AP'][stim] = False
                else:
                    analysis['AP'][stim] = True
                analysis['ISI'][stim] = np.array([x - analysis['spike_times'][stim][i - 1] if i else None for i, x in
                                                  enumerate(analysis['spike_times'][stim])][1:])  # msec
                analysis['Freq'][stim] = 1 / (analysis['ISI'][stim] / 1000)

                if analysis['Freq'][stim].size != 0:
                    analysis['F_null'][stim] = analysis['Freq'][stim][0]
                    if np.argwhere(ap_all >= 1000 * 1 / dt).shape[0] != 0:  # if first spike is after 1 ms
                        half_second_spikes = np.logical_and(analysis['spike_times'][stim] < np.int(
                            ap_all[np.argwhere(ap_all >= 1000 * 1 / dt)[0][0]] * dt + 500),
                                                            analysis['spike_times'][stim] > np.int(
                                                                ap_all[np.argwhere(ap_all >= 1000 * 1 / dt)[0][0]] * dt))  # spikes in half second window
                        if np.sum(half_second_spikes == True) > 1:
                            half_second_spikes_1 = np.delete(half_second_spikes,
                                                             np.argwhere(half_second_spikes == True)[0][0])
                            analysis['F_inf'][stim] = np.mean(analysis['Freq'][stim][half_second_spikes_1])
                        else:
                            analysis['F_inf'][stim] = 0
                    else:
                        analysis['F_inf'][stim] = 0
                else: # if no ISI detected
                    analysis['F_null'][stim] = 0
                    analysis['F_inf'][stim] = 0
        except:
            print(alt_types[alt_ind, 0], alt_types[alt_ind, 1], alt, 'Freq analysis unsuccessful')

        # rheobase
        # find where the first AP occurs
        if np.any(analysis['AP'][:] == True):
            # setup current range to be between the current step before and with first AP
            I_first_spike = I_mag[np.argwhere(analysis['AP'][:] == True)[0][0]]
            I_before_spike = I_mag[np.argwhere(analysis['AP'][:] == True)[0][0] - 1]
            analysis.create_dataset('I_first_spike', data=I_first_spike, dtype='float64')
            analysis.create_dataset('I_before_spike', data=I_before_spike, dtype='float64')
            number_steps_rheo = 100
            stim_time, I_in_rheo, stim_num_rheo, V_m_rheo = stimulus_init(I_before_spike, I_first_spike,
                                                                               number_steps_rheo, initial_period, dt,
                                                                               sec)

            # simulate response to this current range and save
            V_m_rheo = Cb_stellate_Kv(V_init * np.ones((stim_num_rheo)), g2, E2, I_in_rheo, dt, currents_included2,
                                      stim_time, stim_num_rheo, C, shift2, scale2, b_param2, slope_shift2,
                                      np.zeros((len(b_param), stim_num_rheo)),
                                      np.zeros((len(currents_included), stim_num_rheo)), ind_dict)

            # run AP detection
            try:
                for stim in range(I_in_rheo.shape[1]):
                    ap_all, ap_prop = scipy.signal.find_peaks(V_m_rheo[stim, stim_start:], min_spike_height,
                                                              prominence=prominence, distance=1 * 1 / dt)
                    analysis['spike_times_rheo'][stim] = ap_all * dt
                    if analysis['spike_times_rheo'][stim].size == 0:
                        analysis['AP_rheo'][stim] = False
                    else:
                        analysis['AP_rheo'][stim] = True
                # find rheobase as smallest current step to elicit AP and save
                analysis['rheobase'][:] = I_in_rheo[-1, np.argwhere(analysis['AP_rheo'][:] == True)[0][0]] * 1000
            except:
                print(alt_types[alt_ind, 0], alt_types[alt_ind, 1], alt, 'RHEOBASE NO AP')

    # AUC
    # find rheobase of steady state firing (F_inf)
    with h5py.File(fname, "a") as f:
        try:
            # setup current range to be in current step between no and start of steady state firing
            F_inf_start_ind = np.argwhere(f['analysis']['F_inf'][:] != 0)[0][0]
            dI = (high - low) / stim_num
            I_start = I_mag[F_inf_start_ind - 1]
            I_end = I_mag[F_inf_start_ind]
            I_rel = np.arange(I_start, I_end + (I_end - I_start) / 100, (I_end - I_start) / 100)
            I_rel = np.reshape(I_rel, (1, I_rel.shape[0]))
            I_in = np.zeros((stim_time + np.int(initial_period * 1 / dt), 1)) @ I_rel
            I_in[np.int(initial_period * 1 / dt):, :] = np.ones((stim_time, 1)) @ I_rel

            # simulate response to this current step range
            V_m_inf_rheo = Cb_stellate_Kv(V_init * np.ones(stim_num), g2, E2, I_in, dt, currents_included2, stim_time,
                                          stim_num, C, shift2, scale2, b_param2, slope_shift2,
                                          np.zeros((len(b_param), stim_num)),
                                          np.zeros((len(currents_included), stim_num)), ind_dict)

            # save response, analyse and save
            for i in np.array(['spike_times_Finf', 'ISI_Finf', 'Freq_Finf']):
                f['analysis'].require_dataset(i, shape=(I_rel.shape[1],), dtype=dtyp, compression="gzip",
                                              compression_opts=9)
            f['analysis'].require_dataset('F_inf_Finf', shape=(I_rel.shape[1],), dtype='float64')
            f['analysis'].require_dataset('F_null_Finf', shape=(I_rel.shape[1],), dtype='float64')
            for stim in range(I_rel.shape[1]):
                # spike peak detection
                ap_all, ap_prop = scipy.signal.find_peaks(V_m_inf_rheo[stim, stim_start:], min_spike_height,
                                                          prominence=prominence, distance=1 * 1 / dt)
                f['analysis']['spike_times_Finf'][stim] = ap_all * dt
                f['analysis']['ISI_Finf'][stim] = np.array(
                    [x - f['analysis']['spike_times_Finf'][stim][i - 1] if i else None for i, x in
                     enumerate(f['analysis']['spike_times_Finf'][stim])][1:])  # msec
                f['analysis']['Freq_Finf'][stim] = 1 / (f['analysis']['ISI_Finf'][stim] / 1000)
                if f['analysis']['Freq_Finf'][stim].size != 0:
                    if np.argwhere(ap_all >= 1000 * 1 / dt).shape[0] != 0:  # if first spike is after 1 ms
                        half_second_spikes = np.logical_and(f['analysis']['spike_times_Finf'][stim] < np.int(
                            ap_all[np.argwhere(ap_all >= 1000 * 1 / dt)[0][0]] * dt + 500),
                                                            f['analysis']['spike_times_Finf'][stim] > np.int(
                                                                ap_all[np.argwhere(ap_all >= 1000 * 1 / dt)[0][0]] * dt))  # spikes in half second window
                        if np.sum(half_second_spikes == True) > 1:
                            half_second_spikes_1 = np.delete(half_second_spikes,
                                                             np.argwhere(half_second_spikes == True)[0][0])  # remove first spike because it has no ISI/freq
                            f['analysis']['F_inf_Finf'][stim] = np.mean( f['analysis']['Freq_Finf'][stim][half_second_spikes_1])
                        else:
                            f['analysis']['F_inf_Finf'][stim] = 0
                    else:
                        f['analysis']['F_null_Finf'][stim] = 0
                        f['analysis']['F_inf_Finf'][stim] = 0

                else:
                    f['analysis']['F_null_Finf'][stim] = 0
                    f['analysis']['F_inf_Finf'][stim] = 0

            # find AUC relative to F_inf rheobase
            # setup current inputs with current range around F_inf rheobase
            I_first_inf = I_rel[0, np.argwhere(f['analysis']['F_inf_Finf'][:] != 0)[0][0]]
            I_start = I_first_inf - 0.00005
            I_end = I_first_inf + 0.2 * f['data'].attrs['I_high'][()]
            I_rel2 = np.arange(I_start, I_end + (I_end - I_start) / 100, (I_end - I_start) / 100)
            I_rel2 = np.reshape(I_rel2, (1, I_rel2.shape[0]))
            I_in2 = np.zeros((stim_time + np.int(initial_period * 1 / dt), 1)) @ I_rel2
            I_in2[np.int(initial_period * 1 / dt):, :] = np.ones((stim_time, 1)) @ I_rel2
            stim_num = I_in2.shape[1]

            # simulate response to this current step range
            V_m_AUC = Cb_stellate_Kv(V_init * np.ones(stim_num), g2, E2, I_in2, dt, currents_included2, stim_time,
                                     stim_num, C, shift2, scale2, b_param2, slope_shift2,
                                     np.zeros((len(b_param), stim_num)), np.zeros((len(currents_included), stim_num)),
                                     ind_dict)

            # save data and analysis for this current step range including AUC
            f['data'].require_dataset('V_m_AUC', shape=V_m_AUC.shape, dtype='float64')
            f['data']['V_m_AUC'][:] = V_m_AUC
            dtyp = h5py.special_dtype(vlen=np.dtype('float64'))
            for i in np.array(['spike_times_Finf_AUC', 'ISI_Finf_AUC', 'Freq_Finf_AUC']):  # analysis_names:
                f['analysis'].require_dataset(i, shape=(I_rel2.shape[1],), dtype=dtyp, compression="gzip", compression_opts=9)
            f['analysis'].require_dataset('F_inf_Finf_AUC', shape=(I_rel2.shape[1],), dtype='float64')
            f['analysis'].require_dataset('F_null_Finf_AUC', shape=(I_rel2.shape[1],), dtype='float64')
            for stim in range(I_rel2.shape[1]):
                # spike peak detection
                ap_all, ap_prop = scipy.signal.find_peaks(V_m_AUC[stim, stim_start:], min_spike_height,
                                                          prominence=prominence, distance=1 * 1 / dt)
                f['analysis']['spike_times_Finf_AUC'][stim] = ap_all * dt
                f['analysis']['ISI_Finf_AUC'][stim] = np.array(
                    [x - f['analysis']['spike_times_Finf_AUC'][stim][i - 1] if i else None for i, x in
                     enumerate(f['analysis']['spike_times_Finf_AUC'][stim])][1:])  # msec
                f['analysis']['Freq_Finf_AUC'][stim] = 1 / (f['analysis']['ISI_Finf_AUC'][stim] / 1000)
                if f['analysis']['Freq_Finf_AUC'][stim].size != 0:
                    if np.argwhere(ap_all >= 1000 * 1 / dt).shape[0] != 0:  # if first spike is after 1 ms
                        half_second_spikes = np.logical_and(f['analysis']['spike_times_Finf_AUC'][stim] < np.int(
                            ap_all[np.argwhere(ap_all >= 1000 * 1 / dt)[0][0]] * dt + 500),
                                                            f['analysis']['spike_times_Finf_AUC'][stim] > np.int(
                                                                ap_all[np.argwhere(ap_all >= 1000 * 1 / dt)[0][0]] * dt))  # spikes in half second window
                        if np.sum(half_second_spikes == True) > 1:
                            half_second_spikes_1 = np.delete(half_second_spikes,
                                                             np.argwhere(half_second_spikes == True)[0][0])  # remove first spike because it has no ISI/freq
                            f['analysis']['F_inf_Finf_AUC'][stim] = np.mean(
                                f['analysis']['Freq_Finf_AUC'][stim][half_second_spikes_1])
                        else:
                            f['analysis']['F_inf_Finf_AUC'][stim] = 0
                    else:
                        f['analysis']['F_null_Finf_AUC'][stim] = 0
                        f['analysis']['F_inf_Finf_AUC'][stim] = 0
                else:
                    f['analysis']['F_null_Finf_AUC'][stim] = 0
                    f['analysis']['F_inf_Finf_AUC'][stim] = 0
            f['analysis'].require_dataset('AUC', shape=(1,), dtype='float64', compression="gzip", compression_opts=9)
            f['analysis']['AUC'][:] = np.trapz(f['analysis']['F_inf_Finf_AUC'][:], I_rel2[:] * 1000, dI * 1000)
        except:
            f['analysis'].require_dataset('AUC', shape=(1,), dtype='float64', compression="gzip",  compression_opts=9)
            f['analysis']['AUC'][:] = 0


    if f.__bool__():
        f.close()
    gc.collect()
```
