## ExtendedData1 for "Loss or Gain of Function? Effects of Ion Channel Mutations on Neuronal Firing Depend on the Cell Type": STN_fxns.py

```
"""
Functions for STN model


"""
__author__ = "Nils A. Koch"
__copyright__ = 'Copyright 2022, Nils A. Koch'
__license__ = "MIT"

import scipy
import json
import copy
import os
import gc
import h5py
import numpy as np
from numba import types, njit
from numba.typed import Dict
from scipy import signal
from Code.Functions.Utility_fxns import stimulus_init, NumpyEncoder


@njit
def STN(V, g, E, I_in, dt, currents_included, stim_time, stim_num, C, shift, scale, b_param, slope_shift,
                  gating, current, ind_dict, vol):
    """ Simulate current and Vm for  STN model from Otsuka et al 2004 (https://dx.doi.org/10.1152/jn.00508.2003)
        for I_in

        Parameters
        ----------
        V : float
            Initial membrane potentia
        g : dict
            maximal conductance for currents
        E : dict
            Reversal potentials for currents
        I_in : array
            Current stimulus input to model
        dt : float
            fixed time step
        currents_included : dict
            Boolean to include current in simulation or not
        stim_time :
            length of stimulus ()
        stim_num : int
            number of different stimulus currents
        C : float
            membrane capacitance
        shift : dict
            for each gating parameter
        scale : dict
            for each gating parameter
        b_param : dict
            Boltzmann parameter values for each gating parameter
        slope_shift : dict
            for each gating parameter
        gating : array
            array of gating values over time
        current : array
            array of current values over time
        ind_dict: dict
            Dictionary of indices in arrays for gating variables
        vol : float
            cell volume

        Returns
        -------
        V_m : array
            Simulated membrane potential in response to I_in
        """
    # initialize V_m, Ca_conc and E_Ca
    V_m = np.zeros((stim_num, stim_time))
    Ca_conc = np.ones((stim_num)) * 5e-3
    E_Ca = 0.013319502 * np.log(2000 / Ca_conc) * 1000 * np.ones((stim_num))  # mV

    # initialize gating
    # Na activation
    m_inf = (1 / (1 + np.exp((V - shift["m"] - slope_shift["m"] - b_param["m"][0]) / b_param["m"][1]))) ** b_param["m"][2]
    gating[ind_dict['m'], :] = np.ones((stim_num), dtype=np.dtype('float64')) * m_inf

    # Na inactivation
    h_inf = ((1 - b_param["h"][3]) / (1 + np.exp((V - shift["h"] - slope_shift["h"] - b_param["h"][0]) / b_param["h"][1])) + b_param["h"][3]) \
            ** b_param["h"][2]
    gating[ind_dict['h'], :] = np.ones((stim_num), dtype=np.dtype('float64')) * h_inf

    # K activation
    n_inf = (1 / (1 + np.exp((V - shift["n"] - slope_shift["n"] - b_param["n"][0]) / b_param["n"][1]))) ** \
            b_param["n"][2]
    gating[ind_dict['n'], :] = np.ones((stim_num), dtype=np.dtype('float64')) * n_inf

    # T-type Ca activation
    p_inf = (1 / (1 + np.exp((V - shift["p"] - slope_shift["p"] - b_param["p"][0]) / b_param["p"][1]))) ** \
            b_param["p"][2]
    gating[ind_dict['p'], :] = np.ones((stim_num), dtype=np.dtype('float64')) * p_inf

    # T-type Ca inactivation
    q_inf = (1 / (1 + np.exp((V - shift["q"] - slope_shift["q"] - b_param["q"][0]) / b_param["q"][1]))) ** \
            b_param["q"][2]
    gating[ind_dict['q'], :] = np.ones((stim_num), dtype=np.dtype('float64')) * q_inf

    # Ca dependent K activation
    r_inf = (1 / (1 + np.exp((Ca_conc - shift["r"] - slope_shift["r"] - b_param["r"][0]) / b_param["r"][1]))) ** \
            b_param["r"][2]
    gating[ind_dict['r'], :] = np.ones((stim_num), dtype=np.dtype('float64')) * r_inf

    # A-type activation
    a_inf = (1 / (1 + np.exp((V - shift["a"] - slope_shift["a"] - b_param["a"][0]) / b_param["a"][1]))) ** \
            b_param["a"][2]
    gating[ind_dict['a'], :] = np.ones((stim_num), dtype=np.dtype('float64')) * a_inf

    # A-type inactivation
    b_inf = (1 / (1 + np.exp((V - shift["b"] - slope_shift["b"] - b_param["b"][0]) / b_param["b"][1]))) ** \
            b_param["b"][2]
    gating[ind_dict['b'], :] = np.ones((stim_num), dtype=np.dtype('float64')) * b_inf

    # mutant A-type activation
    a_mut_inf = (1 / (1 + np.exp((V - shift["a_mut"] - slope_shift["a_mut"] - b_param["a_mut"][0]) / b_param["a_mut"][1]))) ** \
            b_param["a_mut"][2]
    gating[ind_dict['a_mut'], :] = np.ones((stim_num), dtype=np.dtype('float64')) * a_mut_inf

    # mutant A-type inactivation
    b_mut_inf = (1 / (1 + np.exp((V - shift["b_mut"] - slope_shift["b_mut"] - b_param["b_mut"][0]) / b_param["b_mut"][1]))) ** \
            b_param["b_mut"][2]
    gating[ind_dict['b_mut'], :] = np.ones((stim_num), dtype=np.dtype('float64')) * b_mut_inf

    # L-type Ca activation
    c_inf = (1 / (1 + np.exp((V - shift["c"] - slope_shift["c"] - b_param["c"][0]) / b_param["c"][1]))) ** \
            b_param["c"][2]
    gating[ind_dict['c'], :] = np.ones((stim_num), dtype=np.dtype('float64')) * c_inf

    # L-type Ca inactivation 1
    d1_inf = (1 / (1 + np.exp((V - shift["d1"] - slope_shift["d1"] - b_param["d1"][0]) / b_param["d1"][1]))) ** \
             b_param["d1"][2]
    gating[ind_dict['d1'], :] = np.ones((stim_num), dtype=np.dtype('float64')) * d1_inf

    # L-type Ca inactivation 2
    d2_inf = (1 / (1 + np.exp((Ca_conc - shift["d2"] - slope_shift["d2"] - b_param["d2"][0]) / b_param["d2"][1]))) ** \
             b_param["d2"][2]
    gating[ind_dict['d2'], :] = np.ones((stim_num), dtype=np.dtype('float64')) * d2_inf

    # initialize gating and V_m
    current[ind_dict['Na'], :] = np.zeros((stim_num), dtype=np.dtype('float64'))
    current[ind_dict['Kd'], :] = np.zeros((stim_num), dtype=np.dtype('float64'))
    current[ind_dict['A'], :] = np.zeros((stim_num), dtype=np.dtype('float64'))
    current[ind_dict['A_mut'], :] = np.zeros((stim_num), dtype=np.dtype('float64'))
    current[ind_dict['Leak'], :] = np.zeros((stim_num), dtype=np.dtype('float64'))
    current[ind_dict['L'], :] = np.zeros((stim_num), dtype=np.dtype('float64'))
    current[ind_dict['T'], :] = np.zeros((stim_num), dtype=np.dtype('float64'))
    V_m[:, 0] = V
    gating[ind_dict['Ca_conc'], :] = Ca_conc
    gating[ind_dict['E_Ca'], :] = E_Ca
    t = 1
    while t < stim_time:
        # update Ca conc and E_Ca
        Ca_conc_dt = ((-5.182134834753951e-06 / vol) * (current[ind_dict['T'], :] + current[ind_dict['L'], :])) / 1000 - (2 * Ca_conc)
        Ca_conc = Ca_conc + Ca_conc_dt * dt
        Ca_conc[Ca_conc <= 0] = 0
        E_Ca = 0.013319502 * np.log(2000 / Ca_conc) * 1000  # mV
        gating[ind_dict['Ca_conc'], :] = Ca_conc
        gating[ind_dict['E_Ca'], :] = E_Ca

        # Na activation
        m_inf = (1 / (1 + np.exp((V_m[:, t - 1] - shift["m"] - slope_shift["m"] - b_param["m"][0]) / b_param["m"][1]))) ** \
                b_param["m"][2]
        tau_m = 0.2 + 3 / (1 + np.exp((V_m[:, t - 1] - shift["m"] + 53) / 0.7))
        m_dot = (m_inf - gating[ind_dict['m'], :]) * (1 / (tau_m * scale["m"]))
        gating[ind_dict['m'], :] = gating[ind_dict['m'], :] + m_dot * dt

        # Na inactivation
        h_inf = ((1 - b_param["h"][3]) / (1 + np.exp((V_m[:, t - 1] - shift["h"] - slope_shift["h"] - b_param["h"][0])
                                                     / b_param["h"][1])) + b_param["h"][3]) ** b_param["h"][2]
        tau_h = 24.5 / (np.exp((V_m[:, t - 1] - shift["h"] + 50) / 15) + np.exp(-(V_m[:, t - 1] - shift["h"] + 50) / 16))
        h_dot = (h_inf - gating[ind_dict['h'], :]) * (1 / (tau_h * scale["h"]))
        gating[ind_dict['h'], :] = gating[ind_dict['h'], :] + h_dot * dt

        # K activation
        n_inf = (1 / (1 + np.exp((V_m[:, t - 1] - shift["n"] - slope_shift["n"] - b_param["n"][0]) / b_param["n"][1]))) ** \
                b_param["n"][2]
        tau_n = 11 / (np.exp((V_m[:, t - 1] - shift["n"] + 40) / 40) + np.exp(-(V_m[:, t - 1] - shift["n"] + 40) / 50))
        n_dot = (n_inf - gating[ind_dict['n'], :]) * (1 / (tau_n * scale["n"]))
        gating[ind_dict['n'], :] = gating[ind_dict['n'], :] + n_dot * dt

        # T-type Ca activation
        p_inf = (1 / (1 + np.exp((V_m[:, t - 1] - shift["p"] - slope_shift["p"] - b_param["p"][0]) / b_param["p"][1]))) ** \
                b_param["p"][2]
        tau_p = 5 + 0.33 / (np.exp((V_m[:, t - 1] - shift["p"] + 27) / 10) + np.exp(-(V_m[:, t - 1] - shift["p"] + 102) / 15))
        p_dot = (p_inf - gating[ind_dict['p'], :]) * (1 / (tau_p * scale["p"]))
        gating[ind_dict['p'], :] = gating[ind_dict['p'], :] + p_dot * dt

        # T-type Ca inactivation
        q_inf = (1 / (1 + np.exp((V_m[:, t - 1] - shift["q"] - slope_shift["q"] - b_param["q"][0]) / b_param["q"][1]))) ** \
                b_param["q"][2]
        tau_q = 400 / (np.exp((V_m[:, t - 1] - shift["q"] + 50) / 15) + np.exp(-(V_m[:, t - 1] - shift["q"] + 50) / 16))
        q_dot = (q_inf - gating[ind_dict['q'], :]) * (1 / (tau_q * scale["q"]))
        gating[ind_dict['q'], :] = gating[ind_dict['q'], :] + q_dot * dt

        # Ca dependent K activation
        r_inf = (1 / (1 + np.exp((Ca_conc * 1e-6 - shift["r"] - slope_shift["r"] - b_param["r"][0]) / b_param["r"][1]))) ** \
                b_param["r"][2]
        tau_r = 2
        r_dot = (r_inf - gating[ind_dict['r'], :]) * (1 / (tau_r * scale["r"]))
        gating[ind_dict['r'], :] = gating[ind_dict['r'], :] + r_dot * dt

        # A-type activation
        a_inf = (1 / (1 + np.exp((V_m[:, t - 1] - shift["a"] - slope_shift["a"] - b_param["a"][0]) / b_param["a"][1]))) ** \
                b_param["a"][2]
        tau_a = 1 + 1 / (1 + np.exp((V_m[:, t - 1] - shift["a"] + 40) / 0.5))
        a_dot = (a_inf - gating[ind_dict['a'], :]) * (1 / (tau_a * scale["a"]))
        gating[ind_dict['a'], :] = gating[ind_dict['a'], :] + a_dot * dt

        # A-type inactivation
        b_inf = (1 / (1 + np.exp((V_m[:, t - 1] - shift["b"] - slope_shift["b"] - b_param["b"][0]) / b_param["b"][1]))) ** \
                b_param["b"][2]
        tau_b = 200 / (np.exp((V_m[:, t - 1] - shift["b"] + 60) / 30) + np.exp(-(V_m[:, t - 1] - shift["b"] + 40) / 10))
        b_dot = (b_inf - gating[ind_dict['b'], :]) * (1 / (tau_b * scale["b"]))
        gating[ind_dict['b'], :] = gating[ind_dict['b'], :] + b_dot * dt

        # mutant A-type activation
        a_mut_inf = (1 / (1 + np.exp((V_m[:, t - 1] - shift["a_mut"] - slope_shift["a_mut"] - b_param["a_mut"][0]) / b_param["a_mut"][1]))) ** \
                b_param["a_mut"][2]
        tau_a_mut = 1 + 1 / (1 + np.exp((V_m[:, t - 1] - shift["a_mut"] + 40) / 0.5))
        a_mut_dot = (a_mut_inf - gating[ind_dict['a_mut'], :]) * (1 / (tau_a_mut * scale["a_mut"]))
        gating[ind_dict['a_mut'], :] = gating[ind_dict['a_mut'], :] + a_mut_dot * dt

        # mutant A-type inactivation
        b_mut_inf = (1 / (1 + np.exp((V_m[:, t - 1] - shift["b_mut"] - slope_shift["b_mut"] - b_param["b_mut"][0]) / b_param["b_mut"][1]))) ** \
                b_param["b_mut"][2]
        tau_mut_b = 200 / (np.exp((V_m[:, t - 1] - shift["b_mut"] + 60) / 30) + np.exp(-(V_m[:, t - 1] - shift["b_mut"] + 40) / 10))
        b_mut_dot = (b_mut_inf - gating[ind_dict['b_mut'], :]) * (1 / (tau_mut_b * scale["b_mut"]))
        gating[ind_dict['b_mut'], :] = gating[ind_dict['b_mut'], :] + b_mut_dot * dt

        # L-type Ca activation
        c_inf = (1 / (1 + np.exp((V_m[:, t - 1] - shift["c"] - slope_shift["c"] - b_param["c"][0]) / b_param["c"][1]))) ** \
                b_param["c"][2]
        tau_c = 45 + 10 / (np.exp((V_m[:, t - 1] - shift["c"] + 27) / 20) + np.exp(-(V_m[:, t - 1] - shift["c"] + 50) / 15))
        c_dot = (c_inf - gating[ind_dict['c'], :]) * (1 / (tau_c * scale["c"]))
        gating[ind_dict['c'], :] = gating[ind_dict['c'], :] + c_dot * dt

        # L-type Ca inactivation 1
        d1_inf = (1 / (1 + np.exp((V_m[:, t - 1] - shift["d1"] - slope_shift["d1"] - b_param["d1"][0]) / b_param["d1"][1]))) ** \
                 b_param["d1"][2]
        tau_d1 = 400 + 500 / (np.exp((V_m[:, t - 1] - shift["d1"] + 40) / 15) + np.exp(-(V_m[:, t - 1] - shift["d1"] + 20) / 20))
        d1_dot = (d1_inf - gating[ind_dict['d1'], :]) * (1 / (tau_d1 * scale["d1"]))
        gating[ind_dict['d1'], :] = gating[ind_dict['d1'], :] + d1_dot * dt

        # L-type Ca inactivation 2
        d2_inf = (1 / (1 + np.exp((Ca_conc * 1e-6 - shift["d2"] - slope_shift["d2"] - b_param["d2"][0]) / b_param["d2"][1]))) ** \
                 b_param["d2"][2]
        tau_d2 = 130  # dt
        d2_dot = (d2_inf - gating[ind_dict['d2'], :]) * (1 / (tau_d2 * scale["d2"]))
        gating[ind_dict['d2'], :] = gating[ind_dict['d2'], :] + d2_dot * dt

        # update currents
        current[ind_dict['Leak'], :] = g["Leak"] * (V_m[:, t - 1] - E["Leak"]) * currents_included["Leak"]
        current[ind_dict['Na'], :] = g["Na"] * (gating[ind_dict['m'], :] ** 3) * gating[ind_dict['h'], :] \
                                     * (V_m[:, t - 1] - E["Na"]) * currents_included["Na"]
        current[ind_dict['Kd'], :] = g["Kd"] * (gating[ind_dict['n'], :] ** 4) * (V_m[:, t - 1] - E["K"]) * \
                                     currents_included["Kd"]

        current[ind_dict['L'], :] = g["L"] * ((gating[ind_dict['c'], :]) ** 2) * gating[ind_dict['d1'],:] \
                                    * gating[ind_dict['d2'],:] * (V_m[:, t - 1] - E_Ca) * currents_included["L"]
        current[ind_dict['T'], :] = g["T"] * (gating[ind_dict['p'], :] ** 2) * gating[ind_dict['q'], :] * \
                                    (V_m[:, t - 1] - E_Ca) * currents_included["T"]
        current[ind_dict['A'], :] = g["A"] * (gating[ind_dict['a'], :] ** 2) * gating[ind_dict['b'], :] * \
                                    (V_m[:, t - 1] - E["K"]) * currents_included["A"]
        current[ind_dict['A_mut'], :] = g["A_mut"] * (gating[ind_dict['a_mut'], :] ** 2) * gating[ind_dict['b_mut'], :] * \
                                        (V_m[:, t - 1] - E["K"]) * currents_included["A_mut"]
        current[ind_dict['Ca_K'], :] = g["Ca_K"] * (gating[ind_dict['r'], :]) ** 2 * (V_m[:, t - 1] - E["K"]) * \
                                       currents_included["Ca_K"]

        # update dV/dt
        V_dot = (1 / C) * (I_in[t, :] - (current[ind_dict['Leak'], :] + current[ind_dict['Na'], :] +
                                         current[ind_dict['Kd'], :] + current[ind_dict['L'], :] +
                                         current[ind_dict['T'], :] + current[ind_dict['A'], :] +
                                         current[ind_dict['A_mut'], :] +current[ind_dict['Ca_K'], :]))
        # update V_m
        V_m[:, t] = V_m[:, t - 1] + V_dot * dt
        t += 1
    return V_m


def STN_mut(V_init, g, E, I_in, dt, currents_included, stim_time, stim_num, C, shift, scale, b_param, slope_shift,
            gating, current, prominence, desired_AUC_width, mutations, mut, folder, high, low, number_steps,
            initial_period, sec, vol):
    """ Simulate mutations for STN model

        Parameters
        ----------
        V_init : float
            Initial membrane potential
        g : dict
            maximal conductance for currents
        E : dict
            Reversal potentials for currents
        I_in : array
            Current stimulus input to model
        dt : float
            fixed time step
        currents_included : dict
            Boolean to include current in simulation or not
        stim_time :
            length of stimulus ()
        stim_num : int
            number of different stimulus currents
        C : float
            membrane capacitance
        shift : dict
            for each gating parameter
        scale : dict
            for each gating parameter
        b_param : dict
            Boltzmann parameter values for each gating parameter
        slope_shift : dict
            for each gating parameter
        gating : array
            array of gating values over time
        current : array
            array of current values over time
        prominence : float
            required spike prominence
        desired_AUC_width : float
            width of AUC from rheobase to rheobase+desired_AUC_width
        mutations : dict
            dictionary of mutation effects
        mut : str
            mutation
        folder : str
            folder to save hdf5 file in
        high : float
            upper current step magnitude
        low : float
            lowest current step magnitude
        number_steps : int
            number of current steps between low and high
        initial_period : float
            length  of  I = 0 before current step
        sec:
            length of current step in seconds
        vol : float
            cell volume

        Returns
        -------
        saves mut.hdf5 file to folder
        """
    # setup for simulation
    g_multi = copy.deepcopy(g)
    E_multi = copy.deepcopy(E)
    b_param_multi = copy.deepcopy(b_param)
    shift_multi = copy.deepcopy(shift)
    scale_multi = copy.deepcopy(scale)
    slope_shift_multi = copy.deepcopy(slope_shift)
    currents_included_multi = copy.deepcopy(currents_included)
    gating2 = copy.deepcopy(gating)
    current2 = copy.deepcopy(current)

    g_effect = Dict.empty(key_type=types.unicode_type, value_type=types.float64, )
    for k, v in g_multi.items():
        g_effect[k] = v
    b_param_effect = Dict.empty(key_type=types.unicode_type, value_type=types.float64[:], )
    for k, v in b_param_multi.items():
        b_param_effect[k] = np.array(v)
    shift2 = Dict.empty(key_type=types.unicode_type, value_type=types.float64, )
    for k, v in shift_multi.items():
        shift2[k] = v
    scale2 = Dict.empty(key_type=types.unicode_type, value_type=types.float64, )
    for k, v in scale_multi.items():
        scale2[k] = v
    slope_shift2 = Dict.empty(key_type=types.unicode_type, value_type=types.float64, )
    for k, v in slope_shift_multi.items():
        slope_shift2[k] = v
    E2 = Dict.empty(key_type=types.unicode_type, value_type=types.float64, )
    for k, v in E_multi.items():
        E2[k] = v
    currents_included2 = Dict.empty(key_type=types.unicode_type, value_type=types.boolean, )
    for k, v in currents_included_multi.items():
        currents_included2[k] = v
    ind_dict = Dict.empty(key_type=types.unicode_type, value_type=types.int64, )
    i = 0
    for var in np.array(
            ['m', 'h', 'n', 'a', 'b','a_mut', 'b_mut', 'c', 'd1', 'd2', 'p', 'q', 'r', 'Ca_conc', 'E_Ca']):
        ind_dict[var] = i
        i += 1
    i = 0
    for var in np.array(['Na', 'Kd', 'A', 'A_mut', 'L', 'T', 'Leak', 'Ca_K']):
        ind_dict[var] = i
        i += 1

    #mutation effect
    g_effect['A_mut'] = g['A_mut'] * mutations[mut]['g_ratio']
    b_param_effect['a_mut'][0] = b_param_effect['a_mut'][0] + mutations[mut]['activation_Vhalf_diff']
    b_param_effect['a_mut'][1] = b_param_effect['a_mut'][1] * mutations[mut]['activation_k_ratio']

    # initial simulation for current range from low to high
    V_m = STN(V_init * np.ones((stim_num)), g_effect, E2, I_in, dt, currents_included2, stim_time, stim_num, C, shift2,
              scale2, b_param_effect, slope_shift2, gating2, current2, ind_dict, vol)
    stim_start = np.int(initial_period * 1 / dt)
    min_spike_height = V_m[0, stim_start] + prominence

    # create hdf5 file for the mutation and save data and metadata
    fname = os.path.join(folder, "{}.hdf5".format(mut.replace(" ", "_")))
    with h5py.File(fname, "a") as f:
        data = f.create_group("data")
        data.create_dataset('V_m', data=V_m, dtype='float64', compression="gzip", compression_opts=9)
        metadata = {'I_high': high, 'I_low': low, 'stim_num': number_steps, 'stim_time': stim_time, 'sec': sec, 'dt': dt,
                    'initial_period': initial_period, 'g': json.dumps(dict(g_effect)), 'E': json.dumps(dict(E2)), 'C': C,
                    'V_init': V_init, 'vol': vol, 'ind_dict': json.dumps(dict(ind_dict)),
                    'b_param': json.dumps(dict(b_param_effect), cls=NumpyEncoder),
                    'currents': json.dumps(dict(currents_included2)), 'shift': json.dumps(dict(shift2)),
                    'scale': json.dumps(dict(scale2)), 'slope_shift': json.dumps(dict(slope_shift2)),
                    'prominence': prominence, 'desired_AUC_width': desired_AUC_width, 'mut': str(mut),
                    'min_spike_height': min_spike_height,}
        data.attrs.update(metadata)

    # firing frequency analysis and rheobase
    with h5py.File(fname, "r+") as f:
        I_mag = np.arange(low, high, (high - low) / stim_num)
        analysis = f.create_group("analysis")
        dtyp = h5py.special_dtype(vlen=np.dtype('float64'))
        for i in np.array(['spike_times', 'ISI', 'Freq','amplitude',]):
            analysis.create_dataset(i, (I_mag.shape[0],), dtype=dtyp, compression="gzip", compression_opts=9)
        analysis.create_dataset('F_inf', (I_mag.shape[0],), dtype='float64')
        analysis.create_dataset('F_null', (I_mag.shape[0],), dtype='float64')
        analysis.create_dataset('AP', (I_mag.shape[0],), dtype='bool', compression="gzip", compression_opts=9)
        analysis.create_dataset('rheobase', (1,), dtype='float64', compression="gzip", compression_opts=9)
        for stim in range(I_mag.shape[0]):
            # spike detection
            ap_all, ap_prop = scipy.signal.find_peaks(V_m[stim, stim_start:], min_spike_height, prominence=prominence, distance=1 * 1 / dt)
            analysis['spike_times'][stim] = ap_all * dt
            if analysis['spike_times'][stim].size == 0:
                analysis['AP'][stim] = False
            else:
                analysis['AP'][stim] = True
            analysis['amplitude'][stim] = np.array(ap_prop["peak_heights"])
            analysis['ISI'][stim] = np.array([x - analysis['spike_times'][stim][i - 1] if i else None for i, x in
                                              enumerate(analysis['spike_times'][stim])][1:])  # msec
            analysis['Freq'][stim] = 1 / (analysis['ISI'][stim] / 1000)

            if analysis['Freq'][stim].size != 0:
                analysis['F_null'][stim] = analysis['Freq'][stim][0]
                last_second_spikes = analysis['spike_times'][stim] > np.int(
                    f['data'].attrs['stim_time'] * dt - (500 + initial_period))
                if np.any(last_second_spikes == True):
                    last_second_spikes_1 = np.delete(last_second_spikes, 0)  # remove first spike because it has no ISI/freq
                    analysis['F_inf'][stim] = np.mean(analysis['Freq'][stim][last_second_spikes_1])
            else: # if no ISI detected
                analysis['F_null'][stim] = 0
                analysis['F_inf'][stim] = 0

        # rheobase
        # find where the first AP occurs
        if np.any(analysis['AP'][:] == True):
            # setup current range to be between the current step before and with first AP
            I_first_spike = I_mag[np.argwhere(analysis['AP'][:] == True)[0][0]]
            I_before_spike = I_mag[np.argwhere(analysis['AP'][:] == True)[0][0] - 1]
            analysis.create_dataset('I_first_spike', data=I_first_spike, dtype='float64')
            analysis.create_dataset('I_before_spike', data=I_before_spike, dtype='float64')
            number_steps_rheo = 100
            stim_time, I_in_rheo, stim_num_rheo, V_m_rheo = stimulus_init(I_before_spike, I_first_spike, number_steps_rheo,
                                                                               initial_period, dt, sec)

            # simulate response to this current range and save
            V_m_rheo = STN(V_init * np.ones((stim_num_rheo)), g_effect, E2, I_in_rheo, dt, currents_included2,
                           stim_time, stim_num_rheo, f['data'].attrs['C'], shift2, scale2, b_param_effect, slope_shift2,
                           np.zeros((len(b_param), stim_num_rheo)), np.zeros((len(currents_included), stim_num_rheo)),
                           ind_dict, f['data'].attrs['vol'])
            f['data'].create_dataset('V_m_rheo', data=V_m_rheo, dtype='float64', compression="gzip", compression_opts=9)

            # run AP detection
            analysis.create_dataset('spike_times_rheo', (I_in_rheo.shape[1],), dtype=dtyp, compression="gzip",
                                    compression_opts=9)
            analysis.create_dataset('AP_rheo', (I_in_rheo.shape[1],), dtype='bool', compression="gzip", compression_opts=9)

            for stim in range(I_in_rheo.shape[1]):  # range(stim_num):
                ap_all, ap_prop = scipy.signal.find_peaks(V_m_rheo[stim, stim_start:], min_spike_height,
                                                          prominence=prominence, distance=1 * 1 / dt)
                analysis['spike_times_rheo'][stim] = ap_all * dt
                if analysis['spike_times_rheo'][stim].size == 0:
                    analysis['AP_rheo'][stim] = False
                else:
                    analysis['AP_rheo'][stim] = True
            # find rheobase as smallest current step to elicit AP and save
            analysis['rheobase'][:] = I_in_rheo[-1, np.argwhere(analysis['AP_rheo'][:] == True)[0][0]] * 1000
    # AUC
    with h5py.File(fname, "a") as f:
        F_inf = f['analysis']['F_inf'][:]
        try:
            # find rheobase of steady state firing (F_inf)
            # setup current range to be in current step between no and start of steady state firing
            F_inf_start_ind = np.argwhere(F_inf != 0)[0][0]
            dI = (high - low) / stim_num
            I_start = I_mag[F_inf_start_ind - 1]
            I_end = I_mag[F_inf_start_ind]
            I_rel = np.arange(I_start, I_end + (I_end - I_start) / 100, (I_end - I_start) / 100)
            I_rel = np.reshape(I_rel, (1, I_rel.shape[0]))
            I_in = np.zeros((stim_time + np.int(initial_period * 1 / dt), 1)) @ I_rel
            I_in[np.int(initial_period * 1 / dt):, :] = np.ones((stim_time, 1)) @ I_rel
            stim_num = I_in.shape[1]

            # simulate response to this current step range
            V_m_inf_rheo = STN(V_init * np.ones(stim_num), g_effect, E2, I_in, dt, currents_included2, stim_time,
                               stim_num, C, shift2, scale2, b_param_effect, slope_shift2,
                               np.zeros((len(b_param), stim_num)), np.zeros((len(currents_included), stim_num)),
                               ind_dict, vol)
            # save response, analyse and save
            for i in np.array(['spike_times_Finf', 'ISI_Finf', 'Freq_Finf']):  # analysis_names:
                f['analysis'].require_dataset(i, shape=(I_rel.shape[1],), dtype=dtyp, compression="gzip",
                                              compression_opts=9)
            f['analysis'].require_dataset('F_inf_Finf', shape=(I_rel.shape[1],), dtype='float64')
            f['analysis'].require_dataset('F_null_Finf', shape=(I_rel.shape[1],), dtype='float64')
            for stim in range(I_rel.shape[1]):
                # spike peak detection
                ap_all, ap_prop = scipy.signal.find_peaks(V_m_inf_rheo[stim, stim_start:], min_spike_height,
                                                          prominence=prominence, distance=1 * 1 / dt)
                f['analysis']['spike_times_Finf'][stim] = ap_all * dt
                f['analysis']['ISI_Finf'][stim] = np.array(
                    [x - f['analysis']['spike_times_Finf'][stim][i - 1] if i else None for i, x in
                     enumerate(f['analysis']['spike_times_Finf'][stim])][1:])  # msec
                f['analysis']['Freq_Finf'][stim] = 1 / (f['analysis']['ISI_Finf'][stim] / 1000)
                if f['analysis']['Freq_Finf'][stim].size != 0:
                    last_second_spikes = f['analysis']['spike_times_Finf'][stim] > np.int(
                        f['data'].attrs['stim_time'] * dt - (500 + f['data'].attrs['initial_period']))
                    if np.any(last_second_spikes == True):
                        last_second_spikes_1 = np.delete(last_second_spikes,
                                                         0)  # remove first spike because it has no ISI/freq
                        f['analysis']['F_inf_Finf'][stim] = np.mean(
                            f['analysis']['Freq_Finf'][stim][last_second_spikes_1])
                    else:
                        f['analysis']['F_null_Finf'][stim] = 0
                        f['analysis']['F_inf_Finf'][stim] = 0

            # find AUC relative to F_inf rheobase
            # setup current inputs with current range around F_inf rheobase
            I_first_inf = I_rel[0, np.argwhere(f['analysis']['F_inf_Finf'][:] != 0)[0][0]]
            I_start = I_first_inf - 0.00005
            I_end = I_first_inf + desired_AUC_width * high
            I_rel2 = np.arange(I_start, I_end + (I_end - I_start) / 200, (I_end - I_start) / 200)
            I_rel2 = np.reshape(I_rel2, (1, I_rel2.shape[0]))
            I_in2 = np.zeros((stim_time + np.int(initial_period * 1 / dt), 1)) @ I_rel2
            I_in2[np.int(initial_period * 1 / dt):, :] = np.ones((stim_time, 1)) @ I_rel2
            stim_num = I_in2.shape[1]

            # simulate response to this current step range
            V_m_AUC = STN(V_init * np.ones(stim_num), g_effect, E2, I_in2, dt, currents_included2, stim_time, stim_num,
                          C, shift2, scale2, b_param_effect, slope_shift2, np.zeros((len(b_param), stim_num)),
                          np.zeros((len(currents_included), stim_num)), ind_dict, vol)

            # save data and analysis for this current step range including AUC
            f['data'].require_dataset('V_m_AUC', shape=V_m_AUC.shape, dtype='float64')
            f['data']['V_m_AUC'][:] = V_m_AUC
            for i in np.array(['spike_times_Finf_AUC', 'ISI_Finf_AUC', 'Freq_Finf_AUC']):  # analysis_names:
                f['analysis'].require_dataset(i, shape=(I_rel2.shape[1],), dtype=dtyp, compression="gzip",
                                              compression_opts=9)
            f['analysis'].require_dataset('F_inf_Finf_AUC', shape=(I_rel2.shape[1],), dtype='float64')
            f['analysis'].require_dataset('F_null_Finf_AUC', shape=(I_rel2.shape[1],), dtype='float64')
            for stim in range(I_rel2.shape[1]):
                # spike peak detection
                ap_all, ap_prop = scipy.signal.find_peaks(V_m_AUC[stim, stim_start:], min_spike_height,
                                                          prominence=prominence, distance=1 * 1 / dt)
                f['analysis']['spike_times_Finf_AUC'][stim] = ap_all * dt
                f['analysis']['ISI_Finf_AUC'][stim] = np.array(
                    [x - f['analysis']['spike_times_Finf_AUC'][stim][i - 1] if i else None for i, x in
                     enumerate(f['analysis']['spike_times_Finf_AUC'][stim])][1:])  # msec
                f['analysis']['Freq_Finf_AUC'][stim] = 1 / (f['analysis']['ISI_Finf_AUC'][stim] / 1000)
                if f['analysis']['Freq_Finf_AUC'][stim].size != 0:
                    last_second_spikes = f['analysis']['spike_times_Finf_AUC'][stim] > np.int(
                        f['data'].attrs['stim_time'] * dt - (500 + f['data'].attrs['initial_period']))
                    if np.any(last_second_spikes == True):
                        last_second_spikes_1 = np.delete(last_second_spikes,
                                                         0)  # remove first spike because it has no ISI/freq
                        f['analysis']['F_inf_Finf_AUC'][stim] = np.mean(
                            f['analysis']['Freq_Finf_AUC'][stim][last_second_spikes_1])
                    else:
                        f['analysis']['F_null_Finf_AUC'][stim] = 0
                        f['analysis']['F_inf_Finf_AUC'][stim] = 0
            f['analysis'].require_dataset('AUC', shape=(1,), dtype='float64', compression="gzip",
                                          compression_opts=9)
            f['analysis']['AUC'][:] = np.trapz(f['analysis']['F_inf_Finf_AUC'][:], I_rel2[:] * 1000, dI * 1000)
        except:
            f['analysis'].require_dataset('AUC', shape=(1,), dtype='float64', compression="gzip",
                                          compression_opts=9)
            f['analysis']['AUC'][:] = 0


    # ramp protocol #####################
    # setup ramp current
    sec = 4
    ramp_len = np.int(sec *1000 *1/dt)
    stim_time = ramp_len *2
    I_amp = np.array([0, -0.000001])
    I_amp = np.reshape(I_amp, (1,I_amp.shape[0]))
    I_step = np.zeros((stim_time + np.int(initial_period * 1 / dt), 1))@ I_amp
    I_step[np.int(initial_period * 1 / dt):, :] = np.ones((stim_time, 1)) @ I_amp
    stim_num_step = I_step.shape[1]
    start = np.int(initial_period * 1 / dt)
    I_step[start:np.int(start+ramp_len),0] = np.linspace(0,high, ramp_len)
    I_step[np.int(start + ramp_len):np.int(start + ramp_len*2),0] = np.linspace(high, 0, ramp_len)
    stim_time_step = I_step.shape[0]

    # simulate response to ramp
    V_m_step = STN(-70 * np.ones(stim_num_step), g_effect, E2, I_step, dt, currents_included2, stim_time_step,
                   stim_num_step, C, shift2, scale2, b_param_effect, slope_shift2,
                   np.zeros((len(b_param), stim_num_step)), np.zeros((len(currents_included), stim_num_step)), ind_dict,
                   vol)

    # spike peak detection
    ap_all, ap_prop = scipy.signal.find_peaks(V_m_step[0, stim_start:], min_spike_height,
                                                          prominence=prominence, distance=1 * 1 / dt)
    # save ramp firing properties
    with h5py.File(fname, "a") as f:
        f['data'].create_dataset('V_m_ramp', data=V_m_step, dtype='float64', compression="gzip", compression_opts=9)

        if ap_all.shape[0] != 0:
            if ap_all[0] < stim_start + ramp_len:
                f['analysis'].create_dataset('ramp_I_up', data =I_step[stim_start+ap_all[0], 0] *1000,  dtype='float64')
            else:
                f['analysis'].create_dataset('ramp_I_up', data =np.NaN,  dtype='float64')
            if ap_all[-1] > stim_start + ramp_len:
                f['analysis'].create_dataset('ramp_I_down', data =I_step[stim_start+ap_all[-1], 0]*1000,  dtype='float64')
            else:
                f['analysis'].create_dataset('ramp_I_down', data =np.NaN,  dtype='float64')
            f['analysis'].create_dataset('hysteresis', data =f['analysis']['ramp_I_down'][()] - f['analysis']['ramp_I_up'][()],  dtype='float64')
        else: # if no spikes in response to ramp
            f['analysis'].create_dataset('ramp_I_up', data =np.NaN,  dtype='float64')
            f['analysis'].create_dataset('ramp_I_down', data =np.NaN,  dtype='float64')
            f['analysis'].create_dataset('hysteresis', data =np.NaN,  dtype='float64')
    if f.__bool__():
        f.close()
    gc.collect()


def SA_STN(V_init, g, E, I_in, dt, currents_included, stim_time, stim_num, C, shift, scale, b_param, slope_shift,
           gating, current, prominence, desired_AUC_width, folder, high, low, number_steps, initial_period, sec, vol,
           lin_array, log_array, alt_types, alt_ind, alt):
    """ Sensitivity analysis for STN model

        Parameters
        ----------
        V_init : float
            Initial membrane potential
        g : dict
            maximal conductance for currents
        E : dict
            Reversal potentials for currents
        I_in : array
            Current stimulus input to model
        dt : float
            fixed time step
        currents_included : dict
            Boolean to include current in simulation or not
        stim_time :
            length of stimulus ()
        stim_num : int
            number of different stimulus currents
        C : float
            membrane capacitance
        shift : dict
            for each gating parameter
        scale : dict
            for each gating parameter
        b_param : dict
            Boltzmann parameter values for each gating parameter
        slope_shift : dict
            for each gating parameter
        gating : array
            array of gating values over time
        current : array
            array of current values over time
        prominence: float
            required spike prominence
        desired_AUC_width: float
            width of AUC from rheobase to rheobase + desired_AUC_width
        folder : str
            folder to save hdf5 in
        high : float
            upper current step magnitude
        low : float
            lowest current step magnitude
        number_steps : int
            number of current steps between low and high
        initial_period: float
            length  of  I = 0 before current step
        sec : float
            length of current step in seconds
        vol : float
            cell volume
        lin_array : array
            array of linear shifts from -10 tp 10
        log_array : array
            array of log2 values from -1 to 1
        alt_types : array
            array of arrays of strings of [variable, alteration type]
        alt_ind : int
            index for alt_type
        alt :
            index for lin_array of log_array

        Returns
        -------
        saves hdf5 file to folder
        """
    # setup for simulation
    g_multi = copy.deepcopy(g)
    E_multi = copy.deepcopy(E)
    b_param_multi = copy.deepcopy(b_param)
    shift_multi = copy.deepcopy(shift)
    scale_multi = copy.deepcopy(scale)
    slope_shift_multi = copy.deepcopy(slope_shift)
    currents_included_multi = copy.deepcopy(currents_included)
    gating2 = copy.deepcopy(gating)
    current2 = copy.deepcopy(current)

    g2 = Dict.empty(key_type=types.unicode_type, value_type=types.float64, )
    for k, v in g_multi.items():
        g2[k] = v
    b_param2 = Dict.empty(key_type=types.unicode_type, value_type=types.float64[:], )
    for k, v in b_param_multi.items():
        b_param2[k] = np.array(v)
    shift2 = Dict.empty(key_type=types.unicode_type, value_type=types.float64, )
    for k, v in shift_multi.items():
        shift2[k] = v
    scale2 = Dict.empty(key_type=types.unicode_type, value_type=types.float64, )
    for k, v in scale_multi.items():
        scale2[k] = v
    slope_shift2 = Dict.empty(key_type=types.unicode_type, value_type=types.float64, )
    for k, v in slope_shift_multi.items():
        slope_shift2[k] = v
    E2 = Dict.empty(key_type=types.unicode_type, value_type=types.float64, )
    for k, v in E_multi.items():
        E2[k] = v
    currents_included2 = Dict.empty(key_type=types.unicode_type, value_type=types.boolean, )
    for k, v in currents_included_multi.items():
        currents_included2[k] = v
    ind_dict = Dict.empty(key_type=types.unicode_type, value_type=types.int64, )
    i = 0
    for var in np.array(
            ['m', 'h', 'n', 'a', 'b', 'c', 'a_mut', 'b_mut', 'd1', 'd2', 'p', 'q', 'r', 'Ca_conc', 'E_Ca']):
        ind_dict[var] = i
        i += 1
    i = 0
    for var in np.array(['Na', 'Kd', 'A', 'A_mut', 'L', 'T', 'Leak', 'Ca_K']):
        ind_dict[var] = i
        i += 1

    #### alteration #######################################################################
    if alt_types[alt_ind, 1] == 'shift':
        shift2[alt_types[alt_ind, 0]] = lin_array[alt]
        fname = os.path.join(folder,
                             "{}_{}_{}.hdf5".format(alt_types[alt_ind, 1], alt_types[alt_ind, 0], lin_array[alt]))

    elif alt_types[alt_ind, 1] == 'slope':
        b_param2[alt_types[alt_ind, 0]][1] = b_param2[alt_types[alt_ind, 0]][1] * log_array[alt]
        fname = os.path.join(folder,
                             "{}_{}_{}.hdf5".format(alt_types[alt_ind, 1], alt_types[alt_ind, 0],
                                                    np.int(np.around(log_array[alt], 2) * 100)))
        if b_param2[alt_types[alt_ind, 0]][2] != 1:
            # adjust slope_shift to account for slope not changing (rotating) around 0.5
            V_test = np.arange(-150, 100, 0.001)
            if b_param2[alt_types[alt_ind, 0]].shape[0] == 4:
                steady_state = ((1 - b_param2[alt_types[alt_ind, 0]][3]) / (
                        1 + np.exp(
                    (V_test - b_param2[alt_types[alt_ind, 0]][0]) / b_param2[alt_types[alt_ind, 0]][1])) +
                                b_param2[alt_types[alt_ind, 0]][3]) ** b_param2[alt_types[alt_ind, 0]][2]
                orig = ((1 - b_param[alt_types[alt_ind, 0]][3]) / (
                        1 + np.exp((V_test - b_param[alt_types[alt_ind, 0]][0]) / b_param[alt_types[alt_ind, 0]][1])) +
                        b_param[alt_types[alt_ind, 0]][3]) ** b_param[alt_types[alt_ind, 0]][2]
            else:
                steady_state = (1 / (1 + np.exp(
                    (V_test - b_param2[alt_types[alt_ind, 0]][0]) / b_param2[alt_types[alt_ind, 0]][1]))) ** \
                               b_param2[alt_types[alt_ind, 0]][2]
                orig = (1 / (1 + np.exp(
                    (V_test - b_param[alt_types[alt_ind, 0]][0]) / b_param[alt_types[alt_ind, 0]][1]))) ** \
                       b_param[alt_types[alt_ind, 0]][2]
            orig_V_half = V_test[(np.abs(orig - 0.5)).argmin()]
            V_half_new = V_test[(np.abs(steady_state - 0.5)).argmin()]
            slope_shift2[alt_types[alt_ind, 0]] = orig_V_half - V_half_new

    elif alt_types[alt_ind, 1] == 'g':
        g2[alt_types[alt_ind, 0]] = g2[alt_types[alt_ind, 0]] * log_array[alt]
        fname = os.path.join(folder, "{}_{}_{}.hdf5".format(alt_types[alt_ind, 1], alt_types[alt_ind, 0],
                                                            np.int(np.around(log_array[alt], 2) * 100)))
    ###########################################################################

    # initial simulation for current range from low to high
    V_m = STN(V_init * np.ones((stim_num)), g2, E2, I_in, dt, currents_included2, stim_time, stim_num, C, shift2,
              scale2, b_param2, slope_shift2, gating2, current2, ind_dict, vol)
    stim_start = np.int(initial_period * 1 / dt)
    min_spike_height = V_m[0, stim_start] + prominence

    # create hdf5 file and save data and metadata
    with h5py.File(fname, "a") as f:
        data = f.create_group("data")
        data.create_dataset('V_m', data=V_m, dtype='float64', compression="gzip", compression_opts=9)
        metadata = {'I_high': high, 'I_low': low, 'stim_num': number_steps, 'stim_time': stim_time, 'sec': sec, 'dt': dt,
                    'initial_period': initial_period, 'g': json.dumps(dict(g2)), 'E': json.dumps(dict(E2)), 'C': C,
                    'V_init': V_init, 'vol': vol, 'ind_dict': json.dumps(dict(ind_dict)),
                    'b_param': json.dumps(dict(b_param2), cls=NumpyEncoder),
                    'currents': json.dumps(dict(currents_included2)), 'shift': json.dumps(dict(shift2)),
                    'var_change': str(alt_types[alt_ind, 1]), 'var': str(alt_types[alt_ind, 0]),
                    'scale': json.dumps(dict(scale2)), 'slope_shift': json.dumps(dict(slope_shift2)),
                    'prominence': prominence, 'desired_AUC_width': desired_AUC_width,
                    'alteration_info': str(alt_types[alt_ind, :]), 'alt_index': str(alt),
                    'min_spike_height': min_spike_height,}
        if alt_types[alt_ind, 1] == 'shift':
            meta2 = {'alteration': lin_array[alt]}
        else:
            meta2 = {'alteration': log_array[alt]}
        data.attrs.update(meta2)

    # firing frequency analysis and rheobase
    with h5py.File(fname, "r+") as f:
        I_mag = np.arange(low, high, (high - low) / stim_num)
        analysis = f.create_group("analysis")
        dtyp = h5py.special_dtype(vlen=np.dtype('float64'))
        for i in np.array(['spike_times', 'spike_times_rheo', 'ISI', 'Freq']):
            analysis.create_dataset(i, (I_mag.shape[0],), dtype=dtyp, compression="gzip", compression_opts=9)
        analysis.create_dataset('F_inf', (I_mag.shape[0],), dtype='float64')
        analysis.create_dataset('F_null', (I_mag.shape[0],), dtype='float64')
        analysis.create_dataset('AP', (I_mag.shape[0],), dtype='bool', compression="gzip", compression_opts=9)
        analysis.create_dataset('AP_rheo', (I_mag.shape[0],), dtype='bool', compression="gzip", compression_opts=9)
        for i in np.array(['rheobase']):
            analysis.create_dataset(i, (1,), dtype='float64', compression="gzip", compression_opts=9)
        try:

            for stim in range(I_mag.shape[0]):
                # spike detection
                ap_all, ap_prop = scipy.signal.find_peaks(V_m[stim, stim_start:], min_spike_height,
                                                          prominence=prominence, distance=1 * 1 / dt)
                analysis['spike_times'][stim] = ap_all * dt
                if analysis['spike_times'][stim].size == 0:
                    analysis['AP'][stim] = False
                else:
                    analysis['AP'][stim] = True
                analysis['ISI'][stim] = np.array([x - analysis['spike_times'][stim][i - 1] if i else None for i, x in
                                                  enumerate(analysis['spike_times'][stim])][1:])  # msec
                analysis['Freq'][stim] = 1 / (analysis['ISI'][stim] / 1000)

                if analysis['Freq'][stim].size != 0:
                    analysis['F_null'][stim] = analysis['Freq'][stim][0]
                    if np.argwhere(ap_all >= 1000 * 1 / dt).shape[0] != 0:  # if first spike is after 1 ms
                        half_second_spikes = np.logical_and(analysis['spike_times'][stim] < np.int(
                            ap_all[np.argwhere(ap_all >= 1000 * 1 / dt)[0][0]] * dt + 500),
                                                            analysis['spike_times'][stim] > np.int(
                                                                ap_all[np.argwhere(ap_all >= 1000 * 1 / dt)[0][0]] * dt))  # spikes in half second window
                        if np.sum(half_second_spikes == True) > 1:
                            half_second_spikes_1 = np.delete(half_second_spikes,
                                                             np.argwhere(half_second_spikes == True)[0][0])
                            analysis['F_inf'][stim] = np.mean(analysis['Freq'][stim][half_second_spikes_1])
                        else:
                            analysis['F_inf'][stim] = 0
                    else:
                        analysis['F_inf'][stim] = 0
                else: # if no ISI detected
                    analysis['F_null'][stim] = 0
                    analysis['F_inf'][stim] = 0

        except:
            print(alt_types[alt_ind, 0], alt_types[alt_ind, 1], alt, 'Freq analysis unsuccessful')

        # rheobase
        # find where the first AP occurs
        if np.any(analysis['AP'][:] == True):
            # setup current range to be between the current step before and with first AP
            I_first_spike = I_mag[np.argwhere(analysis['AP'][:] == True)[0][0]]
            I_before_spike = I_mag[np.argwhere(analysis['AP'][:] == True)[0][0] - 1]
            analysis.create_dataset('I_first_spike', data=I_first_spike, dtype='float64')
            analysis.create_dataset('I_before_spike', data=I_before_spike, dtype='float64')
            number_steps_rheo = 100
            stim_time, I_in_rheo, stim_num_rheo, V_m_rheo = stimulus_init(I_before_spike, I_first_spike,
                                                                               number_steps_rheo,
                                                                               initial_period, dt, sec)

            # simulate response to this current range and save
            V_m_rheo = STN(V_init * np.ones((stim_num_rheo)), g2, E2, I_in_rheo, dt, currents_included2, stim_time,
                           stim_num_rheo, C, shift2, scale2, b_param2, slope_shift2,
                           np.zeros((len(b_param), stim_num_rheo)), np.zeros((len(currents_included), stim_num_rheo)),
                           ind_dict, vol)
            try:
                # run AP detection
                for stim in range(I_in_rheo.shape[1]):  # range(stim_num):
                    ap_all, ap_prop = scipy.signal.find_peaks(V_m_rheo[stim, stim_start:], min_spike_height,
                                                              prominence=prominence, distance=1 * 1 / dt)
                    analysis['spike_times_rheo'][stim] = ap_all * dt
                    if analysis['spike_times_rheo'][stim].size == 0:
                        analysis['AP_rheo'][stim] = False
                    else:
                        analysis['AP_rheo'][stim] = True
                # find rheobase as smallest current step to elicit AP and save
                analysis['rheobase'][:] = I_in_rheo[-1, np.argwhere(analysis['AP_rheo'][:] == True)[0][0]] * 1000
            except:
                print(alt_types[alt_ind, 0], alt_types[alt_ind, 1], alt, 'RHEOBASE NO AP')


    # AUC
    with h5py.File(fname, "a") as f:
        # find rheobase of steady state firing (F_inf)
        F_inf = f['analysis']['F_inf'][:]
        try:
            # setup current range to be in current step between no and start of steady state firing
            F_inf_start_ind = np.argwhere(F_inf != 0)[0][0]
            dI = (high- low) / stim_num
            I_start = I_mag[F_inf_start_ind - 1]
            I_end = I_mag[F_inf_start_ind]
            I_rel = np.arange(I_start, I_end + (I_end - I_start) / 100, (I_end - I_start) / 100)
            I_rel = np.reshape(I_rel, (1, I_rel.shape[0]))
            I_in = np.zeros((stim_time + np.int(initial_period * 1 / dt), 1)) @ I_rel
            I_in[np.int(initial_period * 1 / dt):, :] = np.ones((stim_time, 1)) @ I_rel
            stim_num = I_in.shape[1]

            # simulate response to this current step range
            V_m_inf_rheo = STN(V_init * np.ones(stim_num), g2, E2, I_in, dt, currents_included2, stim_time, stim_num, C,
                               shift2, scale2, b_param2, slope_shift2, np.zeros((len(b_param), stim_num)),
                               np.zeros((len(currents_included), stim_num)), ind_dict, vol)

            # save response, analyse and save
            for i in np.array(['spike_times_Finf', 'ISI_Finf', 'Freq_Finf']):
                f['analysis'].require_dataset(i, shape=(I_rel.shape[1],), dtype=dtyp, compression="gzip",
                                              compression_opts=9)
            f['analysis'].require_dataset('F_inf_Finf', shape=(I_rel.shape[1],), dtype='float64')
            f['analysis'].require_dataset('F_null_Finf', shape=(I_rel.shape[1],), dtype='float64')
            for stim in range(I_rel.shape[1]):
                # spike peak detection
                ap_all, ap_prop = scipy.signal.find_peaks(V_m_inf_rheo[stim, stim_start:], min_spike_height,
                                                          prominence=prominence, distance=1 * 1 / dt)
                f['analysis']['spike_times_Finf'][stim] = ap_all * dt
                f['analysis']['ISI_Finf'][stim] = np.array(
                    [x - f['analysis']['spike_times_Finf'][stim][i - 1] if i else None for i, x in
                     enumerate(f['analysis']['spike_times_Finf'][stim])][1:])  # msec
                f['analysis']['Freq_Finf'][stim] = 1 / (f['analysis']['ISI_Finf'][stim] / 1000)
                if f['analysis']['Freq_Finf'][stim].size != 0:
                    if np.argwhere(ap_all >= 1000 * 1 / dt).shape[0] != 0:  # if first spike is after 1 ms
                        half_second_spikes = np.logical_and(f['analysis']['spike_times_Finf'][stim] < np.int(
                            ap_all[np.argwhere(ap_all >= 1000 * 1 / dt)[0][0]] * dt + 500),
                                                            f['analysis']['spike_times_Finf'][stim] > np.int(
                                                                ap_all[np.argwhere(ap_all >= 1000 * 1 / dt)[0][0]] * dt))  # spikes in half second window
                        if np.sum(half_second_spikes == True) > 1:
                            half_second_spikes_1 = np.delete(half_second_spikes,
                                                             np.argwhere(half_second_spikes == True)[0][0])  # remove first spike because it has no ISI/freq
                            f['analysis']['F_inf_Finf'][stim] = np.mean(f['analysis']['Freq_Finf'][stim][half_second_spikes_1])
                        else:
                            f['analysis']['F_inf_Finf'][stim] = 0
                    else:
                        f['analysis']['F_null_Finf'][stim] = 0
                        f['analysis']['F_inf_Finf'][stim] = 0
                else:
                    f['analysis']['F_null_Finf'][stim] = 0
                    f['analysis']['F_inf_Finf'][stim] = 0

            # find AUC relative to F_inf rheobase
            # setup current inputs with current range around F_inf rheobase
            I_first_inf = I_rel[0, np.argwhere(f['analysis']['F_inf_Finf'][:] != 0)[0][0]]
            I_start = I_first_inf - 0.00005
            I_end = I_first_inf + 0.2 * high
            I_rel2 = np.arange(I_start, I_end + (I_end - I_start) / 100, (I_end - I_start) / 100)
            I_rel2 = np.reshape(I_rel2, (1, I_rel2.shape[0]))
            I_in2 = np.zeros((stim_time + np.int(initial_period * 1 / dt), 1)) @ I_rel2
            I_in2[np.int(initial_period * 1 / dt):, :] = np.ones((stim_time, 1)) @ I_rel2
            stim_num = I_in2.shape[1]

            # simulate response to this current step range
            V_m_AUC = STN(V_init * np.ones(stim_num), g2, E2, I_in2, dt, currents_included2, stim_time, stim_num, C,
                          shift2, scale2, b_param2, slope_shift2, np.zeros((len(b_param), stim_num)),
                          np.zeros((len(currents_included), stim_num)), ind_dict, vol)
            # save data and analysis for this current step range including AUC
            f['data'].require_dataset('V_m_AUC', shape=V_m_AUC.shape, dtype='float64')
            f['data']['V_m_AUC'][:] = V_m_AUC
            dtyp = h5py.special_dtype(vlen=np.dtype('float64'))
            for i in np.array(['spike_times_Finf_AUC', 'ISI_Finf_AUC', 'Freq_Finf_AUC']):  # analysis_names:
                f['analysis'].require_dataset(i, shape=(I_rel2.shape[1],), dtype=dtyp, compression="gzip",
                                              compression_opts=9)
            f['analysis'].require_dataset('F_inf_Finf_AUC', shape=(I_rel2.shape[1],), dtype='float64')
            f['analysis'].require_dataset('F_null_Finf_AUC', shape=(I_rel2.shape[1],), dtype='float64')
            for stim in range(I_rel2.shape[1]):
                # spike peak detection
                ap_all, ap_prop = scipy.signal.find_peaks(V_m_AUC[stim, stim_start:], min_spike_height,
                                                          prominence=prominence, distance=1 * 1 / dt)
                f['analysis']['spike_times_Finf_AUC'][stim] = ap_all * dt
                f['analysis']['ISI_Finf_AUC'][stim] = np.array(
                    [x - f['analysis']['spike_times_Finf_AUC'][stim][i - 1] if i else None for i, x in
                     enumerate(f['analysis']['spike_times_Finf_AUC'][stim])][1:])  # msec
                f['analysis']['Freq_Finf_AUC'][stim] = 1 / (f['analysis']['ISI_Finf_AUC'][stim] / 1000)
                if f['analysis']['Freq_Finf_AUC'][stim].size != 0:
                    if np.argwhere(ap_all >= 1000 * 1 / dt).shape[0] != 0:  # if first spike is after 1 ms
                        half_second_spikes = np.logical_and(f['analysis']['spike_times_Finf_AUC'][stim] < np.int(
                            ap_all[np.argwhere(ap_all >= 1000 * 1 / dt)[0][0]] * dt + 500),
                                                            f['analysis']['spike_times_Finf_AUC'][stim] > np.int(
                                                                ap_all[np.argwhere(ap_all >= 1000 * 1 / dt)[0][0]] * dt))  # spikes in half second window
                        if np.sum(half_second_spikes == True) > 1:
                            half_second_spikes_1 = np.delete(half_second_spikes,
                                                             np.argwhere(half_second_spikes == True)[0][
                                                                 0])  # remove first spike because it has no ISI/freq
                            f['analysis']['F_inf_Finf_AUC'][stim] = np.mean(
                                f['analysis']['Freq_Finf_AUC'][stim][half_second_spikes_1])
                        else:
                            f['analysis']['F_inf_Finf_AUC'][stim] = 0
                    else:
                        f['analysis']['F_null_Finf_AUC'][stim] = 0
                        f['analysis']['F_inf_Finf_AUC'][stim] = 0
                else:
                    f['analysis']['F_null_Finf_AUC'][stim] = 0
                    f['analysis']['F_inf_Finf_AUC'][stim] = 0
            f['analysis'].require_dataset('AUC', shape=(1,), dtype='float64', compression="gzip",
                                          compression_opts=9)
            f['analysis']['AUC'][:] = np.trapz(f['analysis']['F_inf_Finf_AUC'][:],
                                                       I_rel2[:] * 1000, dI * 1000)
        except:
            f['analysis'].require_dataset('AUC', shape=(1,), dtype='float64', compression="gzip",
                                          compression_opts=9)
            f['analysis']['AUC'][:] = 0
    if f.__bool__():
        f.close()
    gc.collect()
```
