## ExtendedData1 for "Loss or Gain of Function? Effects of Ion Channel Mutations on Neuronal Firing Depend on the Cell Type": Utility_fxns.py

```
"""
General functions used throughout simulation scripts

"""
__author__ = "Nils A. Koch"
__copyright__ = 'Copyright 2022, Nils A. Koch'
__license__ = "MIT"


import json
import numpy as np
from numba import njit

@njit
def sqrt_log(x, *p):
    """ Function of sum of square root and logarithmic function to fit to fI curves
    Parameters
    ----------
    x : array
        input x value
    *p : array
        parameters for function

    Returns
    -------
    p[0] * np.sqrt(x + p[1]) + p[2] * np.log(x + (p[1] + 1)) + p[3]
    """
    return p[0] * np.sqrt(x + p[1]) + p[2] * np.log(x + (p[1] + 1)) + p[3]


@njit
def capacitance(diam, C_m):
    """ calculate the capacitance of a cell's membrane give the diameter
    Parameters
    ----------
    diam: float
        diameter of spherical cell body
    C_m: float
        specific membrane capacitance of cell

    Returns
    -------
    capacitance: float
        capacitance of cell
    surf_area: flaot
        surface area of cell
    """
    r = diam * 1E-4 / 2  # cm
    l = r
    surf_area = 2 * np.pi * r ** 2 + 2 * np.pi * r * l  # cm^2
    C = C_m * surf_area  # uF
    return (C, surf_area)


@njit
def stimulus_init(low, high, number_steps, initial_period, dt, sec):
    """ initiation of stimulus pulse from stimulus magnitudes I_in_mag
    Parameters
    ----------
    low : float
        lowest current step magnitude
    high : float
        upper current step magnitude
    number_steps : int
        number of current steps between low and high
    initial_period : float
        length of I=0 before current step
    dt : float
        time step
    sec:
        length of current step in seconds

    Returns
    -------
    stim_time: float
        length of I_in in samples
    I_in: array
        I input array
    stim_num: int
        number of current steps
    V_m: array
        membrane potential array initialization of shape:(stim_num, stim_time)
    """
    stim_time = np.int(1000 * 1 / dt * sec)  # 1000 msec/sec * 1/dt
    I_in_mag = np.arange(low, high, (high - low) / number_steps)
    I_in_mag = np.reshape(I_in_mag, (1, I_in_mag.shape[0]))
    I_in = np.zeros((stim_time + np.int(initial_period * 1 / dt), 1)) @ I_in_mag
    I_in[np.int(initial_period * 1 / dt):, :] = np.ones((stim_time, 1)) @ I_in_mag
    stim_num = I_in.shape[1]  # number of different currents injected
    stim_time = stim_time + np.int(initial_period * 1 / dt)
    V_m = np.zeros((stim_num, stim_time))

    return stim_time, I_in, stim_num, V_m

class NumpyEncoder(json.JSONEncoder):
    """ Special json encoder for numpy types """
    def default(self, obj):
        if isinstance(obj, np.integer):
            return int(obj)
        elif isinstance(obj, np.floating):
            return float(obj)
        elif isinstance(obj, np.ndarray):
            return obj.tolist()
        return json.JSONEncoder.default(self, obj)


def init_dict(variable):
    """ Initialize dictionaries for simulations
    Parameters
    ----------
    variable : array
        array of variable name strings to put in dicts

    Returns
    -------
    shift : dict
        with every variable name in variable = 0.
    scale : dict
        with every variable name in variable = 1.
    slope_shift : dict
        with every variable name in variable = 0.
    E : dict
        empty dictionary
    currents_included : dict
        empty dictionary
    b_param : dict
        with every variable name in variable = np.array of 2 zeros
    g : dict
        empty dictionary
    """
    shift = {}
    scale = {}
    slope_shift = {}
    b_param ={}
    g = {}
    E ={}
    currents_included = {}
    for var in variable:
        shift[var] = 0.
        scale[var] = 1.0
        slope_shift[var] = 0.
        b_param[var] = np.zeros(3, dtype=np.dtype('float64'))
    return shift, scale, slope_shift, E, currents_included, b_param, g
```
