## ExtendedData1 for "Loss or Gain of Function? Effects of Ion Channel Mutations on Neuronal Firing Depend on the Cell Type": mut_Cb_stellate_Delta_Kv.py

```
# -*- coding: utf-8 -*-
"""
Script to simulate all KCNA1 mutations in Cb Stellate \Delta Kv1.1 model

"""
__author__ = "Nils A. Koch"
__copyright__ = 'Copyright 2022, Nils A. Koch'
__license__ = "MIT"


import numpy as np
from joblib import Parallel, delayed
import json
import os
from numba import types
from numba.typed import Dict
from Code.Functions.Utility_fxns import capacitance, stimulus_init, init_dict
from Code.Functions.Cb_stellate_fxns_Kv import Cb_stellate_Kv_mut


#%%
# model parameters
dt = 0.01
sec = 2
low = 0
high = 0.0015
number_steps = 200
initial_period = 1000
num_gating = 11
num_current = 7
C, surf_area = capacitance(61.4, 1.50148)
variable = np.array(['m', 'h', 'n', 'n_A', 'h_A', 'm_T', 'h_T', 's', 'u', 's_mut', 'u_mut'])
stim_time, I_in, stim_num, V_m = stimulus_init(low, high, number_steps, initial_period, dt, sec)
shift, scale, slope_shift,  E, currents_included, b_param, g = init_dict(np.array(['m', 'h', 'n', 'n_A', 'h_A', 'm_T', 'h_T', 's', 'u', 's_mut', 'u_mut']))
V_init = -70

# initialize arrays
current = np.zeros((num_current, stim_num))
gating = np.zeros((num_gating, stim_num))

# initialize dictionary
ind_dict = Dict.empty(key_type=types.unicode_type, value_type=types.int64, )
i = 0
for var in np.array(['m', 'h', 'n', 'q', 'r', 'p', 's', 'u', 's_mut', 'u_mut']):
    ind_dict[var] = i
    i += 1
i = 0
for var in np.array(['Na', 'Kd', 'M', 'Kv', 'Kv_mut', 'L', 'Leak']):
    ind_dict[var] = i
    i += 1

# gating parameters
b_param['m'][:] = np.array([-37.,  -3,   1])
b_param['h'] = np.zeros(4)
b_param['h'][:] = np.array([-40.,   4.,   1., 0])
b_param['n'][:] = np.array([-23, -5,   1])
b_param['n_A'][:] = np.array([-27, -13.2,   1.])
b_param['h_A'][:] = np.array([-80., 6.5,   1.])
b_param['m_T'][:] = np.array([-50., -3,   1.])
b_param['h_T'][:] = np.array([-68., 3.75, 1.])
b_param['s'][:] = np.array([-14.16, -10.15, 1.])
b_param['u'] = np.zeros(4)
b_param['u'][:] = np.array([-31., 5.256, 1., 0.245])
b_param['s_mut'][:] = np.array([-14.16, -10.15, 1.])
b_param['u_mut'] = np.zeros(4)
b_param['u_mut'][:] = np.array([-31., 5.256, 1., 0.245])

mut_act_Vhalf_wt = -30.01851851851851
mut_act_k_wt = -7.7333333333333325
s_diff_Vhalf = mut_act_Vhalf_wt - b_param['s'][0]
s_diff_k = mut_act_k_wt - b_param['s'][1]
b_param['s'][1] = b_param['s'][1] + s_diff_k
b_param['u'][1] = b_param['u'][1] + s_diff_k
b_param['s'][0]  = b_param['s'][0] + s_diff_Vhalf
b_param['u'][0]  = b_param['u'][0] + s_diff_Vhalf
b_param['s_mut'][1] = b_param['s_mut'][1] + s_diff_k
b_param['u_mut'][1] = b_param['u_mut'][1] + s_diff_k
b_param['s_mut'][0]  = b_param['s_mut'][0] + s_diff_Vhalf
b_param['u_mut'][0]  = b_param['u_mut'][0] + s_diff_Vhalf

# reversal potentials
E["Na"] = 55.
E["K"] = -80.
E["Ca"] = 22.
E["Leak"] = -70. # as per Molineux et al 2005 and NOT the -38 in Alexander et al 2019

# model currents
currents_included["Na"] = True
currents_included["Kd"] = True
currents_included["Kv"] = True
currents_included["Kv_mut"] = True
currents_included["A"] = False
currents_included["T"] = True
currents_included["Leak"] = True

# model conductances
g["Na"] = 34 * surf_area
g["Kd"] = 9.0556 * surf_area
if currents_included["Kv_mut"] == True:
    g["Kv"] = 1.50159  / 2 * surf_area
else:
    g["Kv"] = 1.50159 / 2 * surf_area * 2
g["Kv_mut"] = 1.50159  / 2 * surf_area
g["A"] = 0 * surf_area
g["T"] = 0.45045 * surf_area
g["Leak"] = 0.07407 * surf_area

# save folder
folder = './KCNA1_mutations/Cb_stellate_Delta_Kv'
if not os.path.isdir(folder):
    os.makedirs(folder)
#
# mutation properties
mutations = json.load(open("./KCNA1_mutations/mutations_effects_dict.json"))

# %%
prominence = 50
desired_AUC_width = high/5
Parallel(n_jobs=8, verbose=9)(
    delayed(Cb_stellate_Kv_mut)(V_init, g, E, I_in, dt, currents_included, stim_time, stim_num, C, shift, scale,
                                b_param, slope_shift, gating, current, prominence, desired_AUC_width, mutations, mut, folder, high, low,
                                number_steps, initial_period, sec) for mut in list(mutations.keys()))
```
