## ExtendedData1 for "Loss or Gain of Function? Effects of Ion Channel Mutations on Neuronal Firing Depend on the Cell Type": mut_STN.py

```
# -*- coding: utf-8 -*-
"""
Script to simulate all KCNA1 mutations in STN model

"""
__author__ = "Nils A. Koch"
__copyright__ = 'Copyright 2022, Nils A. Koch'
__license__ = "MIT"

import numpy as np
from joblib import Parallel, delayed
import json
import os
from numba import types
from numba.typed import Dict
from Code.Functions.Utility_fxns import capacitance, stimulus_init, init_dict
from Code.Functions.STN_fxns import STN_mut


# model parameters
dt = 0.01
sec = 2
low = 0
high = 0.005
number_steps = 100
initial_period = 1000
num_gating = 15
num_current = 8
d = 61.4
r = d/2 * 10**-6 # radius in meters
vol = np.pi * r**3 # volume in liters
C, surf_area = capacitance(d, 1)
stim_time, I_in, stim_num, V_m = stimulus_init(low, high, number_steps, initial_period, dt, sec)
shift, scale, slope_shift,  E, currents_included, b_param, g = init_dict(np.array(['m', 'h', 'n', 'a', 'b', 'a_mut', 'b_mut', 'c', 'd1', 'd2', 'p', 'q', 'r', 'Ca_conc', 'E_Ca']))

# initialize arrays
current = np.zeros((num_current, stim_num))
gating = np.zeros((num_gating, stim_num))

# initialize dictionary
ind_dict = Dict.empty(key_type=types.unicode_type, value_type=types.int64, )
i = 0
for var in np.array(['m', 'h', 'n', 'q', 'r', 'p', 's', 'u', 's_mut', 'u_mut']):
    ind_dict[var] = i
    i += 1
i = 0
for var in np.array(['Na', 'Kd', 'M', 'Kv', 'Kv_mut', 'L', 'Leak']):
    ind_dict[var] = i
    i += 1

# gating parameters
b_param['m'] = np.array([-40., -1. ,   1.])
b_param['h'] = np.zeros(4)
b_param['h'] = np.array([-45.5,  1. ,   1., 0.05])
b_param['n'] = np.array([-41., -1.,   1.])
b_param['p'] = np.array([-56., -1., 1.])
b_param['q'] = np.array([-85., 1., 1.])
b_param['r'] = np.array([0.17, -0.08, 1.])
b_param['a'] = np.array([-45.,   -14.7,   1.])
b_param['b'] = np.array([-90., 7.5,   1.])
b_param['a_mut'] = np.array([-45.,   -14.7,   1.])
b_param['b_mut'] = np.array([-90., 7.5,   1.])
b_param['c'] = np.array([-30.6,  -5.,   1.])
b_param['d1'] = np.array([-60, 7.5, 1.])
b_param['d2'] = np.array([0.1, 0.02, 1.])

# reversal potentials
E["Na"] = 60.
E["K"] = -90.
E["Leak"] = -60.

# model currents
currents_included["Na"] = True
currents_included["Kd"] =True
currents_included["L"] = True
currents_included["T"] = True
currents_included["Ca_K"] = True
currents_included["A"] =True
currents_included["A_mut"] =True
currents_included["Leak"] = True

# model conductances
Kv_ratio = 0.1
g["Na"] = 49. * surf_area
g["Kd"] = 27. * (1-Kv_ratio)* surf_area
g["A"] = 5. * surf_area *0.5
g["A_mut"] = 5. * surf_area *0.5
g["L"] = 5. * surf_area
g["T"] = 5. * surf_area
g["Ca_K"] = 1 * surf_area
g["Leak"] = 0.035 * surf_area
V_init = -70

# save folder
folder = './KCNA1_mutations/STN'
if not os.path.isdir(folder):
    os.makedirs(folder)
#
# mutation properties
mutations = json.load(open("./KCNA1_mutations/mutations_effects_dict.json"))

prominence = 50
desired_AUC_width = high/5
#%%
Parallel(n_jobs=8, verbose=9)(
    delayed(STN_mut)(V_init, g, E, I_in, dt, currents_included, stim_time, stim_num, C, shift, scale, b_param, slope_shift,
            gating, current, prominence, desired_AUC_width, mutations, mut, folder, high, low, number_steps,
            initial_period, sec, vol) for mut in list(mutations.keys()))
```
