## ExtendedData1 for "Loss or Gain of Function? Effects of Ion Channel Mutations on Neuronal Firing Depend on the Cell Type": Cb_stellate_Kv_model.py

```
# -*- coding: utf-8 -*-
"""
Script to run Cb Stellate +Kv1.1 model

"""
__author__ = "Nils A. Koch"
__copyright__ = "Copyright 2022, Nils A. Koch"
__license__ = "MIT"

import numpy as np
import os
from Code.Functions.Utility_fxns import capacitance, stimulus_init, init_dict
from Code.Functions.Cb_stellate_fxns_Kv import Cb_stellate_Kv_mut

# model parameters
dt = 0.01
sec = 2
low = 0
high = 0.0015
number_steps = 200
initial_period = 1000
num_gating = 11
num_current = 7
C, surf_area = capacitance(61.4, 1.50148)
variable = np.array(['m', 'h', 'n', 'n_A', 'h_A', 'm_T', 'h_T', 's', 'u', 's_mut', 'u_mut'])
stim_time, I_in, stim_num, V_m = stimulus_init(low, high, number_steps, initial_period, dt, sec)
shift, scale, slope_shift,  E, currents_included, b_param, g = init_dict(np.array(['m', 'h', 'n', 'n_A', 'h_A', 'm_T', 'h_T', 's', 'u', 's_mut', 'u_mut']))
V_init = -70

# initialize arrays
current = np.zeros((num_current, stim_num))
gating = np.zeros((num_gating, stim_num))

# gating parameters
b_param['m'][:] = np.array([-37.,  -3,   1])
b_param['h'] = np.zeros(4)
b_param['h'][:] = np.array([-40.,   4.,   1., 0])
b_param['n'][:] = np.array([-23, -5,   1])
b_param['n_A'][:] = np.array([-27, -13.2,   1.])
b_param['h_A'][:] = np.array([-80., 6.5,   1.])
b_param['m_T'][:] = np.array([-50., -3,   1.])
b_param['h_T'][:] = np.array([-68., 3.75, 1.])
b_param['s'][:] = np.array([-14.16, -10.15, 1.])
b_param['u'] = np.zeros(4)
b_param['u'][:] = np.array([-31., 5.256, 1., 0.245])
b_param['s_mut'][:] = np.array([-14.16, -10.15, 1.])
b_param['u_mut'] = np.zeros(4)
b_param['u_mut'][:] = np.array([-31., 5.256, 1., 0.245])


# reversal potentials
E["Na"] = 55.
E["K"] = -80.
E["Ca"] = 22.
E["Leak"] = -70. # as per Molineux et al 2005 and NOT the -38 in Alexander et al 2019

# model currents
currents_included["Na"] = True
currents_included["Kd"] = True
currents_included["Kv"] = True
currents_included["Kv_mut"] = True
currents_included["A"] = True
currents_included["T"] = True
currents_included["Leak"] = True

# model conductances
Kv_ratio = 0.1
g["Na"] = 3.4 * surf_area
g["Kd"] = 9.0556 * (1-Kv_ratio) * surf_area
g["Kv"] = 6. * Kv_ratio/2 * surf_area * (2 * int(currents_included["Kv_mut"] == False))
g["Kv_mut"] = 6. * Kv_ratio/2 * surf_area
g["A"] = 15.0159 * surf_area
g["T"] = 0.45045 * surf_area
g["Leak"] = 0.07407 * surf_area


prominence = 50
min_spike_height = 0
desired_AUC_width = high/5

folder = '../Neuron_models'
mut = 'Cb_stellate_Kv'
mutations = {mut:{'g_ratio': 1, 'activation_Vhalf_diff': 0, 'activation_k_ratio':0}}
if not os.path.isdir(folder):
    os.makedirs(folder)

Cb_stellate_Kv_mut(V_init, g, E, I_in, dt, currents_included, stim_time, stim_num, C, shift, scale, b_param, slope_shift,
                    gating, current, prominence, desired_AUC_width, mutations, mut, folder, high, low, number_steps,
                    initial_period, sec)
```
