## ExtendedData1 for "Loss or Gain of Function? Effects of Ion Channel Mutations on Neuronal Firing Depend on the Cell Type": FS_model.py

```
# -*- coding: utf-8 -*-
"""
Script to run FS model

"""
__author__ = "Nils A. Koch"
__copyright__ = "Copyright 2022, Nils A. Koch"
__license__ = "MIT"

import numpy as np
from numba import types
from numba.typed import Dict
import os
from Code.Functions.Utility_fxns import capacitance, stimulus_init, init_dict
from Code.Functions.Pospischil_fxns import Pospischil_mut


# model parameters
dt = 0.01
sec = 2
low = 0
high = 0.001
number_steps = 200
initial_period = 1000
num_gating = 10
num_current = 7
C, surf_area = capacitance(56.9, 1)
stim_time, I_in, stim_num, V_m = stimulus_init(low, high, number_steps, initial_period, dt, sec)
shift, scale, slope_shift, E, currents_included, b_param, g = init_dict(
    np.array(['m', 'h', 'n', 'q', 'r', 'p', 's', 'u', 's_mut', 'u_mut']))

# initialize arrays
current = np.zeros((num_current, stim_num))
gating = np.zeros((num_gating, stim_num))

# initialize dictionary
ind_dict = Dict.empty(key_type=types.unicode_type, value_type=types.int64, )
i = 0
for var in np.array(['m', 'h', 'n', 'q', 'r', 'p', 's', 'u', 's_mut', 'u_mut']):
    ind_dict[var] = i
    i += 1
i = 0
for var in np.array(['Na', 'Kd', 'M', 'Kv', 'Kv_mut', 'L', 'Leak']):
    ind_dict[var] = i
    i += 1

# gating parameters
b_param['m'][:] = np.array([-34.33054521, -8.21450277, 1.42295686])
b_param['h'] = np.zeros(4)
b_param['h'][:] = np.array([-34.51951036, 4.04059373, 1., 0.])
b_param['n'][:] = np.array([-63.76096946, -13.83488194, 7.35347425])
b_param['q'][:] = np.array([-39.03684525, -5.57756176, 2.25190197])
b_param['r'][:] = np.array([-57.37, 20.98, 1.])
b_param['p'][:] = np.array([-45., -9.9998807337, 1.])
b_param['s'][:] = np.array([-14.16, -10.15, 1.])
b_param['u'] = np.zeros(4)
b_param['u'][:] = np.array([-31., 5.256, 1., 0.245])
b_param['s_mut'][:] = np.array([-14.16, -10.15, 1.])
b_param['u_mut'] = np.zeros(4)
b_param['u_mut'][:] = np.array([-31., 5.256, 1., 0.245])

# reversal potentials
E["Na"] = 50.
E["K"] = -90.
E["Ca"] = 120.
E["Leak"] = -70.4

# currents in model
currents_included["Na"] = True
currents_included["Kd"] = True
currents_included["Kv"] = False
currents_included["Kv_mut"] = False
currents_included["L"] = False
currents_included["M"] = True
currents_included["Leak"] = True

# model conductances
g["Na"] = 58. * surf_area
g["Kd"] = 3.9  * surf_area
g["M"] = 0.075 * surf_area
g["Kv"] = 0.
g["Kv_mut"] = 0.
g["L"] = 0.
g["Leak"] = 0.038 * surf_area
tau_max_p = 502
V_init = -70
V_T = -57.9
prominence = 50
min_spike_height = 0
desired_AUC_width = high/5

folder = '../Neuron_models'
if not os.path.isdir(folder):
    os.makedirs(folder)
mut = 'FS'
mutations = {mut:{'g_ratio': 1, 'activation_Vhalf_diff': 0, 'activation_k_ratio':0}}
Pospischil_mut(V_init, V_T, g, E, I_in, dt, currents_included, stim_time, stim_num, C, tau_max_p, shift, scale,
                   b_param, slope_shift, gating, current, prominence, desired_AUC_width, mutations, mut, folder,
                   high, low, number_steps, initial_period, sec)
```
