## ExtendedData1 for "Loss or Gain of Function? Effects of Ion Channel Mutations on Neuronal Firing Depend on the Cell Type": run_simulations.sh

```
# Bash shell script to run all simulations and analysis

# Individual Neuronal models
python ./Neuron_models/RS_pyramidal_model.py
python ./Neuron_models/RS_pyramidal_Kv_model.py
python ./Neuron_models/RS_inhib_model.py
python ./Neuron_models/RS_inhib_Kv_model.py
python ./Neuron_models/FS_model.py
python ./Neuron_models/FS_Kv_model.py
python ./Neuron_models/STN_model.py
python ./Neuron_models/STN_Kv_model.py
python ./Neuron_models/STN_Delta_Kv_model.py
python ./Neuron_models/Cb_stellate_model.py
python ./Neuron_models/Cb_stellate_Kv_model.py
python ./Neuron_models/Cb_stellate_Delta_Kv_model.py


# Sensitivity Analysis
python ./Sensitivity_Analysis/SA_RS_pyramidal.py
python ./Sensitivity_Analysis/SA_RS_pyramidal_Kv.py
python ./Sensitivity_Analysis/SA_RS_inhib.py
python ./Sensitivity_Analysis/SA_RS_inhib_Kv.py
python ./Sensitivity_Analysis/SA_FS.py
python ./Sensitivity_Analysis/SA_FS_Kv.py
python ./Sensitivity_Analysis/SA_STN.py
python ./Sensitivity_Analysis/SA_STN_Kv.py
python ./Sensitivity_Analysis/SA_STN_Delta_Kv.py
python ./Sensitivity_Analysis/SA_Cb_stellate.py
python ./Sensitivity_Analysis/SA_Cb_stellate_Kv.py
python ./Sensitivity_Analysis/SA_Cb_stellate_Delta_Kv.py


# KCNA1 mutations in models
python ./KCNA1_mutatoins/mut_RS_pyramidal_Kv.py
python ./KCNA1_mutatoins/mut_RS_inhib_Kv.py
python ./KCNA1_mutatoins/mut_FS_Kv.py
python ./KCNA1_mutatoins/mut_STN.py
python ./KCNA1_mutatoins/mut_STN_Kv.py
python ./KCNA1_mutatoins/mut_STN_Delta_Kv.py
python ./KCNA1_mutatoins/mut_Cb_stellate.py
python ./KCNA1_mutatoins/mut_Cb_stellate_Kv.py
python ./KCNA1_mutatoins/mut_Cb_stellate_Delta_Kv.py


# Data collection and csv generation
python SA_collection.py
python Plotting_data_collection.py
```
