## ExtendedData1 for "Loss or Gain of Function? Effects of Ion Channel Mutations on Neuronal Firing Depend on the Cell Type": SA_Cb_stellate.py

```
# -*- coding: utf-8 -*-
"""
Script to run sensitivity analysis for Cb Stellate model

"""
__author__ = "Nils A. Koch"
__copyright__ = "Copyright 2022, Nils A. Koch"
__license__ = "MIT"

import numpy as np
from numba import types
from numba.typed import Dict
from joblib import Parallel, delayed
import os
from Code.Functions.Utility_fxns import capacitance, stimulus_init, init_dict
from Code.Functions.Cb_stellate_fxns import SA_Cb_stellate

# model parameters
dt = 0.01
sec = 2
low = 0
high = 0.001
number_steps = 200
initial_period = 1000
num_gating = 9
num_current = 6
C, surf_area = capacitance(61.4, 1.50148)
variable = np.array(['m', 'h', 'n', 'n_A', 'h_A', 'n_A_mut', 'h_A_mut',  'm_T', 'h_T'])
stim_time, I_in, stim_num, V_m = stimulus_init(low, high, number_steps, initial_period, dt, sec)
shift, scale, slope_shift,  E, currents_included, b_param, g = init_dict(np.array(['m', 'h', 'n', 'n_A', 'h_A', 'n_A_mut', 'h_A_mut',  'm_T', 'h_T']))
V_init = -70

# initialize arrays
current = np.zeros((num_current, stim_num))
gating = np.zeros((num_gating, stim_num))

# initialize dictionary
ind_dict = Dict.empty(key_type=types.unicode_type, value_type=types.int64, )
i = 0
for var in np.array(['m', 'h', 'n', 'n_A', 'h_A', 'n_A_mut', 'h_A_mut', 'm_T', 'h_T']):
    ind_dict[var] = i
    i += 1
i = 0
for var in np.array(['Na', 'Kd', 'A', 'A_mut', 'T', 'Leak']):
    ind_dict[var] = i
    i += 1

# gating parameters
b_param['m'][:] = np.array([-37.,  -3,   1])
b_param['h'] = np.zeros(4)
b_param['h'][:] = np.array([-40.,   4.,   1., 0])
b_param['n'][:] = np.array([-23, -5,   1])
b_param['n_A'][:] = np.array([-27, -13.2,   1.])
b_param['h_A'][:] = np.array([-80., 6.5,   1.])
b_param['n_A_mut'][:] = np.array([-27, -13.2,   1.])
b_param['h_A_mut'][:] = np.array([-80., 6.5,   1.])
b_param['m_T'][:] = np.array([-50., -3,   1.])
b_param['h_T'][:] = np.array([-68., 3.75, 1.])


# reversal potentials
E["Na"] = 55.
E["K"] = -80.
E["Ca"] = 22.
E["Leak"] = -70. # as per Molineux et al 2005 and NOT the -38 in Alexander et al 2019

# model currents
currents_included["Na"] = True
currents_included["Kd"] = True
currents_included["A_mut"] = False
currents_included["A"] = True
currents_included["T"] = True
currents_included["Leak"] = True

# model conductances
g["Na"] = 3.4 * surf_area
g["Kd"] = 9.0556 * surf_area
g["A_mut"] = 0.
g["A"] = 15.0159 * surf_area
g["T"] = 0.45045 * surf_area
g["Leak"] = 0.07407 * surf_area


# save folder
folder = '../Sensitivity_Analysis/Data/Cb_stellate'
if not os.path.isdir(folder):
    os.makedirs(folder)

#%% setup for one-factor-at-a-time SA
var = np.array(['m', 'h', 'n', 'n_A', 'h_A'])
type_names = np.append(np.array(['shift' for i in range(var.shape[0])]),
                       np.array(['slope' for i in range(var.shape[0])]))
cur = np.array(['Na', 'Kd', 'A', 'Leak'])
type_names = np.append(type_names, np.array(['g' for i in range(cur.shape[0])]))
var = np.append(var, var)
var = np.append(var, cur)
alt_types = np.c_[var, type_names]
lin_array = np.arange(-10, 11, 1)
log_array = np.logspace(-1,1,21, base=2)

# %% multiprocessing
prominence = 50
desired_AUC_width = high/5

Parallel(n_jobs=8, verbose=9)(
    delayed(SA_Cb_stellate)(V_init, g, E, I_in, dt, currents_included, stim_time, stim_num, C, shift, scale, b_param,
                            slope_shift,gating, current, prominence, desired_AUC_width, folder, high, low, number_steps,
                            initial_period, sec, lin_array, log_array, alt_types, alt_ind, alt)
                    for alt_ind in range(alt_types.shape[0]) for alt in range(21))
```
