## ExtendedData1 for "Loss or Gain of Function? Effects of Ion Channel Mutations on Neuronal Firing Depend on the Cell Type": SA_RS_pyramidal.py

```
# -*- coding: utf-8 -*-
"""
Script to run sensitivity analysis for RS Pyramidal model

"""
__author__ = "Nils A. Koch"
__copyright__ = "Copyright 2022, Nils A. Koch"
__license__ = "MIT"

import numpy as np
from joblib import Parallel, delayed
import os
from Code.Functions.Utility_fxns import capacitance, stimulus_init, init_dict, NumpyEncoder
from Code.Functions.Pospischil_fxns import SA_Pospischil


# model parameters
dt = 0.01
sec = 2
low = 0
high = 0.001
number_steps = 200
initial_period = 1000
num_gating = 10
num_current = 7
C, surf_area = capacitance(61.4, 1)
stim_time, I_in, stim_num, V_m = stimulus_init(low, high, number_steps, initial_period, dt, sec)
shift, scale, slope_shift, E, currents_included, b_param, g = init_dict(
    np.array(['m', 'h', 'n', 'q', 'r', 'p', 's', 'u', 's_mut', 'u_mut']))
tau_max_p = 608
V_init = -70
V_T = -56.2

# initialize arrays
current = np.zeros((num_current, stim_num))
gating = np.zeros((num_gating, stim_num))

# gating parameters
b_param = {}
b_param['m'] = np.array([-34.33054521, -8.21450277, 1.42295686])
b_param['h'] = np.array([-34.51951036, 4.04059373, 1., 0.05])
b_param['n'] = np.array([-63.76096946, -13.83488194, 7.35347425])
b_param['q'] = np.array([-39.03684525, -5.57756176, 2.25190197])
b_param['r'] = np.array([-57.37, 20.98, 1.0])
b_param['p'] = np.array([-45., -9.9998807337, 1.])
b_param['s'] = np.array([-14.16, -10.15, 1.0])
b_param['u'] = np.array([-31., 5.256, 1., 0.245])
b_param['s_mut'] = np.array([-14.16, -10.15, 1.])
b_param['u_mut'] = np.array([-31., 5.256, 1., 0.245])


# reversal potential
E["Na"] = 50.
E["K"] = -90.
E["Ca"] = 120.
E["Leak"] = -70.3

# model currents
currents_included["Na"] = True
currents_included["Kd"] = True
currents_included["Kv"] = False
currents_included["Kv_mut"] = False
currents_included["L"] = False
currents_included["M"] = True
currents_included["Leak"] = True

# model conductances
g["Na"] = 56. * surf_area
g["Kd"] = 6. * surf_area
g["M"] = 0.075 * surf_area
g["Kv"] = 0.
g["Kv_mut"] = 0.
g["L"] = 0.
g["Leak"] = 0.0205 * surf_area

folder = '../Sensitivity_Analysis/Data/RS_pyramidal'
if not os.path.isdir(folder):
    os.makedirs(folder)

#%% setup for one-factor-at-a-time SA
var = np.array(['m', 'h', 'n'])
type_names = np.append(np.array(['shift' for i in range(var.shape[0])]),
                       np.array(['slope' for i in range(var.shape[0])]))
cur = np.array(['Na', 'Kd', 'Leak'])
type_names = np.append(type_names, np.array(['g' for i in range(cur.shape[0])]))
var = np.append(var, var)
var = np.append(var, cur)
alt_types = np.c_[var, type_names]
lin_array = np.arange(-10, 11, 1)
log_array = np.logspace(-1,1,21, base=2)

# %% multiprocessing
prominence = 50
desired_AUC_width = high/5

Parallel(n_jobs=8, verbose=9)(
    delayed(SA_Pospischil)(V_init, V_T, g, E, I_in, dt, currents_included, stim_time, stim_num, C, tau_max_p, shift,
                           scale,
                           b_param, slope_shift, gating, current, prominence, desired_AUC_width, folder, high, low,
                           number_steps, initial_period, sec, lin_array, log_array, alt_types, alt_ind, alt)
    for alt_ind in range(alt_types.shape[0]) for alt in range(21))
```
